# Systematic identification and characterization of novel genes in the regulation and biogenesis of photosynthetic machinery

**DOI:** 10.1101/2022.11.12.515357

**Authors:** Moshe Kafri, Weronika Patena, Lance Martin, Lianyong Wang, Gillian Gomer, Arthur K Sirkejyan, Audrey Goh, Alexandra T. Wilson, Sophia E Gavrilenko, Michal Breker, Asael Roichman, Claire D. McWhite, Joshua D. Rabinowitz, Frederick R Cross, Martin Wühr, Martin C. Jonikas

## Abstract

Photosynthesis is central to food production and the Earth’s biogeochemistry, yet the molecular basis for its regulation remains poorly understood. Here, using high-throughput genetics in the model eukaryotic alga *Chlamydomonas reinhardtii*, we identify with high confidence (FDR<0.11) 70 previously-uncharacterized genes required for photosynthesis. We then provide a resource of mutant proteomes that enables functional characterization of these novel genes by revealing their relationship to known genes. The data allow assignment of 34 novel genes to the biogenesis or regulation of one or more specific photosynthetic complexes. Additional analysis uncovers at least seven novel critical regulatory proteins, including five Photosystem I mRNA maturation factors and two master regulators: MTF1, which impacts chloroplast gene expression directly; and PMR1, which impacts expression via nuclear-expressed factors. Our work provides a rich resource identifying novel regulatory and functional genes and placing them into pathways, thereby opening the door to a system-level understanding of photosynthesis.

**Highlights:**

- High-confidence identification of 70 previously-uncharacterized genes required for photosynthesis
- Proteomic analysis of mutants allows assignment of function to novel genes
- Characterization of 5 novel Photosystem I mRNA maturation factors validates this resource
- MTF1 and PMR1 identified as master regulators of photosynthesis

## INTRODUCTION

The evolution of oxygenic photosynthesis in cyanobacteria ∼2.5 billion years ago fundamentally changed life on Earth (Lyons et al., 2014). Photosynthesis led to a rise in atmospheric oxygen level, enabling the evolution of aerobic respiration and ultimately of the eukaryotic cell (Hedges et al., 2004). One such eukaryotic cell is thought to have then engulfed a cyanobacterium, which then evolved into the organelle known as the chloroplast. In allowing the eukaryotic cell to convert light into energy, this engulfment eventually gave rise to the great diversity of photosynthetic eukaryotes present today (Yoon et al., 2004).

In photosynthetic eukaryotes, the photosynthetic apparatus consists of a series of protein complexes in the chloroplast thylakoid membrane that use light energy to produce NADPH and ATP (Blankenship, 2008). NADPH and ATP, in turn, power CO_2_ assimilation into sugar by the Calvin-Benson-Bassham metabolic cycle (Figure 1A)(Michelet et al., 2013). As a sophisticated system central to cellular fitness, hundreds of genes are required to generate these complexes and regulate their assembly and activity (Rast et al., 2015). Furthermore, photosynthetic complexes are assembled from components encoded in both the nucleus and chloroplast, which requires extensive coordination under the control of the nucleus (Goldschmidt-Clermont, 1998). In plants and green algae, this coordination is known to involve a range of different mechanisms, including posttranscriptional regulation of chloroplast-expressed genes by nuclear-encoded proteins (Choquet and Wollman, 2002), translational regulation of chloroplast-expressed subunits by assembly intermediates of photosynthetic complexes (Choquet and Wollman, 2009), and degradation of unassembled subunits by proteases (Majeran et al., 2000).

**Figure 1.**
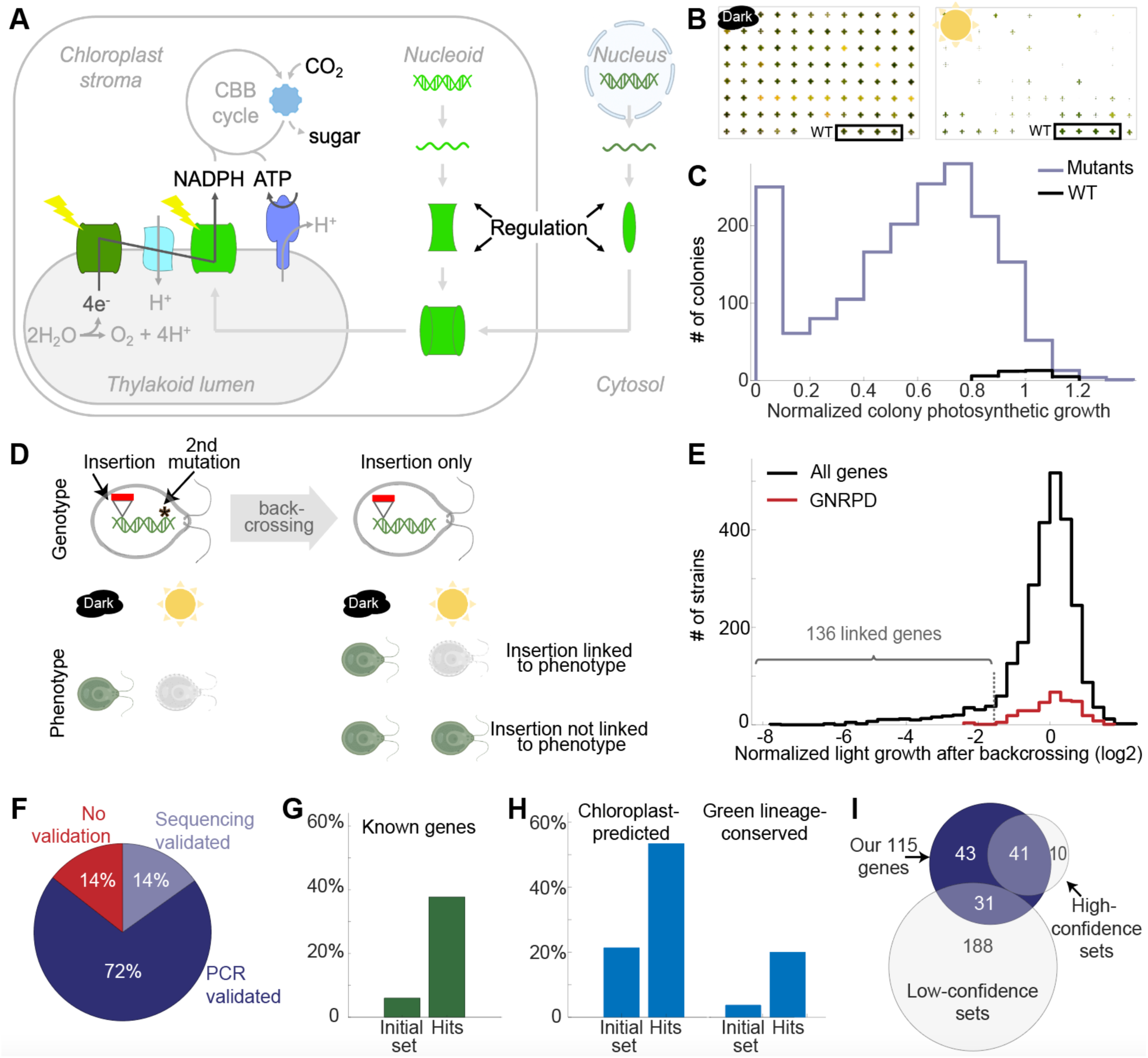
We identified 115 genes required for photosynthesis. (A) In eukaryotic photosynthesis, protein complexes in the thylakoid membranes produce ATP and NADPH to power the CO_2_-fixing Calvin-Benson-Bassham (CBB) Cycle. The complexes are assembled from subunits encoded in the chloroplast and nuclear genomes, under the control of the nucleus. (B) We validated the photosynthetic growth phenotypes of 1,781 previously identified photosynthesis-deficient *Chlamydomonas* mutants (Li et al., 2019). Photosynthesis-deficient mutants can grow in the dark with acetate supplementation, but have growth defects in light without acetate (WT, wild type). Images of a mutant plate are shown after background removal in MATLAB (original photograph in Figure S4A). (C) Colony size distribution of different mutants (blue) or WT (black). The colony size is the median of 4 replicates (2 independent experiments with two duplicates in each, see STAR Methods). Approximately 76% of the mutants showed a pronounced photosynthetic phenotype (normalized colony size <0.8). (D) Most of the mutant strains have additional (second-site) mutations, and the photosynthetic phenotype could originate from them. To evaluate whether our insertion is genetically linked to the photosynthetic phenotype, we used backcrossing to allow segregation between the insertion and the second-site mutation. For higher throughput, we developed and employed a pooled backcrossing method. For details, see Figure S2 and the STAR Methods. (E) Histogram of normalized light growth after backcrossing (log_2_ scale) for all strains (black) and for strains disrupted in Genes whose disruption likely did Not Result in a Photosynthesis Defect” (GNRPD - red). When using a threshold of -1.55, we obtained 136 candidates with an FDR < 0.11 (Figure S2). (F) We validated the insertion mapping of ∼86% of the candidates using PCR and sequencing (Figure S3, STAR Methods). (G) Approximately 39% of our hits had a previously-known role in photosynthesis (29 in *Chlamydomonas* and 16 in land plant homologs), compared to 6% in the initial set. (H) Our photosynthetic hits are enriched in chloroplast-predicted genes (by PredAlgo: Tardif et al., 2012) and green lineage-conserved genes (GreenCut2 genes: Karpowicz et al., 2011). (I) Our 115 photosynthetic hits captured most of the previously-shown high-confidence hits (41 of 51) and increased the confidence of ∼ 12% of the previously low-confidence hits (31 of 219) (see STAR Methods).

Although photosynthesis and its regulation have been extensively studied for 70 years (Bassham et al., 1950; Fromme and Mathis, 2004), phylogenetics suggests that hundreds of genes participating in photosynthesis remain to be identified and characterized. Indeed, approximately half of the GreenCut2 genes —a set of 597 genes that are conserved only in the green photosynthetic eukaryotic lineage, and are therefore likely to be involved in photosynthesis (Karpowicz et al., 2011)— have not been functionally characterized.

Genetic screens have been done in land plants and algae to identify the missing genes participating in photosynthesis. Photosynthesis-deficient mutants have been identified in land plants (primarily *Arabidopsis thaliana* and maize) by screening for leaf coloration (Wilson-Sánchez et al., 2014; Zhao et al., 2020), seedling lethality (Budziszewski et al., 2001), and chlorophyll fluorescence (Meurer et al., 1996; Shikanai et al., 1999). In addition to land plants, the leading unicellular model eukaryotic alga *Chlamydomonas reinhardtii* (Chlamydomonas) is a complementary system that provides advantages of higher throughput and physiology that facilitates the identification and characterization of genes essential to photosynthesis (Rochaix, 2002). Specifically, unlike most plants, Chlamydomonas assembles a functional photosynthetic apparatus in the dark, and mutants with defects in photosynthesis typically grow well in the dark when provided with a source of carbon and energy such as acetate (Levine, 1960). These characteristics have been leveraged for over 50 years to identify and characterize mutants with defects in photosynthesis, including in many core components of the photosynthetic electron transport chain (Gorman and Levine, 1965, 1966; Lu et al., 2020).

In the past decade, several hundred candidates for novel genes involved in photosynthesis have been uncovered by screens of two large Chlamydomonas mutant collections, Niyogi CAL (Dent et al., 2005, 2015; Wakao et al., 2021) and CLiP (Fauser et al., 2022; Li et al., 2019). However, these screens had many false positives and there are indications that fewer than half of these candidates are actually involved in photosynthesis (Li et al., 2019). Current challenges facing the field include 1) determining which of these candidates are genuinely involved in photosynthesis and 2) determining the functions of validated novel photosynthesis genes. The absence of global approaches to validation and functional characterization previously meant that such analyses were done on a slow, painstaking, gene-by-gene basis.

Here, we address these two challenges by combining genetics and proteomics to identify and functionally characterize novel genes required for photosynthesis with high confidence on a global scale. We first identified with high confidence (FDR <0.11) 70 novel and 45 previously-characterized genes required for photosynthesis by confirming linkage of each mutation with the observed photosynthetic defect and validating insertion site mappings. We then determined the proteomic profiles of mutants representing nearly all of these genes to allow their functional characterization, including assigning 34 of them to specific photosynthetic pathways. As proof of principle for the utility of our resource, we performed additional analyses to discover novel factors that advance the understanding of the regulation of the photosynthetic apparatus. We determined how five of these novel factors work with known factors to regulate the mRNA maturation of key Photosystem I subunit PsaA. We also discovered and characterized two posttranscriptional master regulators of photosynthetic apparatus biogenesis, providing insights into how cells leverage chloroplast translational machinery and the regulation of nuclear gene expression to control photosynthetic complex abundance. Together, our dataset opens the door to rapid characterization of novel photosynthesis genes and provides systems-level insights into photosynthesis regulation.

## RESULTS

### A framework for high-confidence identification of genes with roles in photosynthesis

The biggest limit to confidence in previous large-scale Chlamydomonas screens for genes with roles in photosynthesis was that most mutant strains carried disruptions in multiple genes (Li et al., 2019; Wakao et al., 2021). Thus, while a photosynthetic defect could be observed in a mutant, the defect could not be connected with high confidence to a single gene unless many independent mutants in the same gene showed the photosynthetic defect (Fauser et al., 2022; Li et al., 2019).

In the first portion of this work, we overcame this challenge by developing and adapting genetic tools to dramatically improve confidence in the genes responsible for photosynthetic defects of mutants. Specifically, we developed a high-throughput implementation of traditional genetic linkage analysis between a mutation and an observed photosynthetic defect, which allowed us to identify the specific mutation likely responsible for the defect.

### Pooled backcrossing and mapping validation of 70 novel genes in photosynthesis

We started with a set of 1,781 mutants from the CLiP library of Chlamydomonas mutants that we previously identified to have a photosynthetic growth defect (Li et al., 2019). We individually validated each strain’s photosynthetic phenotype using an automated spot test on agar (Figure 1B–1C and STAR Methods).

These mutants were generated by the random insertion of a DNA cassette and insertion sites were mapped by high-throughput sequencing (Li et al., 2019). In addition to our mapped insertion, most of our strains carry one or several additional mutations. We needed to determine if the photosynthetic defect was caused by the mapped insertion or by another unknown mutation. To determine if a given mapped insertion was the likely cause of the observed photosynthetic defect, we determined if it was genetically linked to the defect using backcrossing.

Backcrossing involves mating a mutant of interest with a wild-type strain and analyzing the progeny. This process results in random segregation of the different mutations present in the original mutant strain, thereby allowing the separation of the impact of each mutation on the phenotype of interest — in our case, defective photosynthetic growth. If all progeny carrying a particular insertion exhibit a defect in photosynthetic growth, we conclude that the insertion is genetically linked to the defect, indicating that disruption of the gene likely caused the defect (Figure 1D).

To overcome the limited throughput of traditional backcrossing of only ∼10 mutants per experiment, we developed a pooled backcrossing method that allowed us to backcross nearly 1,000 mutants in each experiment (Figure S2A, STAR Methods, and Breker et al., 2018). We backcrossed pools of hundreds of mutants and then grew the pooled progeny under photosynthetic and heterotrophic conditions. We determined the relative abundance of each insertion after growth under each condition by sequencing the unique DNA barcode(s) associated with that insertion (Figure 1E, Table S1, STAR Methods, and Li et al., 2019). If a certain barcode was depleted in the photosynthetic condition pool, we considered the corresponding insertion linked to the photosynthesis defect and concluded that the disrupted gene is likely required for photosynthesis.

We sought to estimate the frequency of incorrect identification of causal genes in this approach. Such errors could arise in rare cases where the insertion is not causal but merely in the genomic vicinity of the causal mutation. We quantified the frequency of such cases with a false discovery rate (FDR) metric. To calculate the FDR, we used a set of genes whose disruption likely did not result in a photosynthesis defect, and measured their prevalence among our hits (Figure 1E and S2B-S2D). This calculation identified 227 genes linked to a photosynthetic defect with an FDR of 0.3. Using a stricter threshold, we identified 136 genes with an FDR of 0.1 (Figures 1E, S2C-S2D and Table S2); we continued with this set for further analysis. 27 of these 136 genes were represented by two or more independent linked insertions, providing further support of their roles in photosynthesis.

It is known that some of the insertions from the starting collection of 1,781 mutants are mapped to incorrect sites in the genome (Li et al., 2019). Therefore, we validated the mapping of our linked insertions. We first checked for expected insertion sites using colony PCR (Figure S3). In cases where this failed, we used whole-genome sequencing to validate insertions or identify the actual insertion site (Figures 1F, S1, S3, Table S2, and STAR Methods). Altogether, we identified with high fidelity 115 genes required for photosynthesis from our initial set of ∼1,800 photosynthesis-deficient mutants (Figure S1 and Table S2).

Approximately 40% of the 115 genes have a known role in photosynthesis in Chlamydomonas (29 genes) or in land plants (16 genes) (Figure 1G and Table S2), a substantial enrichment compared to ∼6% of the genes in the initial ∼1,800 mutants. The 115 genes are also enriched in metrics associated with photosynthesis: they show a 2.5-fold enrichment in predicted localization to the chloroplast (Predalgo - Tardif et al., 2012) and a 4-fold enrichment in genes conserved specifically in the green lineage (Karpowicz et al., 2011) (Figure 1H).

A subset of our data provides orthogonal validation of previously-identified candidate photosynthesis genes. Our 115 genes required for photosynthesis include 41 of the 51 genes identified with high-confidence (FDR<0.05 and FDR<0.3) in previous large-scale photosynthesis screens based on the CLiP mutant collection (Fauser et al., 2022; Li et al., 2019) (Figure 1I). This high overlap shows the quality of both datasets. Our 115 genes additionally include 31/219 genes that were previously low-confidence candidates (no FDR was calculated) in the CLiP and Niyogi CAL collections (Figure 1I), increasing the confidence that these 31 uncharacterized genes do indeed participate in photosynthesis. Of the remaining 43 genes, 38 had not previously been identified as being required for photosynthesis in any organism.

Altogether, our 115 genes included 70 novel genes whose molecular function in photosynthesis had not been previously characterized in any organism. Given the novelty of these genes, we have noted in Table S2 additional information from other sources that further supports or weakens our confidence in their involvement in photosynthesis. The study of these novel genes represents a new frontier for photosynthesis research.

### Hit validation and protein localization demonstrates the value of our gene list

To experimentally validate the involvement of our novel genes in photosynthesis, we sought to genetically rescue the photosynthetic defect of mutants with insertions in novel genes. Gene rescue involves testing whether transforming a mutant with a wildtype copy of the gene alleviates the phenotype (Figure 2A). Gene rescue is notoriously challenging in Chlamydomonas due to difficulties with PCR amplification and expression of heterologous genes (Mackinder et al., 2017; Neupert et al., 2020; Zhang et al., 2014). Despite these challenges, we managed to rescue mutants in 16 genes out of the 36 genes for which a transformation was attempted. This success rate is close to the maximum that would be expected even if all 36 genes were required for photosynthesis, considering that only 30-50% of transformed constructs express in medium-throughput efforts of this nature in Chlamydomonas (Mackinder et al., 2017; Wang et al., 2022). The photosynthesis genes validated by mutant rescue included 12 genes that had not previously been implicated in photosynthesis in any organism (Table 1, Figure 2, 6B and 6J) and two photosynthesis genes that had not previously been characterized in Chlamydomonas (Figure 2 and Table S3).

**Figure 2.**
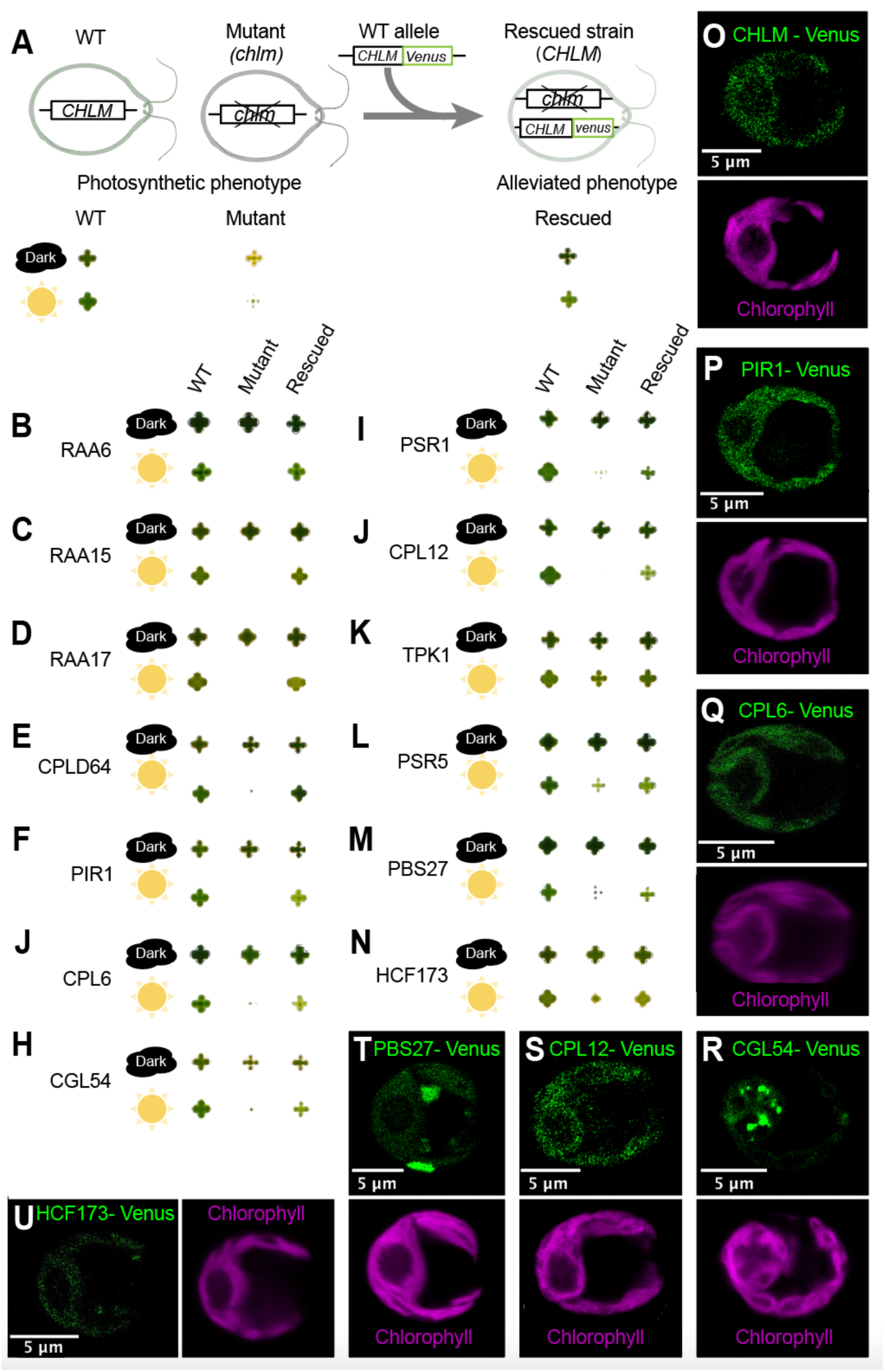
We rescued novel genes and localized their protein products. (A) Illustration of the genetic rescue procedure for the known chlorophyll biosynthesis gene *CHLM*. In the dark with acetate, the *chlm* mutant grows almost as well as wild type but is yellow (Meinecke et al., 2010); under high light, the mutant has a severe growth defect. Transformation of the mutant with a Venus-tagged *CHLM* alleviates both the color and growth phenotypes. (B - N) The colony growth of wild type, mutants, and the mutants we rescued by transforming the wild-type genes. The images were edited for background removal using a MATLAB script (see Figure S4B for the original images). (O) Localization of CHLM-Venus in the wild-type background. A similar localization was observed in the rescued strain. (P-U) localizations of Venus-tagged proteins are shown. CPL6, CGL54, and HCF173 are in the mutant background; PIR1, CPL12, and PBS27 are in the wild-type background due to insufficient expression in the rescued mutant strain.

**Table 1:**
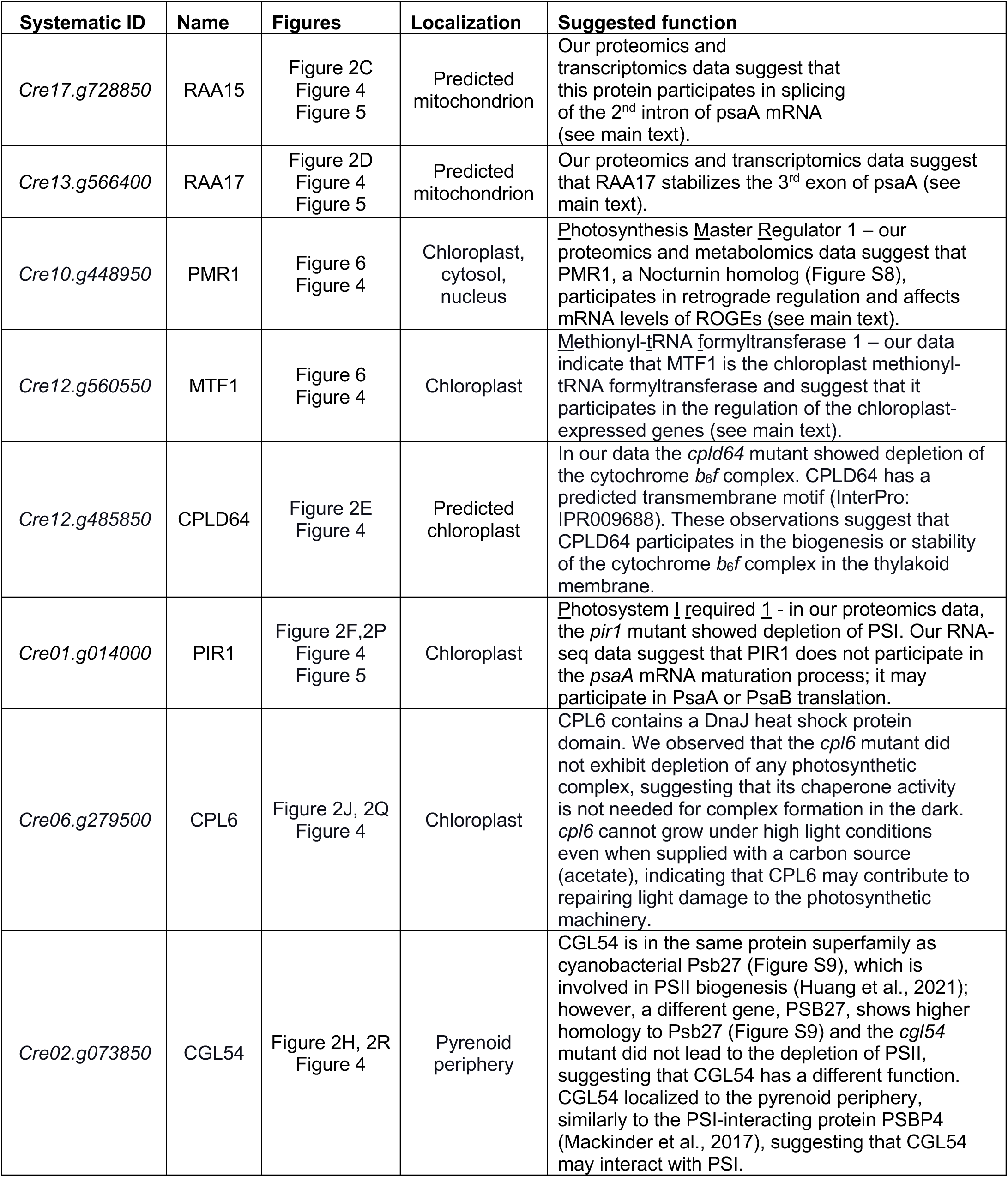
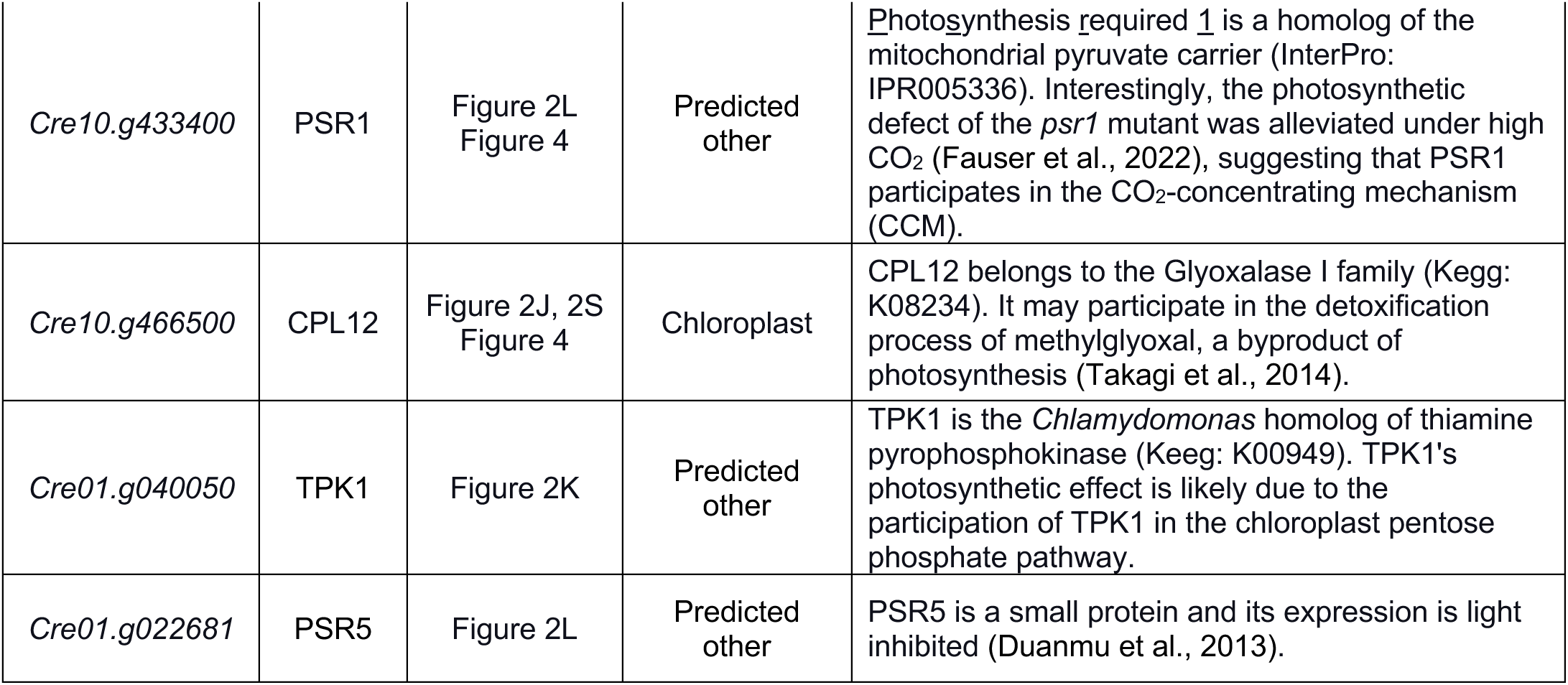
Protein localizations and suggested functions of the rescued novel genes.

Our constructs used for the rescue attempts included a C-terminal fluorescent Venus tag, which allowed us to attempt to experimentally determine protein localizations. Nine of the sixteen proteins showed sufficient expression to allow determination of their localization (Figure 2O-2U, 6F and 6R). While two of the proteins exhibited dual localizations (Figure 2T and 6R), in every case a significant portion of the protein localized to the chloroplast, consistent with the central role of the chloroplast in photosynthesis.

Based on the literature and our data (Table 1), we suggest that of the 12 rescued novel genes, at least four are posttranscriptional regulation factors (RAA17, RAA15, PMR1, and MTF1), four are biogenesis or repair factors for the photosynthetic apparatus (CPLD64, PIR9, CPL6, and CGL54), and three play roles in metabolism (PSR1, CPL12, and TPK1). The validation of these novel genes illustrates how much remains to be learned about photosynthesis and underscores the quality and value of our high-confidence list of novel genes as a starting point for studying the lesser-known areas of photosynthesis.

### One hundred mutant proteomes inform gene functions

To expand our understanding of the 115 genes we identified as required for photosynthesis and to elucidate the specific roles of uncharacterized genes within this set, we sought to use proteome profiling (Figure 3A). Proteome profiling uses mass spectrometry to determine the impact of the loss of a specific gene on the proteome. We reasoned that this would be an informative approach to characterize mutants deficient in photosynthesis because the core activities of photosynthesis are mediated by a series of highly-expressed protein complexes whose abundance is affected by photosynthetic activity, regulation, and biogenesis. Indeed, many known photosynthesis-deficient mutants show differences in protein complex abundance (Johnson et al., 2010; Peng et al., 2006; Westrich et al., 2021). Much of the regulation of the photosynthetic apparatus is thought to occur post-transcriptionally, making protein levels a more informative readout than mRNA (Choquet and Wollman, 2002).

**Figure 3.**
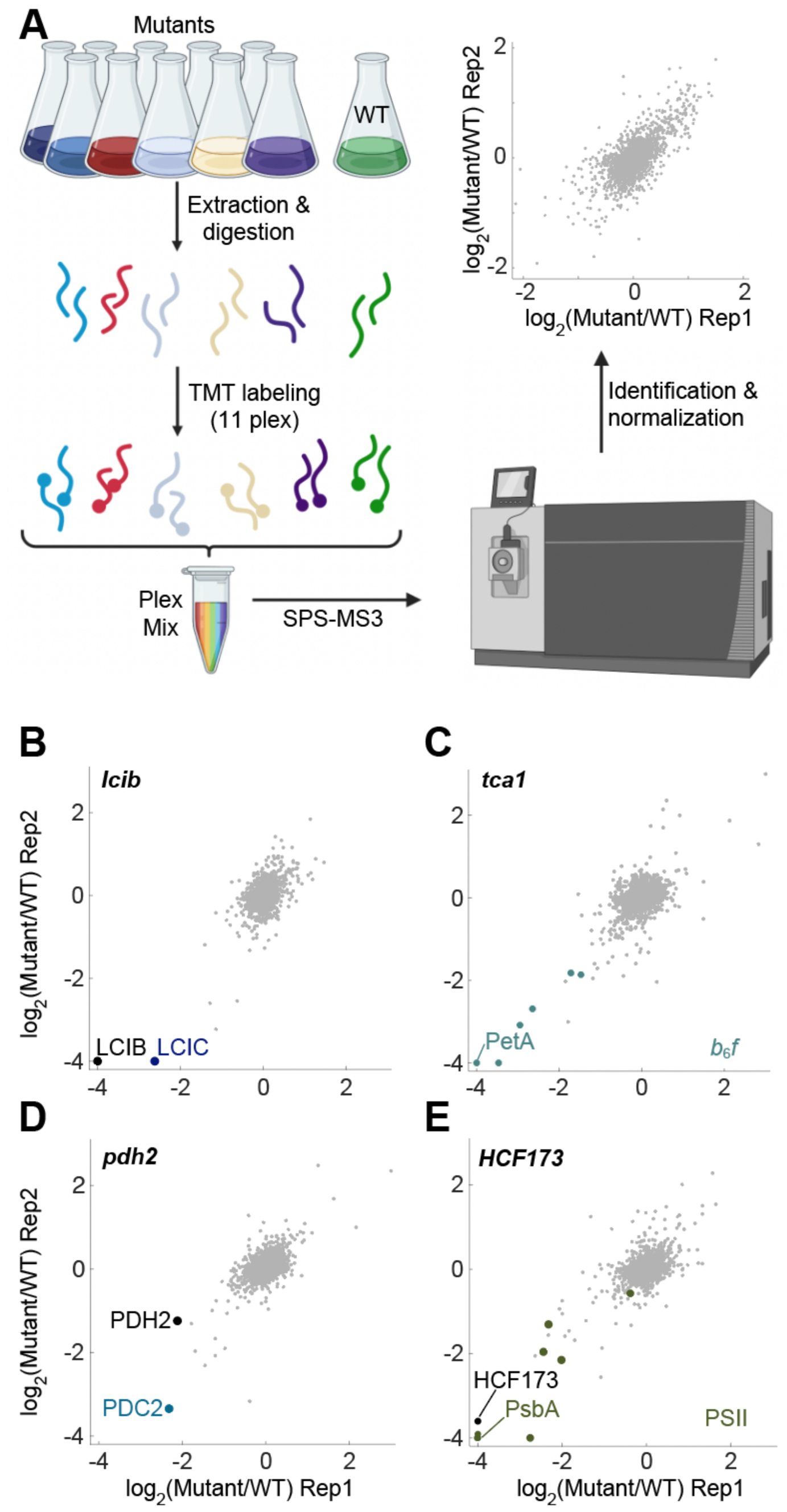
Our proteomic data reproduces known phenotypes and validate predicted phenotypes. (A) In each experiment, ten strains and a wild-type control were grown under dark conditions. After extraction and digestion, we labeled peptides with Tandem Mass Tags (TMT) and analyzed them using SPS-MS3 mass spectrometry. At least two independent experiments were carried out for each mutant (STAR Methods). (B-C) Our data set recaptures known phenotypes. (B) LCIB and LCIC are known to form a complex, and indeed, LCIC is depleted in the *lcib* mutant. (C) As expected, the *tca1* mutant leads to the depletion of the cytochrome *b*_6_*f*. Its strongest effect is on petA. (D-E) Proteome analysis revealed similarities in function between uncharacterized *Chlamydomonas* mutants and their previously characterized plant homologs. (D) Mutation in the *Chlamydomonas* homolog of pyruvate dehydrogenase E1 beta subunit PDH2 led to co-depletion of the alpha subunit PDC2. (E) Mutation in *Chlamydomonas* HCF173 (Cre13g57865), the homolog of AtHCF173, which is necessary for PsbA translation initiation in *Arabidopsis*, led to PsbA depletion together with the rest of the PSII complex. The data represent normalized log_2_ of mutant/WT protein abundance.

Our strains exhibit growth defects when grown in light, which could confound results with downstream proteomic signatures originating from slow growth or stress. To minimize such issues, we grew cells in the dark with acetate as carbon and energy source, taking advantage of the facts that under this condition Chlamydomonas photosynthesis-deficient mutant growth defects are in most cases eliminated, and wild-type cells assemble a functional photosynthetic apparatus (Rochaix, 2002). We obtained proteome profiles of mutants each disrupted for one of 100 genes (Figure S1 and Table S4), with at least two experimental repeats for each gene (Figure 3A and STAR Methods).

Our profiling dataset captured known co-depletion of proteins that form complexes and known regulatory effects. As an example of co-depletion of proteins that form a complex, the mutant lacking the carbonic anhydrase LCIB was also depleted in its known binding partner LCIC (Yamano et al., 2010, Figure 3B). As an example of regulatory effects, we observed depletion of the cytochrome *b*_6_*f* core protein petA in the mutant lacking TCA1, a known trans-acting factor required for petA translation (Wostrikoff et al., 2001). Furthermore, we see that the mutant lacking TCA1 also has lower levels of all other known *b*_6_*f* complex components (Figure 3C), as expected from previous work (Kuras and Wollman, 1994; Wostrikoff et al., 2001).

In addition to recapitulating known phenotypes, our data also illustrated that, in most cases, Chlamydomonas genes behave similarly to their characterized land plant homologs. For example, based on their homology to Arabidopsis proteins, the algal proteins PDH2 and PDC2 are predicted to be the two subunits of pyruvate dehydrogenase E1; indeed, PDH2 and PDC2 are co-depleted in the *pdc2* mutant (Figure 3D). Another example is CrHCF173, a homolog of the Arabidopsis translation initiation factor AtHCF173 that is required for PsbA translation initiation (Schult et al., 2007). As was shown for AtHCF173, we observe that mutation of CrHCF173 leads to the downregulation of psbA and the entire PSII complex (Figure 3E, Minai et al., 2006; de Vitry et al., 1989). The similar behavior of Chlamydomonas mutants compared to their land plant homologs suggests that lessons we learn in Chlamydomonas will also inform our understanding of photosynthesis across the green lineage.

### 23 novel genes impact biogenesis or regulation of individual chloroplast protein complexes

Altogether, ∼2,000 proteins were observable in most of the 100 mutant proteomes (Figure S5C and Table S5), enabling extensive opportunities for analysis. Here, we focus on the major photosynthetic protein complexes (Figure 4).

**Figure 4.**
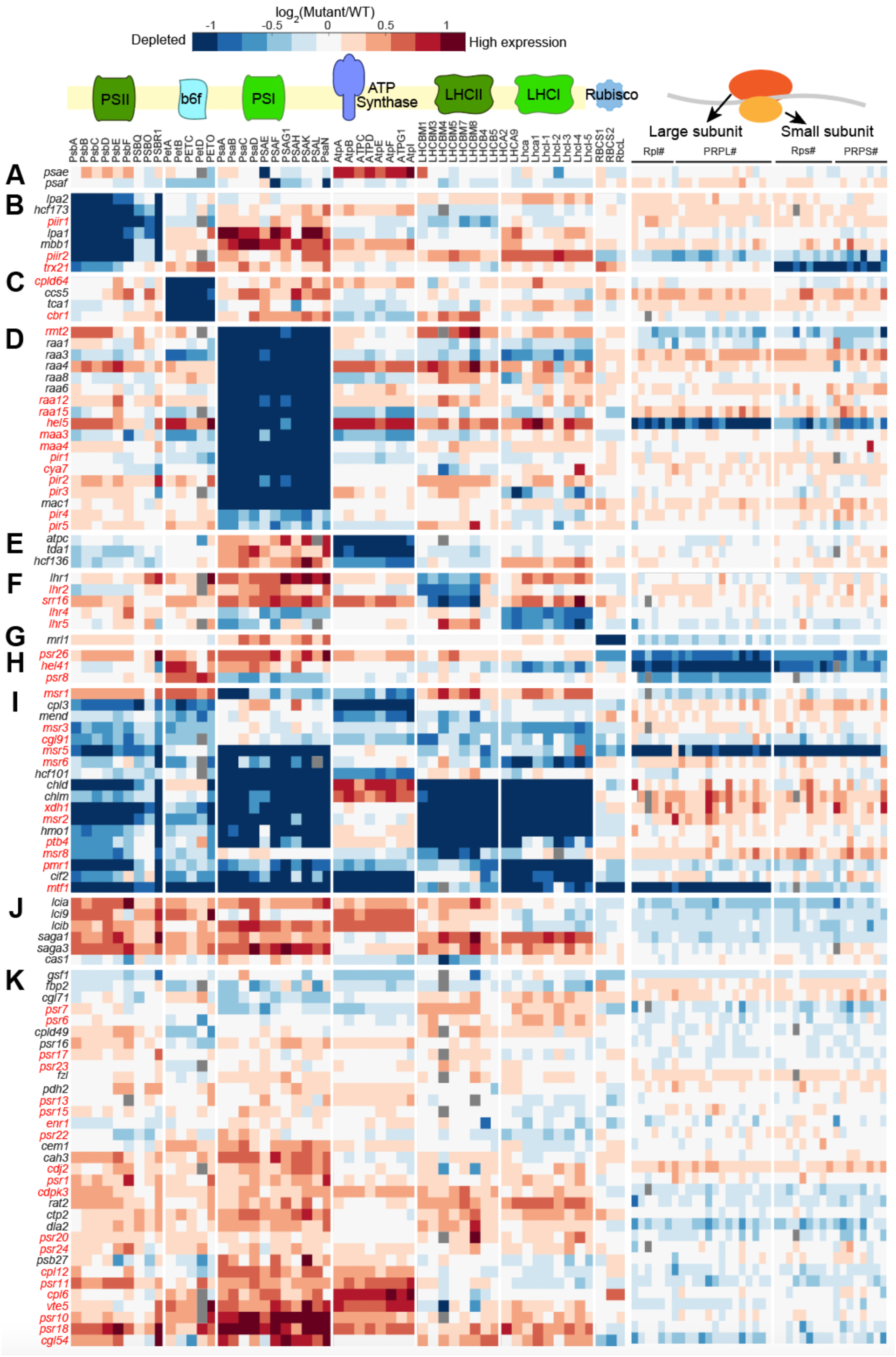
More than half of the genes we profiled are required for accumulation of one or more photosynthetic complexes. Relative abundances of photosynthetic complex and chloroplast ribosomal proteins (columns) in mutants representing 100 genes (rows). Mutants labeled in red corresponding to genes whose function in photosynthesis was not previously characterized. Each data point reflects the average normalized log_2_ (mutant/WT protein abundance) from two independent experiments (see STAR Methods). (A) Mutations in the two core Photosystem I proteins PSAE and PSAF have a local effect on Photosystem I. (B-H) Mutants were grouped according to their impact on photosynthetic complexes: (B) Photosystem II, (C) cytochrome *b*_6_*f*, (D) Photosystem I, (E) ATP synthase, (F) Light-harvesting complexes, (G) Rubisco, or (H) the chloroplast ribosomal proteins. (I) Mutations in 18 genes lead to the depletion of multiple complexes. (J) Mutant genes associated with the CO_2_ concentrating mechanism. (K) Other mutant genes.

While we observed many cases of mutants that impacted individual components of protein complexes, such as mutants that lack the PSI core subunits PSAE and PSAF (Figure 4A), a striking feature of the dataset was that more than half of our mutants showed proteomic defects in one or more entire complexes (Figure 4B-4I). Forty-one mutants led to the primary depletion of just one of the eight chloroplast protein complexes (Figure 4B-4H). These data allowed us to immediately assign 23 novel genes to a role in the biogenesis or regulation of Photosystem II, cytochrome *b*_6_*f*, Photosystem I, the light-harvesting complexes, or the chloroplast ribosome.

#### Photosystem II

Photosystem II uses light energy to extract electrons from water in the first step of the photosynthetic electron transport chain. In our dataset, mutations in seven genes led to the depletion of the entire Photosystem II complex (Figure 4B). Three of these genes were not previously associated with Photosystem II in any organism. One of these novel genes, *PIIR1 (Cre16.g658950),* encodes a protein that is predicted to localize to the chloroplast (Tardif et al., 2012) and has 6-fold higher transcript levels in light as compared to in the dark (Duanmu et al., 2013), so it may participate in the regulation of PSII in response to light. Another novel protein, TRX21 (*Cre01.g037800*), is conserved in land plants and contains a domain with thioredoxin homology. We found that mutation of TRX21 led to depletion of the chloroplast-expressed PSII subunits, suggesting that TRX21 plays a regulatory role in the biogenesis of this complex.

#### Cytochrome b_6_f

Cytochrome *b*_6_*f* pumps protons into the thylakoid lumen powered by photosynthetic electron flow. In our dataset, mutation of four genes led to the depletion of the entire cytochrome *b*_6_*f* complex (Figure 4C). Of these four genes, two novel ones, *CPLD64* (*Cre12.g485850*) and *CBR1* (*Cre12.g501550*), are conserved in land plants (Table S2) and were predicted to localize to the chloroplast (Tardif et al., 2012). We successfully used genetic rescue to validate the role in photosynthesis of CPLD64 (Figure 2E, Table 1), which contains a predicted transmembrane domain. Given their proteomic phenotypes and chloroplast localizations, we speculate that CPLD64 and CBR1 participate in the biogenesis or stability of the cytochrome *b*_6_*f* complex in the thylakoid membrane.

#### Photosystem I

Photosystem I uses light energy to energize electrons, enabling the reduction of NADP to NADPH. In our dataset, mutations in 18 genes led to the depletion of the entire Photosystem I complex (Figure 4D). Twelve of these genes were novel, including *RAA12, RAA15, RAA17-18, HEL5/CPLD46, PIR1*, and *PIR2*, which we describe in detail in later sections. Other interesting novel genes included *RMT2 (Cre12.g524500)*, and *PIR3* (*Cre01.g012200*). *RMT2* was named based on sequence homology to ribulose-1,5 bisphosphate carboxylase/oxygenase (Rubisco) large subunit N-<u>m</u>ethyltransferase (enzyme:EC:2.1.1.127), but we observed that the *rmt2* mutation did not affect Rubisco stability. Rather, it led to the depletion of Photosystem I (Figure 4D), suggesting that RMT2 actually participates in Photosystem I biogenesis. PIR3 is conserved to land plants, has a predicted basic leucine zipper (bZIP) transcription factor domain, and is predicted to localize to the cytosol or nucleus, suggesting that it regulates the transcription of nuclear-expressed Photosystem I genes.

#### Light-harvesting complexes

Light-harvesting complexes channel light excitation energy to the photosystems (Figure 4F). In our dataset, mutations in five genes affected the light-harvesting complexes — these genes include *LHR1 (Cre02.g142266)*, whose Arabidopsis homolog *CYP97A3* is known to be required for light-harvesting complex II biogenesis (Kim and DellaPenna, 2006), and four novel genes. Two of the novel genes, *LHR4 (Cre01.g016350*) and *LHR5 (Cre01.g001000*), were required for normal levels of light-harvesting complex I; and the two other novel genes, including *LHR2* (*Cre14.g616700*), affected the LHCBM proteins, the core complex of light-harvesting complex II. Interestingly, decreased levels of light-harvesting complex I in the *lhr4* and *lhr5* mutants were associated with a mild depletion of Photosystem I, with which light-harvesting complex I is associated, but the decreased levels of light-harvesting complex II proteins in the *lhr1* and *lhr2* mutants were not accompanied by a depletion of either photosystem. These results suggest that light-harvesting complex I affects the stability of Photosystem I, whereas mutants in light-harvesting complex II do not affect the stability of Photosystem II.

#### Chloroplast ribosome

Mutations in three genes, *PSR26* (*Cre50.g761497*), *HEL41* (*Cre07.g349300*), and *PSR8* (*Cre02.g110500*), led primarily to the depletion of chloroplast ribosomal proteins (Figure 4H). This depletion could be a direct or indirect effect, as ribosome abundance responds to the translational needs of the chloroplast (e.g., PRPL17–19 expression is ∼2-fold higher in light vs dark (Duanmu et al., 2013)). The helicase HEL41 was previously found to physically associate with the chloroplast ribosomal large subunit (Westrich et al., 2021) and in our dataset had a particularly strong effect on the abundance of the large subunit, suggesting that HEL41 directly affects the levels of ribosomal proteins by contributing to the biogenesis or stability of the large ribosomal subunit.

### 11 novel genes impact biogenesis or regulation of multiple photosynthetic complexes

Mutations in seven known and eleven novel genes led to the depletion of multiple complexes (Figure 4I). The known genes illustrate how the depletion of multiple complexes can result from different mechanisms. For example, cells lacking CHLD (*Cre05.g242000*) or CHLM (*Cre12.g498550*) show a depletion of both Photosystem I and II complexes (Figure 4I). CHLD and CHLM participate in chlorophyll biogenesis (Meinecke et al., 2010; Walker and Willows, 1997), so their disruption leads to the depletion of the chlorophyll-binding proteins, including subunits of Photosystems I and II, which then result in downregulation of the entire complexes (Figure 4I). Other known mutants are in regulatory genes, for example, the kinase *CPL3* (*Cre03.g185200*) (Li et al., 2019).

The novel genes affecting multiple complexes included the conserved predicted xanthine dehydrogenase/oxidase *XDH1* (*Cre12.g545101*), whose mutation led to decreased levels of Photosystems I and II and their light-harvesting complexes similarly to mutants lacking the CHLD and CHLM chlorophyll biosynthesis enzymes. These observations suggest a role for XDH1 in pigment metabolism, possibly by preventing the activation of chlorophyll degradation by xanthine (Yi et al., 2021). The novel genes also included the conserved predicted chloroplast-localized protein MSR8 (*Cre09.g400312*), which impacted both Photosystem II and light-harvesting complex II when disrupted. MSR8 contains predicted WD-40 repeats (interpro: IPR001680), which are known to serve as platforms for the assembly of protein complexes, and so we speculate that MSR8 may mediate interactions between Photosystem II and light-harvesting complex II. We also observed that the novel genes *PMR1* (*Cre10.g448950*) and *MTF1 (Cre12.g560550*) led to the depletion of the entire photosynthetic apparatus and will discuss their further characterization below.

### Characterization of novel factors that regulate photosynthetic apparatus biogenesis

Our screen and proteomics data provide a high-quality set of novel photosynthesis genes and facilitate the placement of many into specific pathways. The dataset can be used to identify functional relationships between proteins, to characterize the biogenesis of the photosynthetic apparatus, and to study protein regulation. Below, we illustrate how our data can serve as a launching point to advance our understanding of the regulation of the biogenesis of the photosynthetic apparatus.

We hypothesized that many of the novel genes encode proteins that regulate the photosynthetic machinery because many (14/24) of the known genes whose disruption led to strong depletion of the photosynthetic complexes in our proteomic experiment encode regulatory proteins (Figure 4B-4I). While different definitions for “regulatory” genes exist, for the purpose of the high-level analysis presented here, we consider a gene “regulatory” if its abundance changes under different conditions and its presence impacts the levels of one or more complexes. In our search for novel regulators of photosynthetic apparatus biogenesis, we focused on two subsets of our novel genes: ones whose disruption specifically impacted Photosystem I levels and ones whose disruption had broad effects on most or all of the photosynthetic apparatus.

### Novel components regulating Photosystem I *psaA* mRNA maturation

The mRNAs encoding chloroplast-expressed proteins are constitutively expressed, and their protein abundance is primarily regulated post-transcriptionally (Choquet and Wollman, 2002). A central mechanism for this post-transcriptional regulation involves the Regulators of Organelle Gene Expression (ROGE), nuclear-encoded factors that each promote mRNA stability/maturation (M factors) or translation (T factors) of a specific chloroplast-encoded subunit of a photosynthetic complex (Wang et al., 2015). In the absence of a T or M factor, the abundance of the regulated subunit drops, translation of other subunits decreases, and unassembled subunits are degraded, leading to depletion of the entire complex (Choquet and Wollman, 2009).

We identified six known M factors among the genes required for accumulating the entire Photosystem I complex in our proteomics (Figure 4D). One of these M factors, MAC1, is required for *psaC* mRNA stability (Douchi et al., 2016). The other five, RAA1, RAA3, RAA4, RAA6 and RAA8, participate in the maturation of *psaA* mRNA (Glanz et al., 2012; Marx et al., 2015; Merendino et al., 2006; Reifschneider et al., 2016; Rivier et al., 2001). We hypothesized that other genes with similar proteomic patterns might also be M factors. We focused on seven novel genes (*HEL5, RAA17, RAA18, RAA12, RAA15, PIR1,* and *PIR2*), of which we validated three (*RAA17*, *RAA15*, and *PIR1*) by gene rescue (Table 1), whose mutants exhibited strong and specific depletion of the Photosystem I complex (Figure 5A and S7).

**Figure 5.**
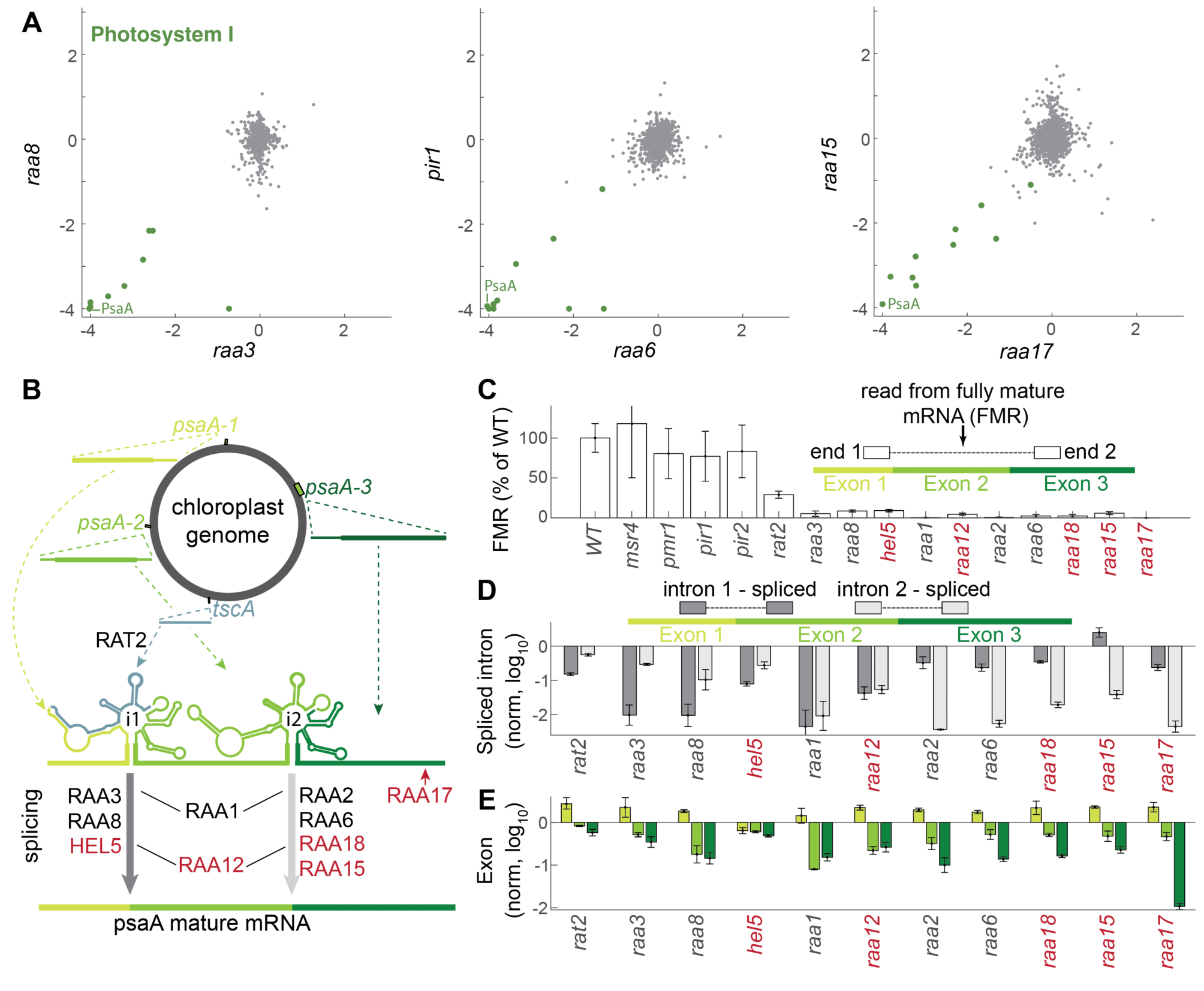
Characterization of five novel *psaA* mRNA maturation factors. (A) Scatterplots of proteomic data of mutants of known *psaA* maturation factors (RAA8, RAA3, and RAA6) and mutants in novel genes with similar proteomic profiles (PIR1, RAA15, and RAA17). For proteomic scatterplots of RAA1, RAA4, HEL5, RAA12, PIR2, and MAA18, see Figure S7. (B) *psaA* mRNA maturation process. *psaA* mRNA starts as four separated RNAs expressed in the chloroplast genome, *psaA1-3* each include an exon, and *tscA* forms part of intron 1. The RNAs hybridize to form two introns that are spliced out (gray arrows) to produce the mature mRNA. This process is mediated by M factors. The known M factors from our transcriptomic dataset are shown in black; the novel factors are shown in red. (C) Fully mature *psaA* mRNA levels were determined using paired-end reads. If the read in one end was in exon 1 and the read in the second end was in exon 3, this mRNA was considered fully matured. (D) *psaA* maturation levels in M factor mutants. The reads are normalized to wild type and shown on a log_10_ scale. When paired reads mapped to adjacent exons, the intron between them was considered spliced out. (E) Normalized reads for each exon in the indicated mutants are depicted. Error bars represent standard error of the mean.

We profiled the chloroplast transcriptome in the seven mutants of interest and reference mutants representing known factors to look for M factors among these novel genes. We did not observe defects in *psaB* or *psaC* mRNA levels in any of the mutants (Figure S7E), and neither *pir1* nor *pir2* (*Cre12.g553800*) affected levels of mature *psaA* mRNA, suggesting that PIR1 and PIR2 play roles in other aspects of Photosystem I biogenesis. However, we observed that mutations in five of the novel genes, *HEL5, RAA17, RAA18, RAA12,* and *RAA15*, result in less than 15% of the wild-type levels of mature *psaA* mRNA, similar to the mutants of known *psaA* mRNA maturation factors in our dataset (Figure 5B and 5C). These results suggest that we identified five novel *psaA* maturation factors.

PsaA is one of the two central chloroplast-encoded components of Photosystem I (Rochaix, 2002). In Chlamydomonas, its maturation involves a sophisticated mRNA splicing mechanism (Goldschmidt-Clermont et al., 1990). PsaA mRNA starts as four separate transcripts that hybridize to form a structure containing two introns, which are spliced out to generate the mature mRNA (Figure 5B). This process is mediated by a ribonucleoprotein complex that includes at least 14 splicing factors (Goldschmidt-Clermont et al., 1990). These splicing factors are classified into three groups based on their impact on the splicing of the two introns. By evaluating the relative splicing of each intron in the mutants using paired-end RNAseq, we were able to classify novel factor HEL5 as impacting intron 1, novel factors RAA15 and RAA18 as impacting intron 2, and novel factor RAA12 as impacting both introns 1 and 2 (Figure 5D). Novel factor RAA17 appears to represent a new maturation group, which we propose directly affects exon 3 stability (Figure 5B-5E).

#### HEL5 is required for splicing psaA intron 1

*HEL5* (*Cre01.g027150*) belongs to the DEAD-box helicase superfamily (Interpro: IPR011545). Its Arabidopsis reciprocal best BLAST hit ISE2 appears to be a general splicing factor that participates in the mRNA processing of chloroplast ribosome subunits, ATP synthase subunit AtpF, and protease ClpP1 (Bobik et al., 2017). While Chlamydomonas HEL5 appears to contribute to the biogenesis of the chloroplast ribosome (Figure S7D), it does not affect ATP synthase or the Clp protease. Instead, we observe that the primary function of HEL5 seems to be the splicing of *psaA* intron 1 (Figure 5C-5D and S7D), illustrating how the specificity of a splicing factor can change across evolution.

#### RAA15 and RAA18 are required for splicing psaA intron 2

Our data suggest that *RAA15* (*Cre17.g728850*) and the predicted protein kinase *RAA18* (*Cre07.g351825*) are novel genes required for photosynthesis and normal levels of Photosystem I (Figure 4D, 5A, and S7). In mutants lacking RAA15 and RAA18, we observed a 96% decrease in mature *psaA* intron 2 compared to wild type, suggesting that these genes encode intron 2 splicing factors (Figure 5D). We caution the reader that RAA18 is predicted to localize to the secretory pathway and thus we are less confident that a mutation in this gene causes the observed photosynthesis phenotypes. Transforming the wild type allele of *RAA15* into the corresponding mutant alleviated the mutant’s growth defects to almost-wildtype levels (Figure 2C), providing confidence that a mutation in this gene causes the observed photosynthesis phenotype. RAA15 was previously pulled down with known intron 2 splicing factors RAA2 and RAA7 (Lefebvre-Legendre et al., 2016; Reifschneider et al., 2016), suggesting that these three factors function together.

#### RAA12 is required for splicing psaA introns 1 and 2

RAA12 (*Cre17.g698750*) is a member of the OctotricoPeptide Repeat (OPR) family of regulatory RNA-binding proteins (Wang et al., 2015) required for photosynthesis (Table S2), whose two mutant alleles showed depletion of Photosystem I (Figure 4D and S6A). Its transcriptomic profile was similar to that of RAA1, a known M factor required for *psaA* intron 1 and 2 splicing (Merendino et al., 2006) (Figure 5D and 5E). Much like *RAA1*, we observed that *RAA12* mutation leads to the depletion of mature forms of both introns 1 and 2 (Figure 5D). Furthermore, similarly to RAA1, RAA12 was previously co-precipitated with known M factors that regulate splicing of both introns 1 and 2: known intron 1 splicing factors RAA4 and RAT2 (Jacobs et al., 2013; Reifschneider et al., 2016), and known intron 2 splicing factor RAA7 (Lefebvre-Legendre et al., 2016). These results suggest that RAA12 is required for the maturation of both introns 1 and 2.

#### RAA17 regulates psaA exon 3 stability

*RAA17* (*Cre13.g566400*) is a gene required for photosynthesis and for normal levels of Photosystem I (Figure 5A). Transforming the wild-type *RAA17* allele into the *RAA17* mutant rescues the mutant’s growth to wildtype-like levels even under high-light conditions (Figure 2D), confirming that *RAA17* is required for photosynthesis. The *RAA17* mutant exhibits almost complete depletion of exon 3 (< 2% of WT levels), a phenotype not exhibited by any of the other mutants of known factors in our dataset, suggesting that *RAA17* is a novel kind of maturation factor that specifically protects the third exon. *RAA17* is a member of the OPR family of RNA-binding proteins; thus, it is possible that it could directly bind the third exon of *psa*A and protect it. The decreased level of exon 3 is likely the cause of the decreased level of the mature form of intron 2 observed in the *raa17* mutant. *RAA17* expression is light-dependent: its expression level is 5-fold higher in light compared to dark (Duanmu et al., 2013), suggesting that it participates in *psaA* dark-to-light acclimation.

#### RAT2 is required for psaA maturation but is not a limiting factor in the dark

RAT2 is a previously known *psaA* maturation factor that participates in processing the intron 1 RNA component *tscA* ((Balczun et al., 2005), Figure 5B and S7F). As expected, a mutant strain lacking RAT2 showed photosynthetic defects in our screen, but surprisingly, it did not lead to the depletion of PSI in our protein profiling (Figure 4K). Analysis of the cells by RNA-seq provided a potential explanation for this discrepancy: the *rat2* mutant has approximately 30% of the WT mature *psa*A reads (Figure 5C-5E), which is more than two-fold more than we see in any other maturation factor mutant in our dataset. We propose that this level of mature *psa*A mRNA is sufficient for Photosystem I production in the dark, conditions under which materials were collected for our proteomic analysis, as there is a lower demand on the level of the protein complex and thereby a lower rate of translation as controlled by the T factor TAA1 (Lefebvre-Legendre et al., 2015; Young and Purton, 2014). Under light conditions requiring active photosynthesis, relatively low levels of *psaA* mRNA would not meet the higher demand for PSI production, thereby contributing to the photosynthesis defect.

#### Additional insights into psaA mRNA maturation

In addition to characterizing five novel M factors, our RNA profiling provides new insights into the overall maturation process of *psaA*. In nearly all mutants that primarily impact one intron (with *raa15* being the only exception), we observed that splicing of the other intron is also impacted (Figure 5D), suggesting that each splicing site requires integrity of the other for maximal activity. In all mutants except for *hel5*, exon 1 levels are higher than in WT, suggesting that unspliced exon 1 is more stable than the mature mRNA (Figure 5E).

Together, the above findings broaden our understanding of Photosystem I maturation and regulation and illustrate how our data can be used to rapidly functionally characterize novel factors with roles in photosynthesis.

### Identification of master regulators of photosynthesis

One of the most striking observations from our data was the identification of three genes whose mutants exhibited decreased levels of all four electron transport chain complexes without affecting the abundance of other chloroplast complexes such as the chloroplast RNA polymerase. One of these three genes, PMR1, behaves as a classical “master regulator,” as we show below that it regulates multiple nuclear-expressed factors that, in turn, each regulate one or two chloroplast genes. By contrast, the other two genes, CIF2 and MTF1, may appear at first glance to be housekeeping genes required for chloroplast translation initiation. For example, CIF2 (*Cre07.g341850*) likely functions as the chloroplast translation initiation factor 2 (IF2), which attaches the fMet-tRNA to the translation initiation complex, based on its homology to the characterized Arabidopsis IF2*, FUG1* (Miura et al., 2007), and CIF2’s physical interaction with the Chlamydomonas chloroplast ribosome (Westrich et al., 2021). However, as we show below, CIF2 and MTF1 may also play regulatory roles.

#### MTF1 is the chloroplast’s methionyl-tRNA formyltransferase (MTF) and is required for translation of nearly all chloroplast-encoded proteins

*MTF1 (Cre12.g560550)* is a conserved gene whose mutant shows a severe photosynthetic phenotype. In our proteomic experiments, loss of *MTF1* expression had the strongest phenotype — the disruption of this gene resulted in the depletion of the entire photosynthetic apparatus and nearly all chloroplast-expressed proteins (Figure 4I and 6A). We validated this phenotype by genetic rescue, which alleviated the observed growth defect in the mutant to nearly WT growth under high light conditions (Figure 6B), and recovered WT levels of chloroplast-expressed proteins (Figure 6C).

**Figure 6.**
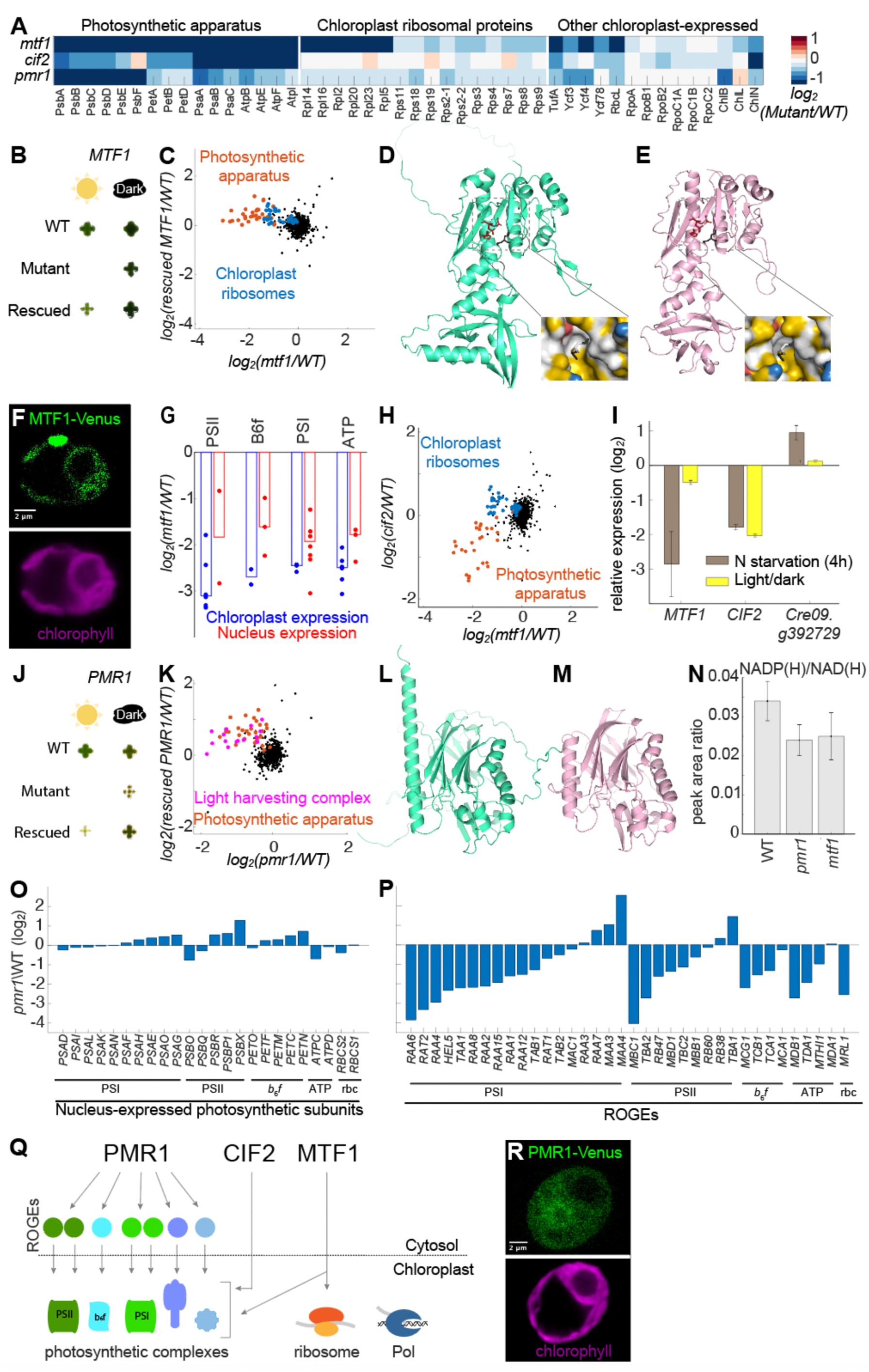
Novel master regulators of photosynthetic complexes. (A) Protein levels of chloroplast-expressed genes in *mtf1, cif2*, and *pmr1* mutants. The data represent the mean of two independent experiments. (B) Colony growth is shown for wild type (WT), the *mtf1* mutant, and the *mtf1* mutant rescued by transforming the wild-type allele under the same conditions as Figure 2A. The background of the images was removed using a MATLAB script (see Figure S4B for the original images). (C) Rescuing the *mtf1* mutant reverses its proteomic phenotype. The data represent the mean of four biological replicates for the *mtf1* mutant and two biological replicates for the rescued mutant. (D-E) Comparison between the AlphaFold-predicted MTF1 structure (D) and the crystal structure of *E. coli* MTF (E) (Schmitt et al., 1998). The conserved active site residues (Asn108, His110, and Asp146, (Schmitt et al., 1996); Asn160, His162, and Asp198 in MTF1) are shown in red and fMet in black. For a better comparison of the active sites, we used YRB (Hagemans et al., 2015), a script that displays the hydrophobic pockets (yellow) and negative charges (red) on a protein surface. In both active sites, we can see hydrophobic pockets below the fMet (to stabilize it) and a negative charge above it from the active site residues. (F) Localization of Venus-tagged MTF1 (green) and chlorophyll autofluorescence (magenta). (G) Comparison of chloroplast-expressed protein levels (blue) to nucleus-expressed protein levels (red) for photosynthetic complexes in the *mtf1* mutant. Each dot represents a protein, and the bar represents the median. (H) Comparison of *mtf1* and *cif2* proteomic data. The data represent the mean of four biological replicates. (I) Expression data (Boyle et al., 2012; Duanmu et al., 2013) for *MTF1*, *CIF2*, and *Cre09.g392729* (encoding the MTF1 ortholog, which is predicted localize to the mitochondria). (J) Colony growth is shown for wild type, *pmr1* mutant, and rescued *pmr1* mutant, as described in panel B. (K) Rescuing the *pmr1* mutant reverses its proteomic phenotype. The data represent the mean of four biological replicates for *pmr1* mutant and two biological replicates for the rescued mutant. (L-M) Comparison between the AlphaFold-predicted PMR1 structure (L) and the crystal structure of human Nocturnin (M) (Abshire et al., 2018). (N) NADP(H) and NAD(H) levels were measured (using LC-MS) in wild type, *mtf1*, and *pmr1*. The data represent three biological replicates ± SE. (O) mRNA levels of nucleus-expressed photosynthetic subunits in *pmr1* relative to wild type. (P) mRNA levels of Regulators of Organelle Gene Expression (ROGEs) in *pmr1* relative to wild type. The data in (P-Q) represent two biological replicates. (Q) Master regulator model. PMR1 regulates mRNA levels of Regulators of Organelle Gene Expression (ROGEs), which are necessary for biogenesis of the chloroplast-expressed subunits of the photosynthetic machinery. CIF2 and MTF1 directly affect the translation of different groups of chloroplast-expressed genes: CIF2 primarily affects photosynthesis genes, whereas MTF1 also affects the ribosomal proteins. (R) Localization of PMR1-Venus (green) and chlorophyll autofluorescence (magenta).

MTF1 was previously annotated as a putative methionyl-tRNA formyltransferase (MTF) based on sequence similarity to known enzymes. Methionyl-tRNA formyltransferases generate fMet-tRNA, which is the tRNA needed for translation initiation in bacteria (Kaledhonkar et al., 2019). In contrast to bacteria, eukaryotes do not use fMet-tRNA for cytosolic translation, but the chloroplast and mitochondria within eukaryotic cells require this tRNA for translation initiation. Indeed, we found that MTF1 has a similar AlphaFold-predicted structure to the known *E. coli* enzyme MTF, with the active site key residues and hydrophobic pocket conserved (Figure 6D-6E) (Jumper et al., 2021)(Schmitt et al., 1998). These similarities validate the annotation of MTF1 as a methionyl-tRNA formyltransferase.

In theory, MTF1 could provide fMet-tRNA for the chloroplast or the mitochondria. We found that Venus-tagged MTF1 localized exclusively to the chloroplast (Figure 6F). The strong effect of *mtf1* mutants on chloroplast-expressed proteins also suggest that it is active in the chloroplast. Consistent with the idea that MTF1 primarily affects chloroplast-encoded photosynthetic subunits, we observed that in the *mtf1* mutant, chloroplast-expressed subunits tended to be more depleted than their nuclear-expressed counterparts (Figure 6G), suggesting that the depletion of the nuclear-expressed subunits was a secondary effect due to degradation of incompletely assembled complexes. Together, our results strongly suggest that MTF1 is the methionyl-tRNA formyltransferase that mediates chloroplast translation initiation.

#### Factors required for chloroplast translation initiation mediate differential photosynthetic complex regulation

Traditionally, core machinery components such as CIF2 and MTF1 would be considered housekeeping genes that are required for translation but do not play a regulatory role. However, two lines of evidence suggest that CIF2 and MTF1 act as photosynthetic master regulators: first, they each are required for production of a different subset of chloroplast-expressed proteins; and second, their expression is not constitutive and reflects the cell’s regulatory needs.

If MTF1 and CIF2 were simply constitutive parts of the core translation machinery as their *E. coli* homologs are assumed to be (Marzi et al., 2003), we would have expected that MTF1 and CIF2 would be required for translation of all chloroplast-expressed proteins. Surprisingly, we found that MTF1 and CIF2 were not required for normal levels of several chloroplast-expressed proteins. For example, *mtf1* and *cif2* mutations did not affect levels of the two chloroplast-expressed proteins required for chlorophyll biosynthesis in the dark, chlB and chlL (Figure 6A). Consistent with this observation, *mtf1* and *cif2* mutants were green when grown in the dark (Figure S8A), whereas strains without the chlB/L/N complex are yellow in the dark (Cahoon and Timko, 2000). *mtf1* and *cif2* mutants also did not show downregulation of chloroplast-expressed RNA polymerase (Figure 6A, *Rpo* genes). Notably, MTF1 and CIF2 affected different subsets of genes: CIF2 was only required for the photosynthetic machinery (less than half of all chloroplast-expressed proteins), whereas MTF1 also affected ribosomal large subunits (Figure 6H). These observations indicate that MTF1 and CIF2 promote the translation of specific subsets of proteins, a property associated with regulatory factors (Macedo-Osorio et al., 2021).

Further consistent with regulatory roles, MTF1 and CIF2 are themselves differentially regulated. MTF1 is downregulated under nitrogen starvation but not in the dark, whereas CIF2 is downregulated under both nitrogen starvation and in the dark (Figure 6I. Data from: Boyle et al., 2012; Duanmu et al., 2013). These different expression patterns may reflect differential regulatory needs: under nitrogen starvation, downregulating MTF1 and CIF2 allows the cell to downregulate most chloroplast translation to conserve nitrogen. In contrast, in the dark, downregulating only CIF2 allows the cell to downregulate the photosynthesis machinery but not the ribosome, retaining translation capacity for non-photosynthetic functions.

Together, these observations suggest that CIF2 and MTF1 participate in the regulation of chloroplast-expressed genes and illustrate how translation initiation machinery can be leveraged to co-regulate multiple protein complexes.

#### CCR4-NOT family member PMR1 regulates photosynthesis through ROGE mRNA levels

Parallel to our discovery that CIF2 and MTF1 likely regulate the chloroplast translation of multiple photosynthetic complexes, we also identified the novel protein PMR1 as a master regulator of multiple photosynthetic complexes acting at the level of nuclear gene expression control. The mutant deficient in *PMR1* (*Cre10.g448950*) showed severe photosynthetic growth deficiency and depletion of all electron transport chain components, most significantly Photosystems I and II, and light-harvesting complex I (Figures 4I, 6A, and S8B). These defects were all rescued by transforming the mutant strain with the wild-type allele (Figure 6J-K).

PMR1 is a member of the CCR4-NOT family and shows the highest sequence homology (Figure S9C) and a similar predicted structure (Figure 6L-M) to Nocturnin (KEGG K18764), a protein that has been identified as a circadian-controlled master regulator that affects metabolism and hundreds of transcripts in animals (Abshire et al., 2020; Green et al., 2007; Kawai et al., 2010). Consistent with Nocturnin-like characteristics, we observed that PMR1 has periodic expression (Figure S8C, data from Strenkert et al., 2019), and the disruption of its expression affects protein levels of most of the photosynthetic complexes (Figure 6A, Figure S8B) and influences the levels of thousands of mRNAs (Figure S8D).

Members of the CCR4-NOT family regulate mRNA post-transcriptionally (Miller and Reese, 2012). Nocturnin was originally proposed to directly regulate mRNAs (Abshire et al., 2018) by affecting stability (Baggs and Green, 2003) and/or export from the nucleus (Kawai et al., 2010). Moreover, its active site is very similar to known deadenylases (CNOT6L and PDE12), which directly regulate mRNA stability. Furthermore, a recent paper showed that human and fly Nocturnin act as phosphatases that convert NADP(H) to NAD(H) (Estrella et al., 2019), which then has secondary effects on the transcriptome.

In order to determine if PMR1 acts as a NADP(H) phosphatase similar to the human Nocturnin, we metabolically analyzed the *pmr1* mutant. If PMR1 is NADP(H) phosphatase, we would expect the mutant to show an increase in the ratio of NADP(H) to NAD(H). Instead, we observed that the *pmr1* mutant showed a slight decrease in this ratio (Figure 6N). This effect was likely nonspecific, as the *mtf1* mutant showed a similar decrease (Figure 6N). These results suggest that PMR1 does not act as NADP(H) phosphatase *in vivo* and more likely regulates mRNA levels directly, similarly to other members of the CCR4-NOT family (Mittal et al., 2011).

Our RNA-seq analysis suggests that PMR1 regulates the levels of photosynthetic complexes through broad control of the Regulators of Organelle Gene Expression (ROGE), nuclear-encoded factors that each regulate the mRNA stability or translation of one or two chloroplast-expressed genes (Wang et al., 2015). The *pmr1* mutant did not show significant depletion of nuclear-encoded subunits of photosynthetic complexes (Figure 6O). Instead, the *pmr1* mutant exhibited strong depletion of 22 ROGEs that together regulate all major photosynthetic complexes, most notably ROGEs required for biogenesis of Photosystems I and II (Figure 6P; u= 0.01, Wilcoxon rank sum test comparing the ROGE mRNA distribution to the distribution of all measured mRNAs). Since the depletion of even one ROGE can lead to the depletion of an entire photosynthetic complex, we propose that this downregulation of ROGEs explains the observed broad downregulation of all photosynthetic complexes in the *pmr1* mutant (Figure 6Q).

If PMR1 directly regulates the mRNA of nuclear-expressed genes, we would expect it to localize to the cytosol and/or nucleus. Consistent with this idea, fluorescently-tagged PMR1 localized to the cytosol and nuclear periphery (Figure 6R). Intriguingly, a substantial fraction of the protein also localizes to the chloroplast. This additional site of localization suggests the possibility that PMR1 participates in retrograde regulation — signaling from the chloroplast to the nucleus and cytosol to regulate nuclear-expressed genes (Chan et al., 2016).

## DISCUSSION

Even though photosynthesis is central to life on Earth, many of the genes required for it remain uncharacterized or even unknown. In this study, we identified with high confidence (FDR < 0.11) 115 genes required for photosynthesis, including 70 whose functions in photosynthesis had not been characterized in any organism. Our confidence in the identification of these genes is supported by a statistical framework as well as gene rescue of mutants representing 12 novel genes.

We then showed that mutant proteomes provide key insights into the functions of these genes in photosynthesis, in many cases allowing the assignment of novel genes to specific pathways. The quality of the proteomic data is supported by the recapitulation of many known phenotypes, and the specificity of protein depletion has guided our follow-up studies reported here. While we focused on mutants that entirely lacked core photosynthetic complexes, we note that the proteomes are also useful for mutants where core complexes were not depleted. For example, although mutations in CO_2_-concentrating mechanism-associated genes such as LCIB (Wang and Spalding, 2006), SAGA1 (Itakura et al., 2019), SAGA3 (Fauser et al., 2022), CAS1 (Wang et al., 2016), and LCI9 (Mackinder et al., 2017) do not affect the core photosynthesis complexes, each mutant proteome shows a distinctive pattern that may aid in the understanding of their contribution to the biogenesis and regulation of the CO_2_-concentrating mechanism.

We illustrated here the value of this resource by employing transcriptomics, protein localization, and metabolomics to further characterize seven novel genes, yielding insights that contribute to the basic understanding of photosynthesis regulation.

### Master regulators MTF1, CIF1 and PMR1 coordinate photosynthetic complex expression

To date, many individual nuclear-encoded factors have been identified that each regulate one or two chloroplast-encoded proteins post-transcriptionally (Choquet and Wollman, 2002). However, in order to respond effectively to changing conditions, the cell must simultaneously regulate multiple photosynthetic complexes. Our results suggest that MTF1, CIF2, and PMR1 are master regulators that contribute to these responses by coregulating multiple complexes.

MTF1 and CIF2 are part of the chloroplast translation machinery, and our data suggest that the cell can use variations on this machinery to differentially regulate multiple complexes. The observation that not all chloroplast-expressed proteins are dependent on MTF1 and CIF2 also raises intriguing questions for future studies about how the remaining proteins are translated. For example, the *E. coli* CIF2 homolog IF2 is thought to be essential for initiation of all translation (Madison et al., 2012), and it remains unclear how translation of chloroplast ribosomal proteins in Chlamydomonas is initiated in the absence of CIF2.

Our data suggest that PMR1 is a master regulator that operates on a different principle: it regulates the mRNA levels of 22 nuclear-encoded Regulators of Organelle Gene Expression, which then each regulate the mRNA stability or translation of one or two chloroplast-encoded subunits of photosynthetic complexes (Figure 6Q). Furthermore, its multiple localization to the chloroplast, cytosol, and nucleus suggests that PMR1 might participate in retrograde regulation, where it could sense signals in the chloroplast that regulate its activity in the cytosol and nucleus.

### Our data support a regulatory role for ROGEs

We identified five novel Regulators of Organelle Gene Expression (ROGEs) that are essential for the biogenesis of Photosystem I. Including these novel genes, 75% (16/21) of genes with known functions in our dataset that lead to the depletion of an entire complex are ROGEs (Figure 4), demonstrating their significant impact on the biogenesis of photosynthetic complexes.

It is currently debated whether ROGEs play a regulatory role or are merely required for complex biogenesis (Wang et al., 2015). Existing evidence supporting a regulatory role includes 1) Different ROGEs affect different chloroplast-encoded genes (Choquet and Wollman, 2002); 2) ROGEs are differentially transcriptionally regulated (Lefebvre-Legendre et al., 2015); 3) several ROGEs can co-regulate the same protein (Table S6, (Boulouis et al., 2011; Lefebvre-Legendre et al., 2016)); 4) proteins with a stronger effect on growth, including the largest subunit of each complex, tend to be regulated by more ROGEs (Table S6); and 5) ROGEs participate in feedback loops (Boulouis et al., 2011; Choquet and Wollman, 2009), a classical transcription network motif (Milo et al., 2002).

Our data now add two new observations that further support a regulatory role for ROGEs. The first is that different ROGEs can be limiting factors in different conditions, e.g., RAT2 is not a limiting factor for *psaA* expression in the dark but is in the light (Figure 4K and 5C-5E). The second is that a master regulator, PMR1, appears to use ROGEs to coregulate the abundance of multiple complexes.

Together, these points raise the intriguing possibility that during the endosymbiosis process, as transcriptional regulation in the chloroplast was lost (Choquet and Wollman, 2002) the ROGEs evolved to generate a regulatory network quantitatively regulating chloroplast expressed proteins, in a condition-dependent manner, under the control of master regulators.

Interestingly, while ROGEs are important regulatory factors for chloroplast-expressed genes in both algae and land plants, there are more characterized ROGEs in algae. This difference could be due to convergent evolution; a similar number of ROGEs may regulate *psaA* in Arabidopsis, but they have not yet been identified because they do not have sequence homology to the Chlamydomonas factors (Ozawa et al., 2020). Alternatively, the regulatory needs of unicellular algae may be different from the regulatory needs of plants. For example, because the photosynthetic machinery in unicellular algae is in growing cells, algae may have a higher need for regulation of protein allocation to the photosynthetic apparatus vs. the rest of the cell. Plants, where the photosynthetic cells are differentiated and not growing, may in turn need more regulation of other processes such as plastid differentiation.

### Much fundamental biology in photosynthesis remains to be discovered

In this study, we identified with high confidence 115 genes required for photosynthesis, 70 of which had not previously been characterized in any organism. More than 65% of these 115 genes have homologs in land plants. In most cases, the functions of these conserved genes appear to be similar in Chlamydomonas and land plants, supporting the value of Chlamydomonas as a model system and expanding the significance of our findings. In several cases, homologous genes appear to have evolved different functions: we identified two such cases, CGL54 (Table 1) and RMT2 (Figure 4D). Approximately 35% of the hits have no clear homologs, which could reflect homolog search failure due to sequence divergence (Vakirlis et al., 2020; Weisman et al., 2020) and/or different evolutionary innovations in the algal lineage such as the algal-specific CO_2_-concentrating mechanism, the study of which has the potential to enhance crop yields (Mackinder, 2018). Thus, much remains to be discovered, and we anticipate that future studies of the previously uncharacterized genes identified here and explored in our proteomics dataset will contribute to further fundamental discoveries in photosynthesis.

## STAR METHODS

### KEY RESOURCES TABLE

| REAGENT or RESOURCE | SOURCE | IDENTIFIER |
| --- | --- | --- |
| Chemicals, Peptides, and Recombinant proteins |  |  |
| UltraPure Low-Melting Point Agarose | Invitrogen | Cat# 16500100 |
| TRIzol™ Reagent | Invitrogen | Cat# 15596026 |
| Guanidine hydrochloride | Sigma | Cat# 369080-1KG |
| Cetyltrimethylammonium bromide | Sigma | Cat# 57-09-0 |
| HEPES | Sigma | Cat# H3375-25G |
| NEM | Sigma | Cat# 128-53-0 |
| DTT | Sigma | Cat# 3483-12-3 |
| EPPS | Sigma | Cat# 000010 |
| Methanol | Fisher scientific | Cat# A456-4 |
| Acetonitrile | Fisher scientific | Cat# A955-4 |
| Chloroform | VWR | Cat# BDH83626.400 |
| H<sub>2</sub>O | VWR | Cat# 87003-652 |
| Formic acid | Sigma | Cat# F0507 |
| NH<sub>4</sub>HCO<sub>3</sub> | Sigma | Cat# 09830 |
| MAX efficiency transformation Reagent for Algae | Invitrogen | Cat# A24229 |
| Critical Commercial Assays |  |  |
| Phusion High-Fidelity DNA polymerase | New England BioLabs | Cat# M0530L |
| MinElute Gel Extraction Kit | QIAGEN | Cat# 28606 |
| Gibson Assembly Master Mix | NEB | Cat# E2611L |
| MT10plex Isobaric Label Reagent Set plus TMT11-131C Label Reagent | Thermo | Cat# A34808 |
| QIAprep Spin Miniprep Kit | QIAGEN | Cat# 27106 |
| Experimental Models: Organisms/Strains |  |  |
| *C.reinhardtii*: wild-type CC-4453 | Chlamydomonas Resource Center | CC-4533 cw15 |
| *C.reinhardtii*: wild-type CC-1690 | Chlamydomonas Resource Center | CC-1690 |
| CLiP library mutants in table S1 | (Li et al., 2019); Chlamydomonas Resource Center | https://www.chlamycollection.org/ |
| *E.coli* Stellar Competent Cells | Takara | Cat# 636763 |
| Chlamydomonas rescued strains listed in Table S7 | This paper, Chlamydomonas Resource | https://www.chlamycollection.org/ |
| Oligonucleotides and Recombinant DNA |  |  |
| pLM005 | (Mackinder et al., 2016); GenBank | KX077945.1 |
| pRAM118 | (Itakura et al., 2019); GenBank | MK357711 |
| Plasmid constructs generated and listed in Table S7 | This paper, Chlamydomonas Resource Center | https://www.chlamycollection.org/ |
| Software and Algorithms |  |  |
| MATLAB | MathWorks |  |
| python |  |  |
| Bowtie 2 | Bowtie | http://bowtie-bio.sourceforge.net/bowtie2/manual.shtml |
| Cutadapt 1.18. | cutadapt | https://cutadapt.readthedocs.io/en/v1.18/ |
| Fiji | (Schindelin et al., 2012) | https://imagej.net/software/fiji/downloads |
| El-MAVEN software | Elucidata | https://www.elucidata.io/el-maven |
| other |  |  |
| Electroporation Cuvette, 2mm gap | Bulldog Bio. | Cat# 12358-346 |
| Ibidi USA μ-Slide 8 well, Glass bottom | Ibidi | Cat# NC0704855 |
| Poly-L-lysine coated glass slides | Sigma | Cat# P0425 |
| Lumigrow Lumibar lights | Lumigrow Lumibar | Cat# 8100-5502 |
| Vacuum filter flask, with a fritted glass support base. | Wilmad Labglass | Cat# BP-1752-001 |
| Nylon membrane filters (0.5μm pore size) | GVS Magna™ | Cat# 1213776 |
| Oasis HLB 96-well μElution Plate, 2 mg Sorbent per Well, 30 μm, 1/pk | Waters | Cat# 186001828BA |
| Electroporator | NEPA GENE | NEPA21 type II |
| SP5 Confocal Microscope | Leica | TCS SP5 |
| Singer Rotor HAD | Singer Instruments | Cat# ROT-001 |
| PhenoBooth imager | Singer Instruments | N/A |
| Typhoon FLA9500 fluorescence scanner | GE Healthcare | N/A |

### CONTACT FOR REAGENT AND RESOURCE SHARING

Further information and requests for resources and reagents should be directed to and will be fulfilled by the Lead Contact, Martin C. Jonikas

### EXPERIMENTAL MODEL AND SUBJECT DETAILS

#### Strains and culture conditions

We performed all experiments on Tris Acetate Phosphate (TAP) TAP or Tris Phosphate (TP) media with revised trace elements (Kropat et al., 2011). TP media had the same recipe as TAP, but the acetic acid was omitted and HCl was added instead to adjust the pH to 7.5. We propagated strains robotically on TAP agar as previously described (Zhang et al., 2014). All mutants used in this study were from the CLiP library (Li et al., 2019). We used the library’s parental strain, CC-4533, as wild-type. We backcrossed mutants to a CC-1690 mt+ transformant carrying a hygromycin resistance cassette (WT-hyg), which has high mating efficiency with the CLiP strains.

We performed spot tests and back-crossing with a subset of 1,781 out of the 3,109 mutants deficient in photosynthetic growth identified previously (Li et al., 2019). This subset had been propagated in the laboratory as colony arrays in 96-colony format since the library’s original construction; whereas propagation of the remaining strains had stopped by the time this study began.

We focused our efforts on characterizing insertions with mapping confidence levels of 1-3 (Li et al., 2019). The 1,781 mutants carried insertions into 1,616 genes mapped with confidence levels 1-3.

### METHOD DETAILS

#### Automated spot tests

We used a RoToR robot (Singer) to replicate colony arrays in 384-colony format from the TAP agar plates on which the 1,781 mutants were propagated onto three agar plates: one TAP, and two TP. We grew the TAP plate in the dark for about a week before imaging; and we acclimated the two TP plates overnight at ∼100 μE/m^2^/s, and then moved them to high light ∼750 μE/m^2^/s for 2-3 days before imaging (using Lumigrow Lumibar lights, catalog number 8100-5502; equal levels of red, blue, and white light). We photographed the plates using a PhenoBooth imager (Singer). We performed the experiment in four replicates: two independent experiments with a technical replicate in each experiment.

To calculate the “normalized colony photosynthetic growth” we analyzed the pictures using MATLAB. We used a MATLAB script to identify and remove the background and to calculate colony size (each green pixel of a colony was given a value 0.5-1 depending on its intensity; and these values were added to obtain the colony size). We then normalized the colony size in each plate by the median size of the 10 largest colonies. We then normalized the size of each colony on the high light plates by the size of the corresponding colony on the corresponding TAP dark plate. We performed the second normalization to rule out mutants with a slow growth phenotype that is not specific to photosynthesis.

### Pooled backcrossing

We performed initial backcrossing experiments with two subsets of mutants labeled MK (26 plates) and AB (10 plates), which together contained the 1,781 mutants, with some mutants being present in both subsets. After obtaining initial results with these subsets, we re-arrayed the most promising mutants in 96-colony format onto four plates labeled NP. The NP plates included 1) mutants containing insertions linked to photosynthetic defects in the initial backcrosses, 2) insertions in genes that were identified as high-confidence hits in our previous study (Li et al., 2019), and 3) mutants that were yellow or brown. Additionally, to check the method’s replicability, we generated a control plate which contained mutations in genes that were not hits and carried insertions whose disruption likely did not result in a photosynthesis defect. The genes disrupted in mutants on the control plate included 1) known flagellar genes and 2) genes that were represented by more than 35 barcodes, no more than 2 of which were hits in our original pooled photosynthesis screen (Li et al., 2019) (in other words, many mutants were available for these genes and the vast majority of these mutants did not exhibit a photosynthesis defect). Using the NP and control plates, we performed a final backcrossing experiment that included two biological repeats of the NP plates and one biological repeat of the control plate.

The backcrossing approach was adapted from the pooled mating (Multiplexed Bulked-Segregant Pool) protocol described previously (Breker et al., 2018). Our protocol is illustrated in Figure S2. Each experimental replicate consisted of the following steps:

1. Mating: We scraped and pooled mt-mutant strains from 96-colony format arrays into flasks containing low-nitrogen gamete-induction medium (Breker et al., 2018). 60-150 colonies were pooled into each 250 ml flask containing 50 ml of gamete-induction medium. We resuspended a similar quantity of WT-hyg into separate flasks containing the same media. We used a cell counter to verify that the strains and the WT-hyg cells were at a similar concentration. Flasks were shaken at 90 RPM for 5-7h in low light (∼40 μE) for mating induction. Then for each flask of mutant strains, 700ul of mutant strains (mt-) and 700ul of WT-hyg were mixed in a 1.5 ml Eppendorf tube, incubated at low light (∼40 μE) without shaking for one hour, then gently spread on two TAP agar plates. The plates were incubated overnight in very low light (∼30 μE). In the morning, the plates were wrapped in aluminum foil and kept in the dark for 7 days.
2. Meiosis: Most of the unmated cells were removed by scraping the agar surface using a sharp razor, and the plates were moved to low light (∼30 μE) for meiosis induction and initial proliferation for ∼5 days. A light microscope was used to check the sporulation efficiency (Jiang and Stern, 2009). The strains were pooled into liquid media (TP) for competitive growth.
3. Light and cassette selections (competitive growth): We added hygromycin to our media to ensure that only backcrossed strains were measured. The mutant library does not have hygromycin resistance, so the original CLiP mutants cannot grow on this media. The WT-hyg strain has hygromycin resistance but does not have barcodes, so it will not affect the barcode counting. We inoculated pooled strains at ∼2 × 10^4^ cells ml^−1^ into TAP + hygromycin (15 μg/ml) 1L bottles for dark growth (3 replicates) and TP + hygromycin (15 μg/ml) 1L bottles for high light growth (3 replicates; except of the 1^st^ experiment where we also did hygromycin (15 μg/ml)+ paromomycin (5 μg/ml) conditions). We bubbled air into the bottles and stirred them using magnetic stirrers at 200 rpm. We exposed the TP cultures to 100 μE for overnight light acclimation, then to 750 μE for the remainder of the growth (using Lumigrow Lumibar lights, catalog number 8100-5502; equal levels of red, blue, and white light). When the cells reached a concentration of approximately 2 × 10^6^ cells ml^−1^, we harvested 10^8^ cells for DNA extraction by centrifugation and flash-freezing the pellet in liquid nitrogen.

| Name | Plates in backcrossing | Competition experiments |
| --- | --- | --- |
| Exp1<br>(AB replicate 1) | AB set (10 plates) | 2 TAP Hygromycin dark and<br>2 TP Hygromycin light;<br>1 TAP hygromycin + paromomycin<br>dark and 2 TP hygromycin +<br>paromomycin light |
| Exp2A<br>(MK 1-12 replicate 1) | 1<sup>st</sup> half of MK set<br>(12 plates) | 3 TAP hygromycin dark and<br>3 TP hygromycin light |
| Exp2B<br>(MK 13-26 replicate 1) | 2nd half of MK set<br>(14 plates) | 3 TAP hygromycin dark and<br>3 TP hygromycin light |
| Exp3A<br>(MK 1-12 replicate 2) | 1<sup>st</sup> half of MK set<br>(12 plates) | 3 TAP hygromycin dark and<br>3 TP hygromycin light |
| Exp3B<br>(MK 13-26 replicate 2) | 2nd half of MK set<br>(14 plates) | 3 TAP Hygromycin dark and 3 TP<br>Hygromycin light |
| Exp4<br>(AB replicate 2) | AB set (10 plates) +<br>3 plates from MK<br>set. | 3 TAP hygromycin dark and 3 TP<br>hygromycin light |
| NP plates | 2 biological<br>replicates of NP set<br>(4 plates) +<br>1 biological repeats<br>of control set (1<br>plate). | For each biological replicate:<br>3 TAP hygromycin dark and 3 TP<br>hygromycin light |

Next, we extracted the DNA and prepared the barcode libraries as described (Fauser et al., 2022), and sent the libraries for Illumina sequencing at the Princeton Genomics Core Facility.

After demultiplexing the we trimmed the initial reads using cutadapt version 1.18. Sequences were trimmed using the command “cutadapt -a <SEQ> -e 0 -q 33 -m 21 -M 23”, where <SEQ> is GGCAAG for 5’ data and TAGCGC for 3’ data. Next, The barcode read counts for each dataset were calculated in python, filtered to only include barcodes present in the original library (Li et al., 2019), and normalized to a total of 1 million.

#### Barcode normalization and growth score calculation

We calculated the “normalized light growth after backcrossing” metric as follows:

1. We used the correlation between the different experimental repeats of each condition to check for swapped samples. Based on these results, we corrected 2 swapped sample pairs: (1) TAP dark sample 3 from Exp3A (MK 1-12 rep2), with TP light sample 1 from Exp2B (MK13-26 rep1); (2) TAP dark sample 1 of NP biological replicate 1, with TAP dark sample 3 of NP biological replicate 2.
2. We averaged the read count of each barcode across the different replicate samples for each condition, using median if we had three replicates or geometric mean if we had only two.
3. To reduce the noise, we removed samples with very low read counts in the TAP condition (<7 in the first experiment and <10 in the rest).
4. We calculated the relative growth as log2 (averaged TP light reads / averaged TAP dark reads). In the first experiment, we had two different conditions; one was grown in hygromycin and paromomycin, and the other only in hygromycin; we analyzed them separately.
5. Normalizing of the NP experiment results – the overall distribution of relative growth rates in the NP experiment was shifted because most of the strains in this competition have a photosynthetic defect, so we scaled the results from this experiment by 0.6 to get a similar distribution to the other experiments.
6. For the final “growth score,” we used the median of the five experiments with the strongest photosynthetic growth defects (for all but 122 genes, it is the same as using all the data). We used the five experiments with the photosynthetic growth defects because there are slightly different conditions between experiments, which can affect the phenotype. Furthermore, in some repeats, we were unable to see an effect because we did not manage to remove all the diploid cells. Lastly, the possibility that the mutants will have a phenotype “by chance” in more than five different experiments is very low, so even slightly lower effects for genes with many experimental repeats can be tolerated. The growth score and the light/dark ratio of backcrossing experiments for all the strains are shown in Table S1. We used the “growth score” to set the 0.34 threshold, to identify hits, and to calculate the FDR (see below, and Figure S2). To reduce noise, we counted as hits only the strains that had reads above the threshold in at least two experiments.
7. FDR calculation (see also Figure S2) – to calculate the False Discovery Rate (FDR) we first estimated how many of the 1,616 mutated of genes in our starting set are required for photosynthesis. We sampled 350 genes at random from the 1,616 and searched the literature for genes among them that are required for photosynthesis. Approximately 6.25% of the genes were known to be required for photosynthesis. Considering previous estimates indicating that approximately half of the genes required for photosynthesis remain to be discovered (Li et al., 2019), we estimate that an additional 6.25% of the genes in the initial set are also required for photosynthesis; thus, we estimate that 12.5% of the initial genes are required for photosynthesis, and the remaining 1,414 (87.5% of the initial 1,620 genes) in our starting set are not required for photosynthesis. Next we defined a set of genes that we called “Genes whose disruption likely did Not Result in a Photosynthesis Defect” (GNRPD). We assigned genes from our set of 1,616 to GNRPD if they were represented by more than 20 insertions, where at most two mutants showed a photosynthetic defect in the Li et al experiment. ∼1% of the GNRPDs (2/204) were among the 136 hit genes identified with a phenotype threshold of 0.34. We assume that the same percentage (∼1%) of the 1,414 estimated genes in our starting set that are not required for photosynthesis in the original mutant set, will go into the hits, resulting in a calculated FDR < 0.11 when using a threshold of 0.34. With a threshold of 0.49, the same calculation yields a FDR < 0.3.

#### Validating insertion sites by PCR

We adapted the check PCR protocol from the CLiP website (https://www.chlamylibrary.org/about), where we used the G1 and G2 primers to validate the existence of the expected insertion (Figure S3). We used the primers suggested for each strain on the CLiP website. We considered the mapping validated if we got a larger PCR product for the mutant than for the wild type, or if we obtained a PCR product for the wild type and not for the mutant in at least 2 experiments (Figure S3).

#### Validating insertion sites by DNA sequencing

The strains were grown in the dark condition, and the DNA was extracted using the same method as above. The DNAs were sent to Princeton Genomics Core Facility for library preparation and whole genome sequencing.

The paired-end 150nt reads were aligned to a reference file that combined the v5.5 Chlamydomonas genome (from Phytozome), the chloroplast and mitochondrial genomes (from NCBI: chloroplast_BK000554.2.gb and mitochondrion_U03843.1.gb) and our CIB1 cassette (Li et al 2019), using the command “bowtie2 –sensitive-local -k 10 -I 100 -X 650 -S”. The resulting SAM files were filtered to extract only read pairs indicating insertion junctions (where the primary alignment was discordant with one side aligning to the CIB1 cassette and the other side aligning to the genome). The resulting genomic positions corresponding to likely cassette insertion positions were clustered (using scipy.cluster.hierarchy.fclusterdata(t=3000, criterion=’distance’, method=’average’)). For each mutant, all clusters containing 4 or more reads were plotted to show the detailed read locations and orientations, as well as the putative insertion positions according to the original library data (Li et al 2019).

Additionally, for each such plot, the concordant read pairs spanning each genomic position were counted and plotted. The resulting plots were evaluated manually to determine the most likely insertion position(s), based on the numbers of matching reads, whether the reads originated from both sides of the insertion position (much less likely for junk fragments), and whether there were concordant read pairs spanning the position (real cassette insertions should not have concordant read pairs spanning them, since the cassette is much longer than the sequenced fragment size).

#### Selection of 115 high-confidence hits

In our experiment, 148 mutants in 136 genes showed normalized light growth after backcrossing that fell below the 0.34 threshold (FDR = 0.1).

First, we validated that the insertions were mapped to the correct genes. We validated the mapping for 119/136 of those genes (87.5%) by PCR and DNA sequencing (Figure 1F and Table S2). The 19 unvalidated genes were removed from the list.

Next, we removed some of the hits to improve the quality of the data set as described below:

1. Six genes (*Cre06.g262900, Cre03.g158950, Cre12.g521450, Cre13.g578600, Cre17.g728700, Cre02.g106950*) were represented by only one mutation that was in a strain that also included a mutation in an established photosynthetic gene or in a gene with multiple hits in our data set. In these cases, we assumed that the phenotype originated from the well-established gene and removed the 2^nd^ gene from the hit list.
2. Five strains had two hits in each (LMJ.RY0402.172741*: Cre13.g584250 + Cre12.g554400,* LMJ.RY0402.187220*: Cre11.g481115 + Cre07.g326010,* LMJ.RY0402.210483*: Cre10.g458700 + Cre03.g211185,* LMJ.RY0402.166642*: Cre03.g155001 + Cre16.g660390 & Cre16.g660430,* LMJ.RY0402.125697*: Cre01.g036400 + Cre01.g015500*). While both genes may be required for the photosynthetic growth, it is more probable that one is the real hit and the other is piggybacking on its phenotype. Hence, we counted them as one and concentrated on the one more likely to be connected to photosynthesis (*Cre13.g584250, Cre11.g481115, Cre10.g458700, Cre03.g155001, Cre01.g015500*). In Table S2, we state the reason for the choice and mention that the effect can be from the other gene.
3. We removed *Cre09.g407650* from the gene hits list because we observed in the proteomic data that *Cre09.g407650* is not downregulated in the corresponding mutant (Figure S5). The insertion in that mutant was in the 3’ UTR, consistent with a mild effect on protein levels.

We then added nine genes as described below:

In our statistical analysis, we looked at genes with insertion mapping confidence levels of 1-3 and excluded confidence level 4 insertions because only 58% of these mutants are correctly mapped (Li et al., 2019). However, there were 3 cases where we did validate the insertion of confidence level 4 hits (LMJ.RY0402.124891: *Cre16.g665750*, LMJ.RY0402.207089: *Cre01.g040050*, LMJ.RY0402.097626: *Cre12.g501550*), so we added those three genes to the hit list.

Last, we added six genes based on manual analysis of the data (LMJ.RY0402.176891: *Cre01.g022681*, LMJ.RY0402.119871: *Cre06.g273700*, LMJ.RY0402.091258: *Cre09.g415500*, LMJ.RY0402.174216: *Cre09.g415700*, LMJ.RY0402.049481: *Cre02.g091750*, LMJ.RY0402.049829*: Cre11.g467573*). In most of these cases, the gene was not a hit in the original analysis because it was not a hit in one replicate, but the replicate is not reliable due to an obvious reason such as very low reads. After removing a problematic experiment, the gene is a hit. In Table S2, we mention in each of these cases why the gene was included in the hit list.

After these edits, our list contained 115 high-confidence genes.

#### Comparison to hits from previous large-scale studies

We compared our 155 high-confidence genes to two sets of hits: 1) previously-identified high-confidence hits, and 2) previously low-confidence hits; which we obtained from three previous large-scale studies (Fauser et al., 2022; Li et al., 2019; Wakao et al., 2021).

**Previously-identified high-confidence hits** consisted of high-confidence hits from (Li et al., 2019) and genes in the photosynthesis clusters in (Fauser et al., 2022). Fauser et al. clustered mutants together based on their phenotype in over 100 different conditions. The work identified two clusters of genes relevant to photosynthesis. The first cluster is the light-sensitive group, where all the hits are relevant to our study; the second cluster is the photoautotrophic light-insensitive. In this second cluster, the clustering is based on phenotypes across many conditions; however, the only condition similar to our experiments is Photoautotrophic 1-3, so we took only the genes whose mutants exhibited pronounced phenotype in this condition: *Cre14.g616600, Cre01.g016514, Cre03.g194200, Cre03.g188700, Cre10.g423500, Cre06.g259100, Cre11.g467712*. We merged the hits from Li and Fauser. This procedure yielded 51 high confidence hits, of which 41 were also high-confidence hits in our study. **Previously low-confidence hits** consisted of a subset of the 260 low-confidence hits from Li et al. (Li et al., 2019) and the 253 low-confidence hits from Wakao et al (Wakao et al., 2021) that were represented in the collection of mutants we analyzed. Neither data set had FDR calculations. While both datasets include genes truly required for photosynthesis, methodological limitations of the studies mean that these datasets also include a substantial number of false positives, making validation by our orthogonal method valuable. In low-confidence hits from Li et al., many of the genes are represented by only one mutant, and others are represented by several mutants but only a small fraction of these mutants shows a photosynthetic phenotype. So, there is a high chance that the photosynthetic phenotype comes from a second-site mutation. In the Wakao study, the authors showed that in most cases their insertion is linked to the photosynthetic phenotype; however, their insertions typically were associated with large deletions that affected several genes. Wakao et al. chose to assign the phenotype to one of the disrupted genes in each of the mutants, primarily based on the literature. Although this connection is often correct, it does not have an experimental/statistical basis.

To create the low-confidence data sets, we first merged the Li and Wakao datasets with 260 and 253 hits respectively. We then took the subset of this merged list of genes that overlaps with the ∼1,616 genes that were included in our initial data set. If a gene was also in the previously-identified high-confidence hits, it was removed from this list. This procedure yielded 219 previously low-confidence hits, of which 31 were high-confidence hits in our study.

#### Mutant gene rescue protocol

The plasmids for complementation were generated as described previously (Wang et al., 2022). 4 of the 16 plasmids were based on the pLM005 backbone, and the remaining 12 were based on the pRAM118 plasmid where the paromomycin resistance cassette was replaced with a hygromycin resistance cassette (Itakura et al., 2019). All plasmids expressed the gene of interest from a PSAD promoter and appended a Venus-3xFLAG tag to the protein sequence.

In the gene rescue protocol, we transformed mutant cells with the linearized plasmid expressing the gene disrupted in the mutant. The linearization and transformation process was carried out as previously described (Wang et al., 2022), until the selection, which was carried out as follows. For plasmids with hygromycin resistance cassette, we used hygromycin-based selection. The cells were plated on 1.5% agar TAP plates with hygromycin (20 μg/ml) and paromomycin (μg/ml) and placed under very dim light for five days, then transferred to light (∼100 μE) for 1-2 weeks until colonies of a sufficient size for picking appeared. For plasmids with paromomycin resistance cassette, we could not use drug selection because CLiP strains already have paromomycin resistance, so we used light selection instead. This selection could be used only for mutants that grow poorly under light conditions. For each of these strains, we included a control where we transformed the mutant with a different plasmid of similar size to determine if transformation with any plasmid could reverse the phenotype, e.g. by creating a second-site suppressor mutation. We only considered a rescue successful when the transformation of the correct gene led to growth under light conditions and the control transformation did not. We plated the cells on 1.5% agar TP plates with paromomycin (20 μg/ml). We gradually increased the light intensity to allow for light acclimation. We left the plate on the shelf overnight for five days under 30 μE, three days under ∼100 μE, and finally 3-4 days under ∼600-700 μE light.

Next, we validated the rescues by robotic spot tests. After the rescue, we picked ∼40 transformants from each rescued mutant to check their photosynthetic phenotype. We used RoToR robot (Singer) to replicate plate with transformants, wild type and mutants to TP and TAP plates, in order to check their growth under TP highlight (800-1100μE) compared to their growth under TAP dark conditions. Then we took 2-4 promising colonies (3 replicates for each) into the plate with wild type and the original mutants (RP 1-4 plates). We used those plates to validate our rescued phenotype. We have at least two independent experiments for each RP plate.

Gene rescue is notoriously challenging in Chlamydomonas due to difficulties with PCR amplification and expression of heterologous genes (Mackinder et al., 2017; Neupert et al., 2020; Zhang et al., 2014), so we perform this part as a “screen”. We used plasmids with the 36 genes we managed to clone (Cre01.g014000, Cre01.g015500, Cre01.g016350, Cre01.g022681, Cre01.g040050, Cre02.g073850, Cre02.g106950, Cre02.g142266, Cre03.g158950, Cre03.g188700, Cre05.g243800, Cre05.g248600, Cre06.g258566, Cre06.g262900, Cre06.g279500, Cre07.g350700, Cre09.g396920, Cre10.g420561, Cre10.g433400, Cre10.g448950, Cre10.g466500, Cre11.g467682, Cre12.g485850, Cre12.g498550, Cre12.g521450, Cre12.g524250, Cre13.g566400, Cre13.g578650, Cre13.g584250, Cre13.g608000, Cre16.g658950, Cre16.g675246, Cre17.g728850, Cre12.g560550, Cre09.g396250, Cre16.g687294), to try a rescue its mutant strain once, and continued with the strains that we managed to rescue. Our success rate of ∼44% is close to the maximum expected even if all were real hits, considering that only 30-50% of transformed constructs express in medium-throughput efforts (Wang et al., 2022). Many of the failed rescues are likely due to challenges with transformation into Chlamydomonas (Mackinder et al., 2017; Neupert et al., 2020; Wang et al., 2022; Zhang et al., 2014), detrimental effects of the GFP tag or the constitutive promoter with some of the genes, and the inherent limitations of our approach, including that rescue of each mutant was only attempted once.

For the rescued mutants, the plasmid used for the rescue, and the Antibiotic resistance, see Table S7.

#### Confocal microscopy

We performed confocal imaging as described previously (Wang et al., 2022). Colonies were transferred to a 96-well microtiter plate with 100 μL TP liquid medium in each well and then pre-cultured in air under 150 μmol photons m^−2^ s^−1^ on an orbital shaker. After ∼16 hr of growth, 10 μL cells were transferred onto an μ-Slide 8-well glass-bottom plate (Ibidi) and 200 μL of 1% TP low-melting-point agarose at ∼35 °C was overlaid to restrict cell movement. Cell samples were imaged using a Leica SP5 confocal microscope with the following settings: Venus, 514 nm excitation with 530/10 nm emission; and chlorophyll, 514 excitation with 685/40 nm emission. All confocal microscopy images were analyzed using Fiji (Schindelin et al., 2012). For each strain, a confocal section through a cell showing the predominant localization pattern was captured and analyzed.

#### Proteomic analysis

Based on our screen results we chose mutants in 100 genes for proteomic profiling (Figure S1 and Table S4). The list includes 3 novel genes that were not in the final hits but are hits in other data sets: *PSR23* and *PIIR2* are high confidence genes in (Li et al., 2019), and *PSR24* is a hit in 2 hit lists: low confidence in (Li et al., 2019) and in (Wakao et al., 2021).

We grew starter cultures in TAP dark for about a week, then moved them to ∼700 ml of TAP (initial concentration ∼ 10^5^ per ml) in 1L bottles and continued growth in the dark. We bubbled air into the bottles and stirred them (using a magnetic stirrer) set to 200 RPM until they reached ∼2×10^6^ cells ml^-1^. We pelleted ∼5X10^7^ cells in 50 ml falcons, transferred the pellets to 1.5 ml tubes, pelleted them again, froze them on dry ice, and stored them at −80 °C.

For each proteomic 11-plex, we prepared 10 samples + a wild-type control. The wild-type control we used in most 11-plexes had been previously harvested in one experiment and frozen in aliquots to reduce the noise between the experiments.

#### Sample processing and mass spectrometry

TMT-labeled (11plex) peptides were prepared mostly as previously described (Gupta et al., 2018). Frozen cell pellets were resuspended in 6 M guanidine hydrochloride (GdCl), 2% cetyltrimethylammonium bromide (CTAB), 50 mM HEPES, 1mM EDTA, and 5mM dithiothreitol (DTT) (pH 7.4). The resuspension lyses the algae to visual homogeneity. Mutant algae cultures grow to different densities and generate pellets of different mass. Diversity in pellet mass was normalized by diluting cells to that of the least dense culture by visual inspection. The final volume ranged from 200-1200 uL. 200 uL of each resuspension was removed to a new Eppendorf prechilled on ice. The lysed algae were sonicated at 20% power for 25 s. Proteins were denatured further at 60 °C for 20 min. After cooling, cysteines were alkylated by the addition of 20 mM N-ethylmaleimide for 30 min, followed by quenching with DTT (10 mM). The protein solutions (200 uL) were charged with 800 uL MeOH, vortexed for 1 min, supplemented with 400 μl chloroform, vortexed for 1 min, followed by addition of 600 μl water and vortexing (1 min). The precipitated proteins were brought to the extraction interface by centrifugation (2 min, 20,800 x g), followed by removal of the upper layer. The protein interface was washed and pelleted from the chloroform phase by the addition of 600 μl MeOH, followed by vortexing (1 min) and centrifugation as described above. The wash solution was removed, and the pellet was washed with 1 ml MeOH. After the removal of MeOH, the pellets were resuspended in 50uL of 6 M GdCl and 10 mM EPPS (3-[4-(2- hydroxyethyl)-1- piperazinyl]propane sulfonic acid) (pH 8.5). The resuspended pellets were frozen.

Pellets were thawed and their protein concentrations quantified using the BCA assay from Pierce with the BSA standard curve diluted in 10 mM EPPS pH 8.5 6M GdCl. 30 ug of each pellet was diluted to 15uL with 10mM EPPS pH 8.5 in 6M GdCl. The 15 uL of 2 μg/μL denatured protein solution was diluted with 75 uL 20 ng/μL LyseC in 10mM EPPS pH8.5, vortexed and allowed to digest overnight at room temperature. A second round of digestion followed with the addition of 270 μL of 20 ng/μL each LyseC and Trypsin in 10 mM EPPS pH 8.5, vortexing and overnight incubation at 37C. The solvent was removed under reduced pressure in a SpeedVac and resuspended in 30 ul of 200 mM EPPS (pH 8.0) to a concentration of 1 g/L. Ten microliters were removed from each resuspension and charged with 2μl of different TMT-isobaric mass tag N-hydroxysuccinimide (NHS) ester (20 g/liter). The acylation proceeded overnight at RT and was quenched at RT with 0.5 uL of 5% hydroxylamine for 20 min, followed by 1 uL of 5% phosphoric acid.

Peptides were enriched from the acidified TMT labeling reactions by solid-phase extraction using a Waters Oasis HLB Elution 96-well plate (3 mg/well). One well per multiplexed quantitative proteomics experiment was wetted with 400uL MeOH and then hydrated with 200uL water. The 11 labeling reactions are pooled and diluted into 400ul and allowed to adsorb HLB resin under gravity flow. The adsorbed peptides were washed with 100 μL water, followed by centrifugation for 1 min at 180 rpm. The peptides were eluted with sequential additions of 100 μl of 35% acetonitrile (1% formic acid [FA]) and 100 μl of 70% acetonitrile (0.1% FA). Eluent solvent was removed under reduced pressure in in a SpeedVac. The peptides were resuspended in 20 uL of 1% FA and subjected to quantitative multiplexed proteomics by nano-ultraperformance liquid chromatography-tandem mass spectrometry (nanoUPLC-MS/MS).

Peptides were separated on a 75 μm inner diameter microcapillary column. The tip for the column was pulled inhouse and the column was packed with approximately 0.5 cm (5 μm, 100 Å, Michrom Bioresources) followed by 40 cm of Waters BEH resin (1.7 μm, 120 Å). Separation was achieved by applying a 3−22% Acetonitrile gradient in 0.125%, formic acid with 2% DMSO over 165 min at ∼300 nL/min. Electrospray ionization was enabled by applying a voltage of 2.0 kV through an IDEX high-pressure fitting at the inlet of the microcapillary column. TMT3 data collection was performed as previously described (Gupta et al., 2018). The instrument was operated in data-dependent mode (10 ions/scan) with an MS1 survey scan performed at a resolution setting of 120k (m/z 200) with a scan range of m/z 350 to 1,350, an RF (radio frequency) lens of 60%, automatic gain control (AGC) target of 106, and a maximum injection time of 100 ms. Ions with charge states 2-6 were filtered by intensity with a threshold of 5e3. A dynamic exclusion window of +/-10ppm for 90s was used. MS2 quadrupole isolated ions (0.5 isolation window) were activated with CID at 35% collision energy and Q 0.25 and analyzed in the ion trap with an AGC target of 1.5e4 and 75ms maximum injection time. 10 data dependent MS3 synchronous precursor selections (2 isolation window) were selected from range 400-2000 m/z. The MS3 activation is HCD with 55% collision energy. The ions are analyzed in the orbitrap at 50,000 resolution with an AGC of 1.5e5 and an maximum injection time of 100 ms.

#### Mass spectrometry data analysis

Mass spectrometry raw data was analyzed using GFY software licensed from Harvard (Nusinow et al., 2020) to quantified proteins relative abundance.

We normalized each protein’s abundance in each sample by that protein’s abundance in the corresponding wild type sample, then normalized the protein’s abundance in the sample by the sample’s median to account for systematic difference likely coming from technical difference in the amounts of proteins entered into the TMT labeling. To decrease the noise between the different 11-plexes (Figure S5A-S5B) we normalized each protein by its median in the 11-plex. This dramatically decreased the noise while maintaining most of the signal (Figure S5C).

#### Chloroplast RNAseq

The RNA seq experiments were split into two experiments; each experiment had its own wild type. In each experiment, we had 2-3 replicates for each mutant strain and 2-4 replicates for the wild type.

The strains were grown in the same conditions as for the proteomic analysis. When the cultures reached ∼ 2×10^6^ cells/ml, we pelleted 13 ml of culture in 15 ml round Falcon tubes. We then used TRIzol extraction (following manufacturer’s protocol) to obtain the total RNA. The RNA was sent to Princeton Genomics Core Facility for RNAseq and Next Generation Sequencing. The chloroplast mRNA does not have polyA, so they used the Qiagen FastSelect – rRNA Plant Kit for rRNA depletion. Then generated libraries using PrepX™ RNA-Seq for Illumina Library kit to generate the library for RNAseq.

mRNA analysis: First non-coding RNA sequence was filtered out: each dataset was aligned (using the bowtie2 –fast command) against the dataset of non-coding RNAs (Gallaher et al., 2018), and only unaligned reads were included in the rest of the analysis. Next the reads were aligned against a reference file containing the updated chloroplast and mitochondrial genomes (Gallaher et al., 2018), a set of Chlamydomonas rRNA sequences (downloaded from https://www.arb-silva.de/), and Chlamydomonas nuclear coding sequences (v5.5 from Phytozome, file Creinhardtii_281_v5.5.cds_primaryTranscriptOnly.fa), using the bowtie2 --fast option. For each sample, the number of reads in each chloroplast gene was calculated in python, with each side of each read considered separately, and with gene positions based on the chloroplast gff3 file from (Gallaher et al., 2018).

The reads were used to estimate the mRNA levels of the different chloroplast-expressed photosynthetic genes. The reads were normalized by the total chloroplast gene reads. Our RNA seq reads were Paired-end, allowing us to create a second analysis of where each side maps on the genome. For example, this allowed us to count the number of mRNAs where one side is in exon 1 and the other in exon 3. The overall coverage was much higher in our second experiment, so we normalized the 1^st^ experiment using the wild-type ratio between the experiments, allowing us to present them together.

#### Nuclear RNAseq

The mRNA of *pmr1* (2 independent experiments) and wild type (2 independent experiments) was also used for polyA-based RNAseq. The library preparation and Next Generation Sequencing were done at Princeton Genomics Core Facility.

The paired-end reads were aligned against the primary transcriptome (v5.5, from Phytozome) using the bowtie2 --fast command, and the number of reads aligning to each transcript were counted in python for each sample.

We normalized the number of reads to 50M then we averaged (using geomean) the 2 experimental repeats of *pmr1* and the 2 experimental repeats of wild type, and then calculated the relative reads by log2(*pmr1*/ wild type).

#### Metabolomics analysis

The protocol was adapted from (Yuan et al., 2008). In short, we grew starter cultures at TAP dark for about a week, then moved to ∼700ml of TAP (initial concentration ∼ 10^5^ per ml) in 1L bottles kept in the dark. We bubbled air into the bottles and stirred them (using a magnetic stirrer) set to 200 RPM until they reached ∼2×10^6^ cells ml^-1^. We harvested ∼ 10^7^ cells using vacuum filter, and immediately dunked the filter’s membrane into 1.5 ml of 40:40:20 (v/v/v) methanol:acetonitrile:H_2_O solution with 0.5% formic acid to extracted the metabolites. All reagents were precooled to −20C and the protocol was done on ice. After neutralizing by NH_4_HCO_3_ (132 µL) and pelleting, we took 100ul supernatant for LC-MS.

The LC-MS method was modified from (Yang et al., 2022). Water-soluble metabolite measurements were obtained by running samples on the Orbitrap Exploris 480 mass spectrometer (Thermo Scientific) coupled with hydrophilic interaction chromatography (HILIC). An XBridge BEH Amide column (150mm X 2.1 mm, 2.5 uM particle size, Waters, Milford, MA) was used. The gradient was solvent A (95%:5% H_2_O:acetonitrile with 20 mM ammonium acetate, 20 mM ammonium hydroxide, pH 9.4) and solvent B (100% acetonitrile) 0min,90% B; 2min,90% B; 3min,75% B; 7min,75% B; 8min,70% B; 9min, 70%B; 10 min, 50% B; 12 min, 50% B; 13 min, 25% B; 14 min, 25% B; 16 min, 0.5% B, 20.5 min, 0.5% B; 21 min, 90% B; 25 min, 90% B. The flow rate was 150 mL/min with an injection volume of 5 mL and a column temperature of 25 °C. The MS scans were in polarity switching mode to enable both positive and negative ions across a mass range of 70–1000 m/z, with a resolution of 120,000. Data were analyzed using the EI-MAVEN software (v 0.12.0, Elucidata).

We included a total of 3 replicates from each strain from 2 independent experiments.

## SUPPLEMENTAL INFORMATION

Supplemental Information includes 9 figures and 7 tables.

## AUTHOR CONTRIBUTIONS

M.K. and M.C.J. conceived the project. M.K and W.P. performed data analysis. M.K. grew strains for mass spectrometry. L.M. and M.W. prepared samples, performed mass spectrometry, and established the protein quantification pipeline. M.K. and A.G performed spot tests. M.B., F.R.C. and M.K. established the pooled backcrossing method. M.K. performed the pooled backcrossing experiments. M.K., G.G. and A.G. performed insertion mapping validation by colony PCR and sequencing. L.W., M.K., A.K.S., S.E.G. and A.T.W. performed mutant rescue and protein localization by confocal microscopy. M.K., A.R., and J.D.R. performed and analyzed the metabolomic experiments. C.D.M conducted the prediction of protein structure. M.K. and M.C.J. wrote the manuscript with input from all authors.

## ACKNOWLEDGMENTS

We thank Michelle Warren-Williams for media preparation and assistance with propagating strains; the Princeton University genomic core facility and its manager Wei Wang for their help with DNA and RNA sequencing and library preparation; Princeton University Confocal Microscopy manager Gary Laevsky for instrumentation support; members of the Jonikas laboratory and Felix Willmund for helpful discussions; Olivier Vallon, Yana Kazachkova, Silvia Ramundo, Shan He, Alice Lunardon, Jessica H. Hennacy, Sabrina Ergun, Moritz T. Meyer, Eric Franklin for feedback on the manuscript; and Marie Bao, as part of Life Science Editors, for help with editing the manuscript. The project was funded by the Princeton Catalysis Initiative, U.S. National Institutes of Health grant R35GM128813, U.S. National Foundation grant MCB-1914989, European Molecular Biology Organization fellowship ALTF 1006-2017, Human Frontier Scientific Program fellowship LT000031/2018-L, HHMI/Simons Foundation grant 55108535, and the Lewis-Sigler Scholars Fund. Martin Jonikas is a Howard Hughes Medical Institute Investigator.

## SUPPLEMENTAL FIGURES

**Figure S1:**
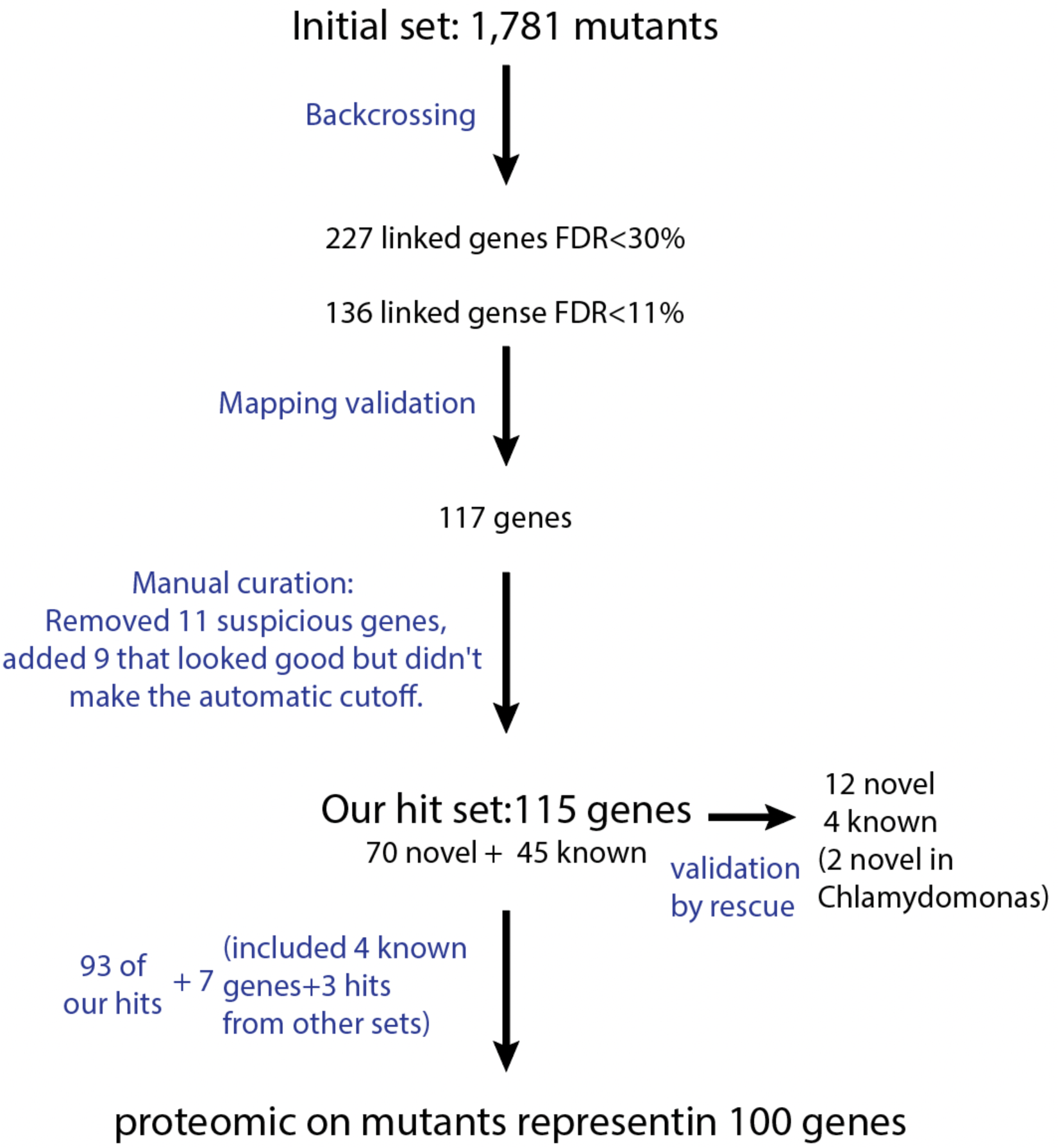
Summary of mutant and gene numbers at different stages of this project, related to Figures 1-6. For a detailed description of the process, please see the main text and STAR Methods. Mutant and gene IDs are provided in Supplementary Table S1.

**Figure S2.**
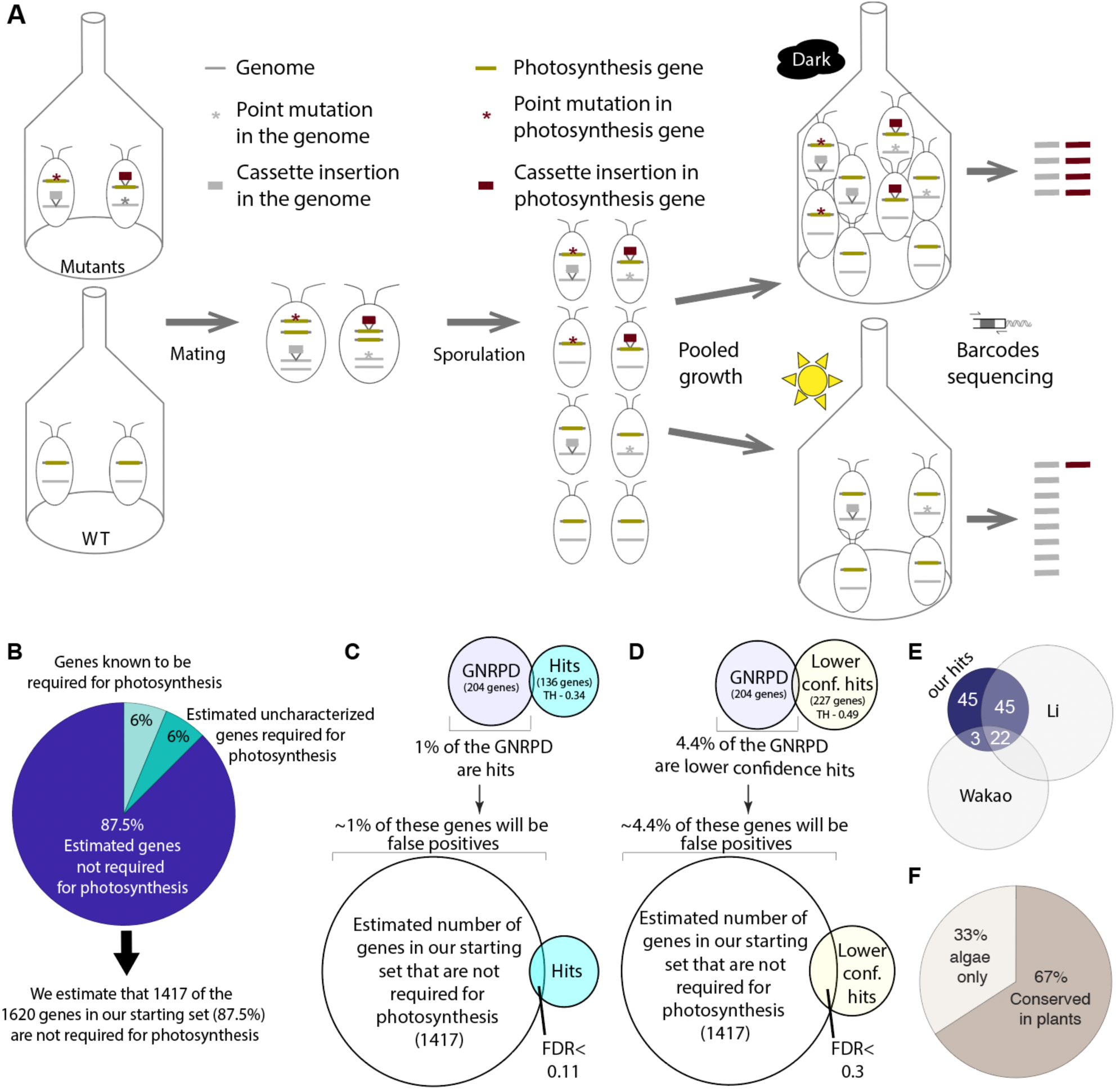
Pooled backcrossing and False Discovery Rate calculation, related to Figure 1. **(A)** We mated paromomycin-resistant mt-mutants with a hygromycin-resistant mt+ strain. **The mutants carried cassette insertions and additional mutations.** The resulting progeny included mixed genotypes where the insertions and the second site mutations were separated. We grew the progeny under a dark control condition, where all viable strains grew; and photoautotrophically, where mutants in genes required for photosynthesis were depleted. By sequencing the pools of barcodes associated with insertions, we could identify barcodes that were depleted under the photoautotrophic condition, and thus were genetically linked to genes required for photosynthesis. (B) Calculation of the “estimated number of genes in our starting set that are not required for photosynthesis”. Our dataset included 1,616 genes with confidence level <4. We sampled 350 genes at random from the 1,616 and screened the literature for genes among them that are required for photosynthesis. 6.25% of the genes were known to be required for photosynthesis. Considering previous estimates indicating that approximately half of the genes required for photosynthesis remain to be discovered (Li et al., 2019), we estimate that an additional 6.25% of the genes in the initial set are also required for photosynthesis; thus, we estimate that 87.5% of the genes in our starting set are not required for photosynthesis. Given these numbers, the “estimated number of genes in our starting set that are not required for photosynthesis” is 1414 (87.5% of the initial 1,616 genes). (C) The False Discovery Rate (FDR) calculation is based on a set of specific genes that we called “Genes whose disruption likely did Not Result in a Photosynthesis Defect” (GNRPD). Genes from our set of 1620 genes were assigned to GNRPD if they were represented by more than 20 insertions in Li et al experiment and at most two mutants showed a photosynthetic defect. ∼1% of the GNRPDs were among the 136 hit genes identified with a phenotype threshold of 0.34. We assume that the same ratio (∼1%) of the “estimated number of genes in our starting set that are not required for photosynthesis” (see B) in the original mutant set will go into the hits, resulting in FDR <0.11. (D) The same calculation as (C) only for lower-confidence hits (threshold of 0.49). Those genes have a higher false-discovery rate, but they still include many genes genuinely required for photosynthesis. (E) 25 of our 115 hits (22%) were also hits in (Wakao et al., 2021), and 70 of the 115 (61%) were also hits in (Li et al., 2019) (F) More than 65% of our hits are conserved in land plants.

**Figure S3.**
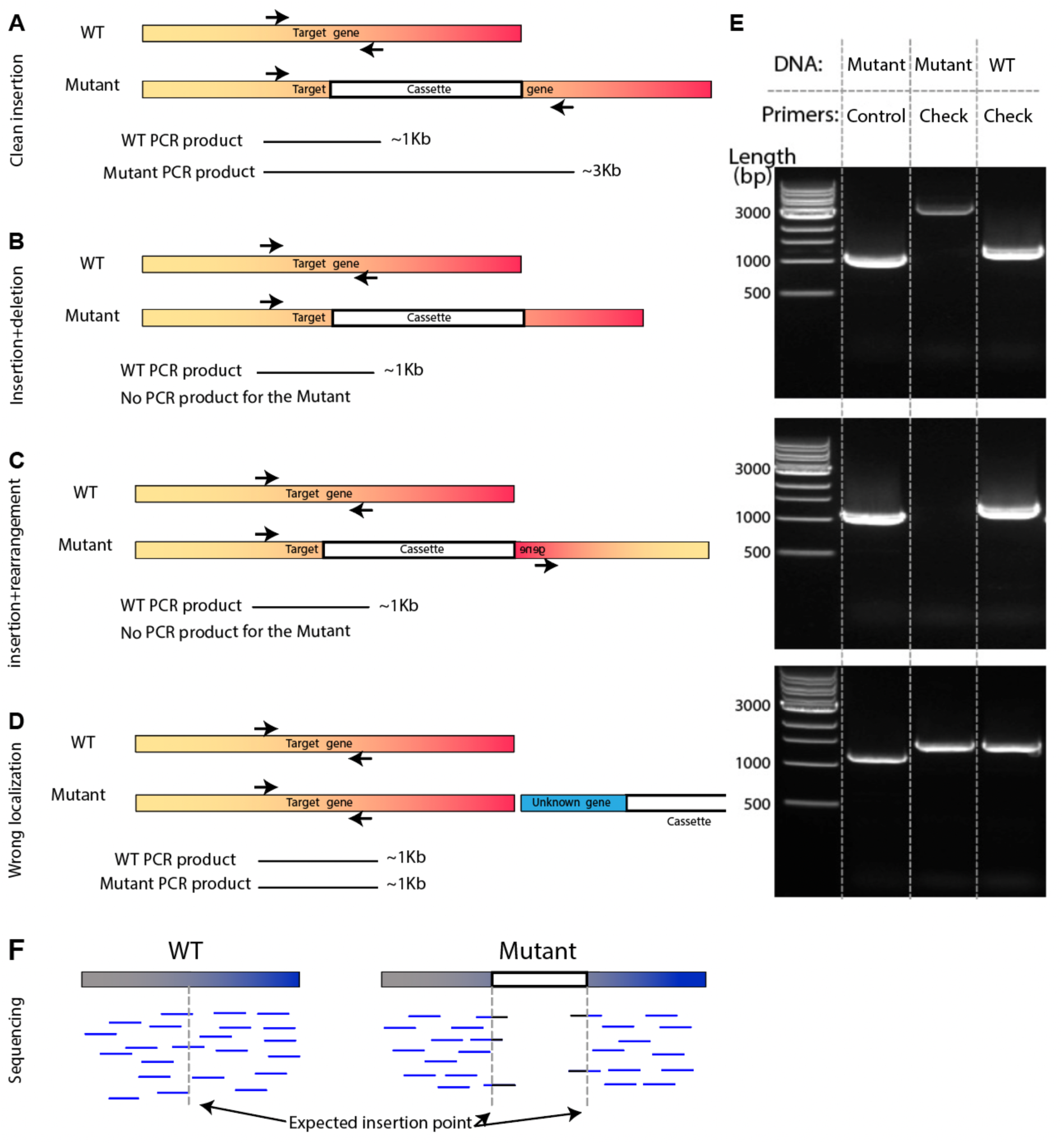
Mapping validation by colony PCR and sequencing, related to Figure 1. (A-D) Four options of cassette insertion and the expected PCR product. (A) Clean insertion – the cassette integrates into the genome cleanly; in this situation, the PCR product of the mutant will be about 2Kb longer than the WT product. (B) Insertion with significant deletion – in this case, the deletion associated with the insertion removed one of the PCR primers, so we will get the PCR product for WT but not from the mutants. (C) Insertion with rearrangement – in this case, the primer sequence is there but lost its orientation, so again we will get PCR products for WT but not for the mutant. (D) When the insertion isn’t in our expected gene, we will get the same length of PCR product from the WT and the mutants. (Note that we can get this pattern also if the insertion is associated with a deletion of a similar size). (E) Example of colony PCR results. The control lane is amplified mutant DNA using control primers to verify the mutant DNA quality. In the upper example, the mutant is ∼2Kb longer than the WT, as expected from a clean insertion (A). In the middle example, we have a band for the WT but not for the mutants. Such a result was taken to validate an insertion site if it was reproduced at least twice, and is expected for scenarios (B) and (C). The lower example was counted as a failure to validate the mapping and is expected for (D). Note that when we fail to get a product with WT we used different primers or the sequencing method. (F) Mapping validation by sequencing (The paired-end 150nt reads). A positive mapping is where we found in the expected area a chimera reads (one side mapped to the genome and another to the cassette) and a “hole” in the genome coverage. For more details see STAR Methods.

**Figure S4.**
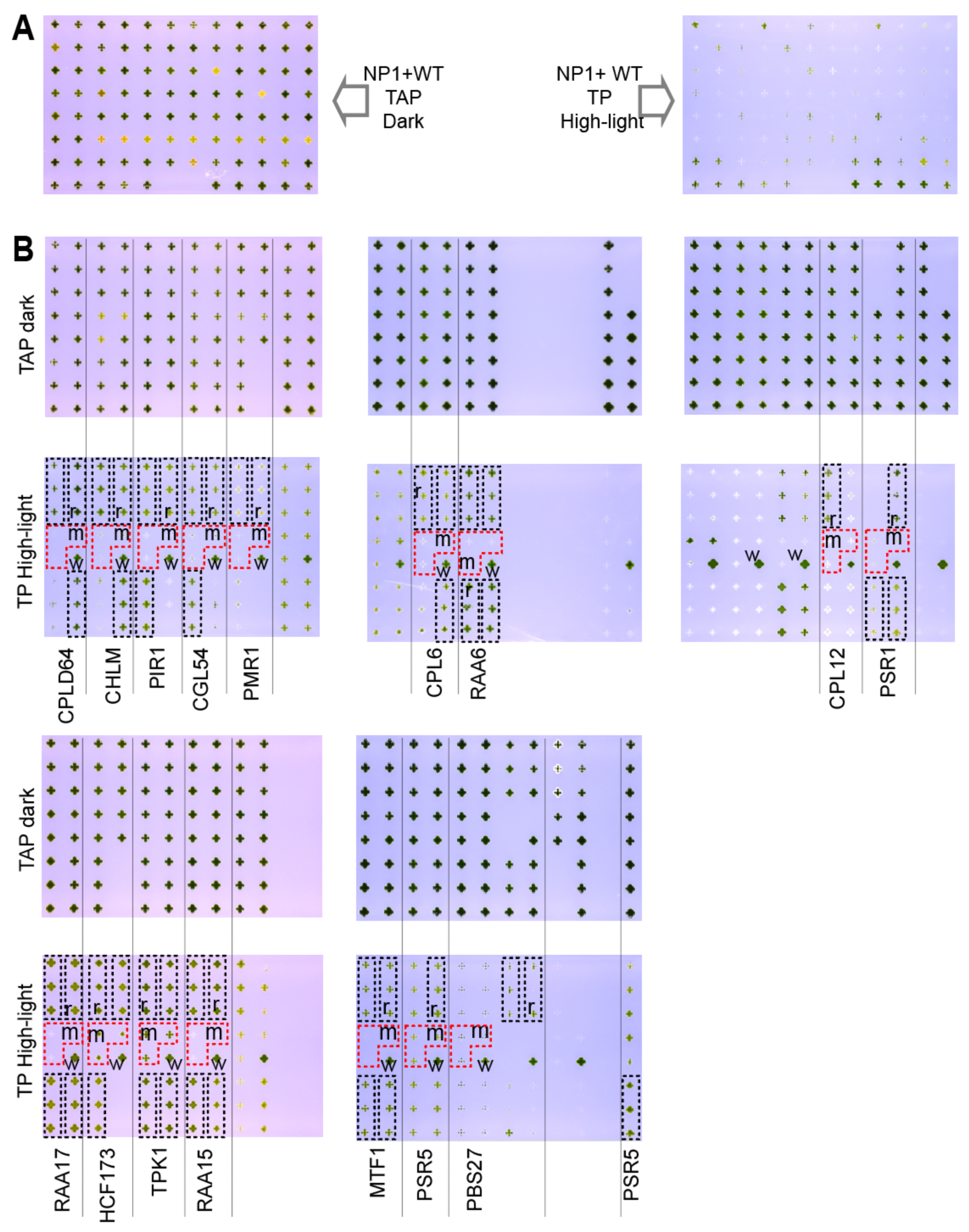
Unprocessed images for Figures 1, 2, and 6, related to Figures 1, 2, and 6. (A) The unprocessed image for Figure 1B. (B) The unprocessed image for Figures 2 and 6. In each high-light plate, the three copies of the original mutants are outlined in dashed red and every triplicate of the rescued strains is outlined in dashed black. To reduce the effect of location on the plate, we put one WT next to each mutant trio. the “r” indicates the rescued strain used in the main figure. Similarly, “m” indicates the mutants and “w” the WT used in the main figure. There are differences in the rescue efficiency between the different rescued strains, even in the same mutant. Many parameters could contribute to those differences, including insertion site and expression level.

**Figure S5.**
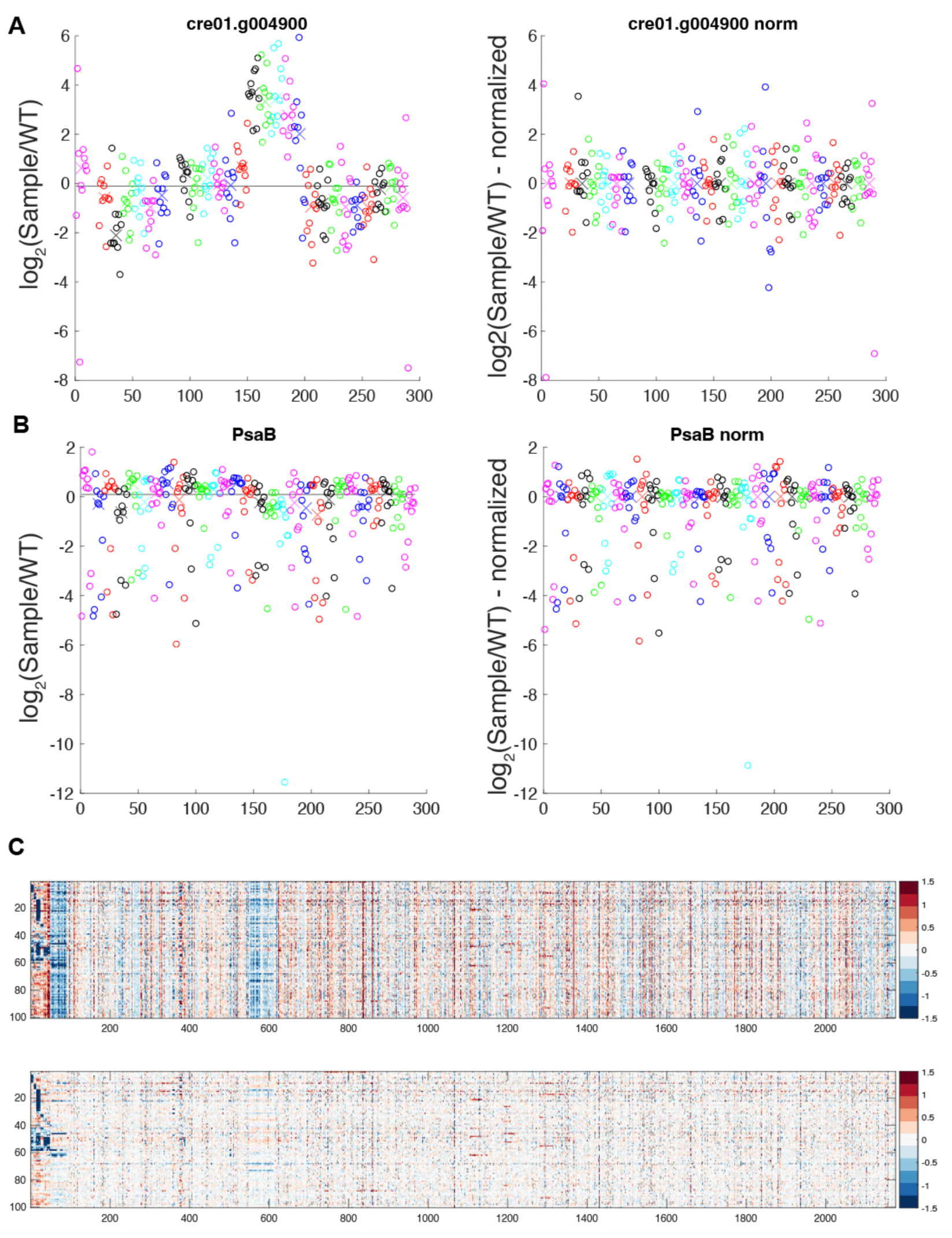
Normalization of the proteomic data, related to Figures 3-4. (A-B) Example of the data of two proteins (Cre01.g004900 (A) and PsaB (B)) before and after 11-plex-median-based normalization. Each group (10 samples, grouped by color, and the group median shown as x in the same color) is the data from one proteomic 11-plex (10 samples and WT). We can see a difference between the groups (11-plexes), which we removed by normalizing using the group median. The black lines represent the median of all the samples. (C) The normalization reduces the noise of the data. Protein levels are shown for proteins measured at least at 65% of the experiments in the 100 mutants. The data is the average of two repeats on the log_2_ scale. The upper panel is before and the lower panel is after the 11-plex-median-based normalization. We can see that the normalization removes much of the noise and maintains most of the signal. The first ∼90 proteins are the ones shown in Figure 4.

**Figure S6.**
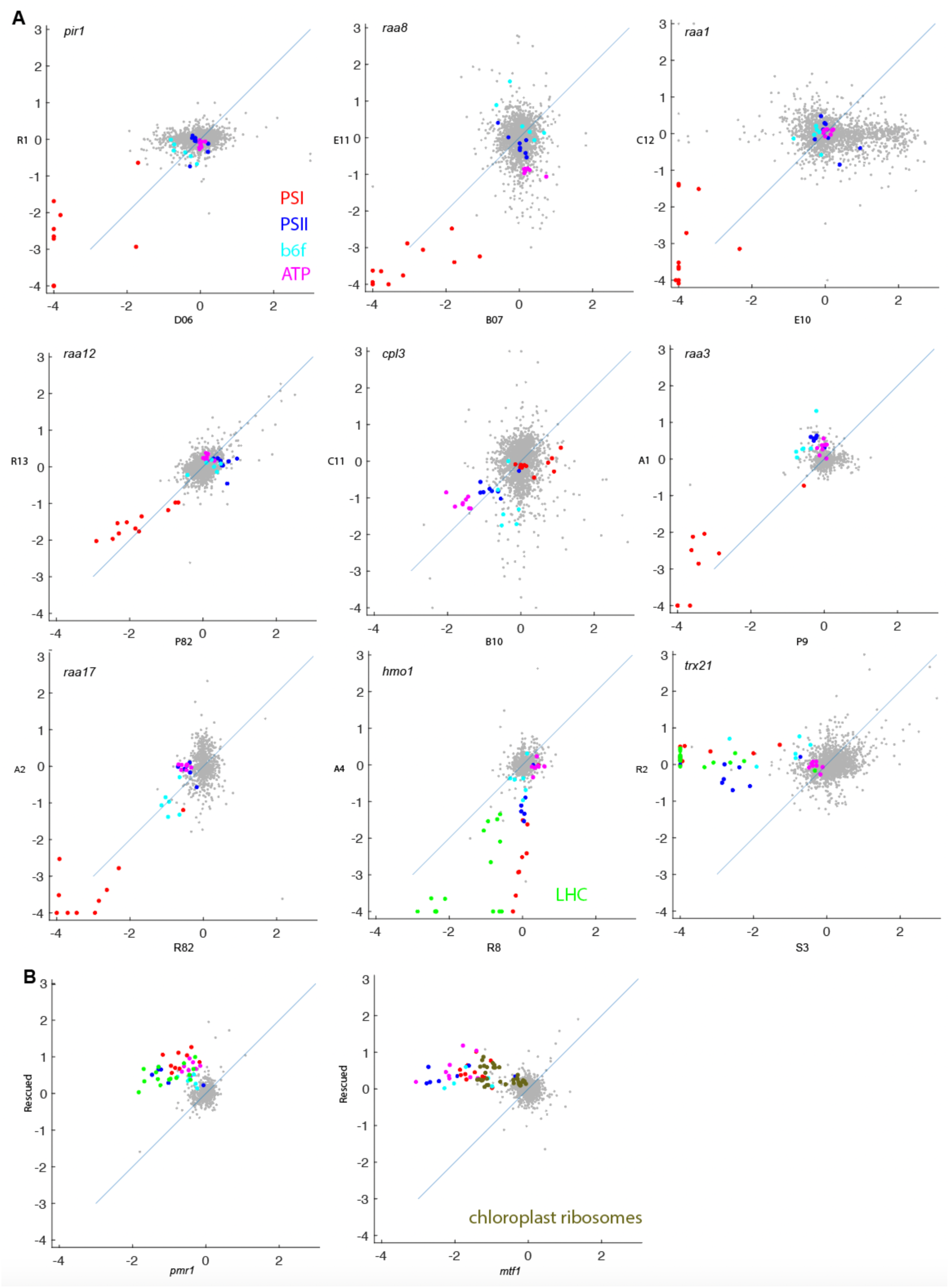
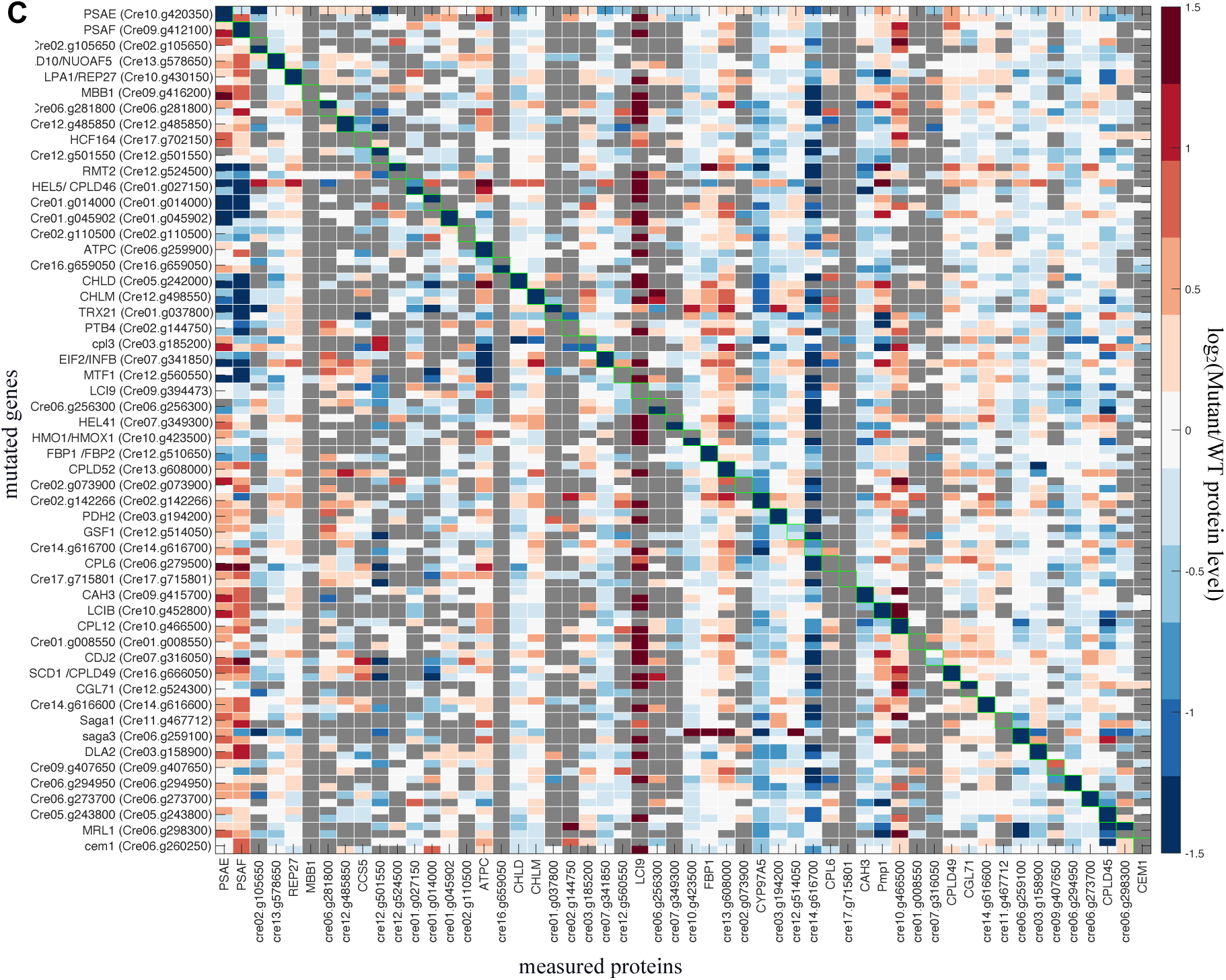
Proteomic controls, related to Figure 4. (A) Genes for which have 2 mutant alleles in our hit set, and we collected (meaningful) proteomics data for both mutant alleles. Each axis represents one allele’s log_2_(mutant/WT) proteomic data. The sample name is shown near each axis. (B) Genes for which we rescued their mutants and collected proteomics data for both the mutants and the rescued strains. In all cases, our data suggest that the impact on the photosynthetic complexes is from our mutant gene, except in the case of TRX21. The two mutants have different phenotypes: one was yellow and had a decreased abundance of chlorophyll-binding proteins (including PSII), and the other was green and only affected PSII-suggesting that the yellow mutant has an additional mutation leading to the additional proteomic phenotype. Additionally, 5 genes (HCF173, CPLD64, CHLM, RAA6, RAA17) showed strong proteomic and photosynthetic phenotypes, and their rescue restored the mutant to WT-like growth. This demonstrates that only in rare cases (1/16) does the prominent proteomic phenotype come from a second mutation. (C) Validation that the protein is absent from its mutant. We can see downregulation of the proteins in all the samples (when we have protein in our data set) except for Cre09.g407650, suggesting that Cre09.g407650 is a false positive. The insertion in Cre09.g407650 is in the 3’ UTR and was linked to the phenotype; this insertion is likely not the reason for the photosynthetic phenotype, demonstrating how proteomics can help identify false positives.

**Figure S7.**
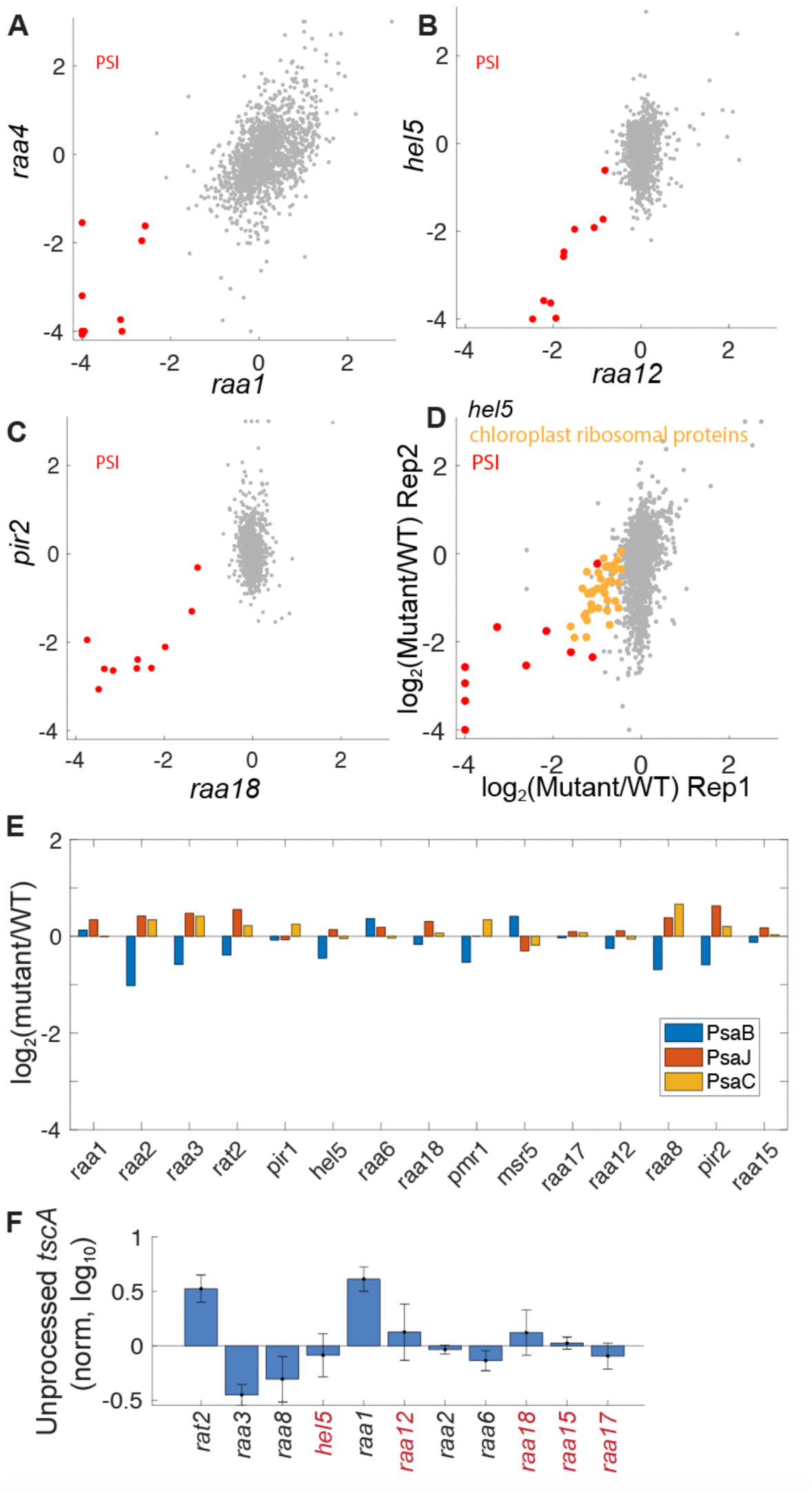
Supplemental data for PSI regulators, related to Figure 5. (A-C) Scatterplots of proteomic data in mutants in known *psaA* maturation factors (RAA1, RAA4) and mutants in novel genes with similar proteomic profiles (HEL5, RAA12, PIR2 and RAA18). The data reflect the average normalized log_2_ (mutant/WT protein abundance) from two independent experiments. (D) The proteomic data of *hel5* mutants. (E) The mRNA levels (normalized to WT) of *psaB,J,C* in the different mutants. The only effect is on *psaB* levels and it is less than two-fold, which should not affect translation levels (Choquet and Wollman, 2002). (F) RAT2 and RAA1 are required for *tscA* processing. This requirement (Balczun et al., 2005; Merendino et al., 2006) suggests that *tscA* processing is carried out in conjunction with the splicing complex organized around RAA1. Error bars represent standard error of the mean.

**Figure S8.**
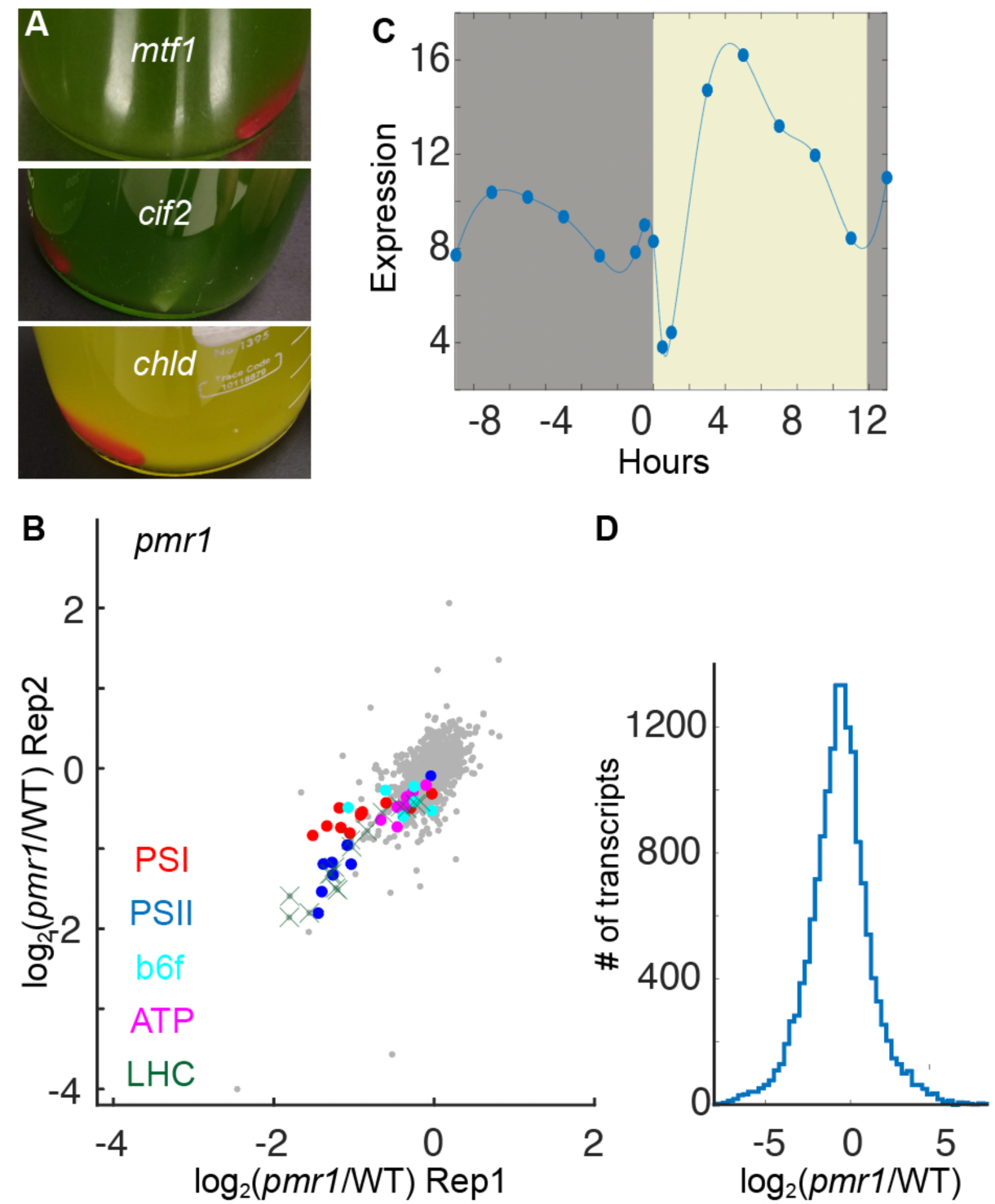
Supplemental data for the master regulators, related to Figure 6. (A) Images of strains grown in the TAP dark condition. Note that mtf1 and cif2 are green, whereas chld (mutant in chlorophyll formation) is yellow. (B) pmr1 mutant proteomic effect. (C) PMR1 periodic expression. The light period is shown in yellow, and the dark period is shown in gray. The data are from (Strenkert et al., 2019). (D) A histogram of pmr1 transcriptome.

**Figure S9.**
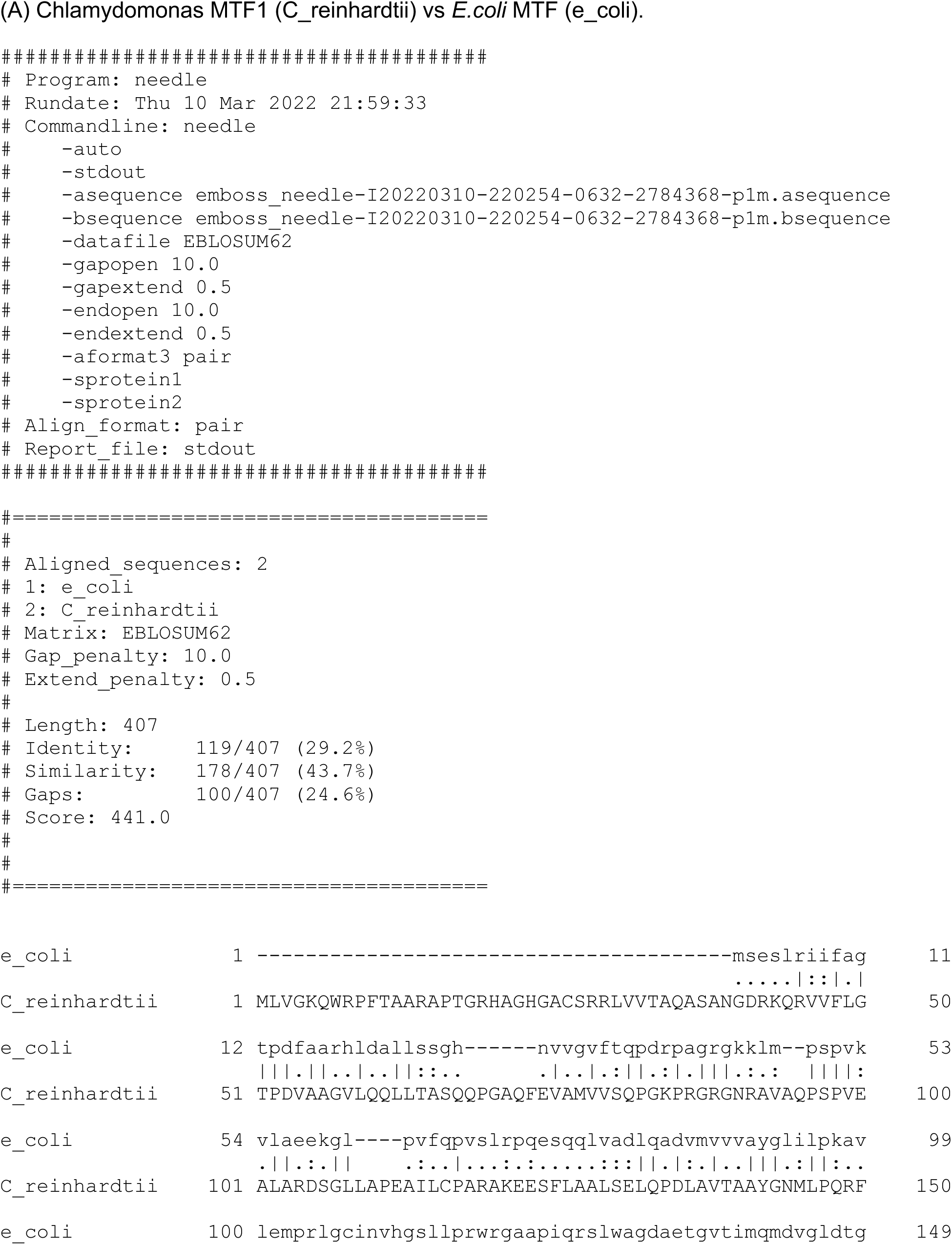
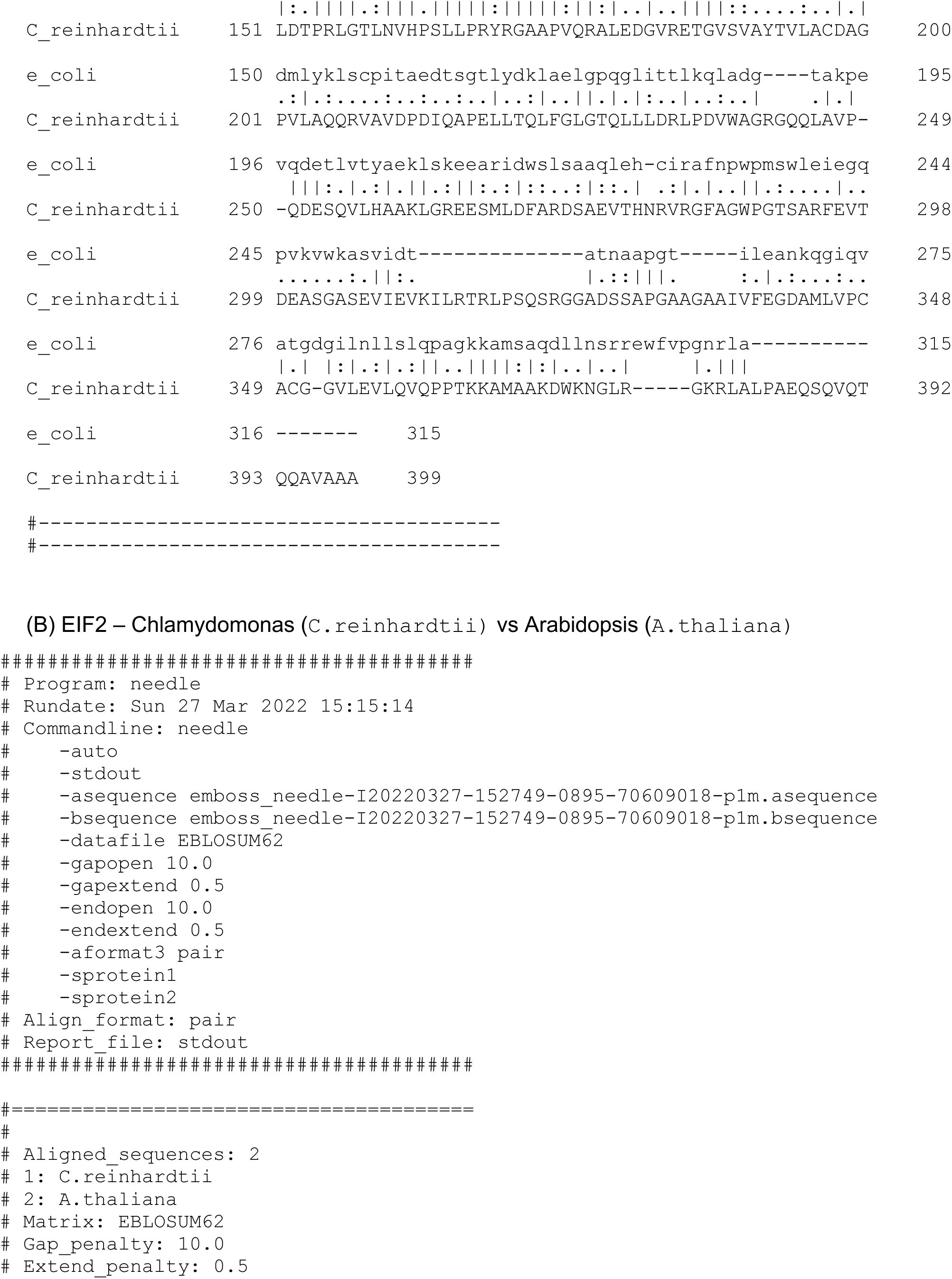
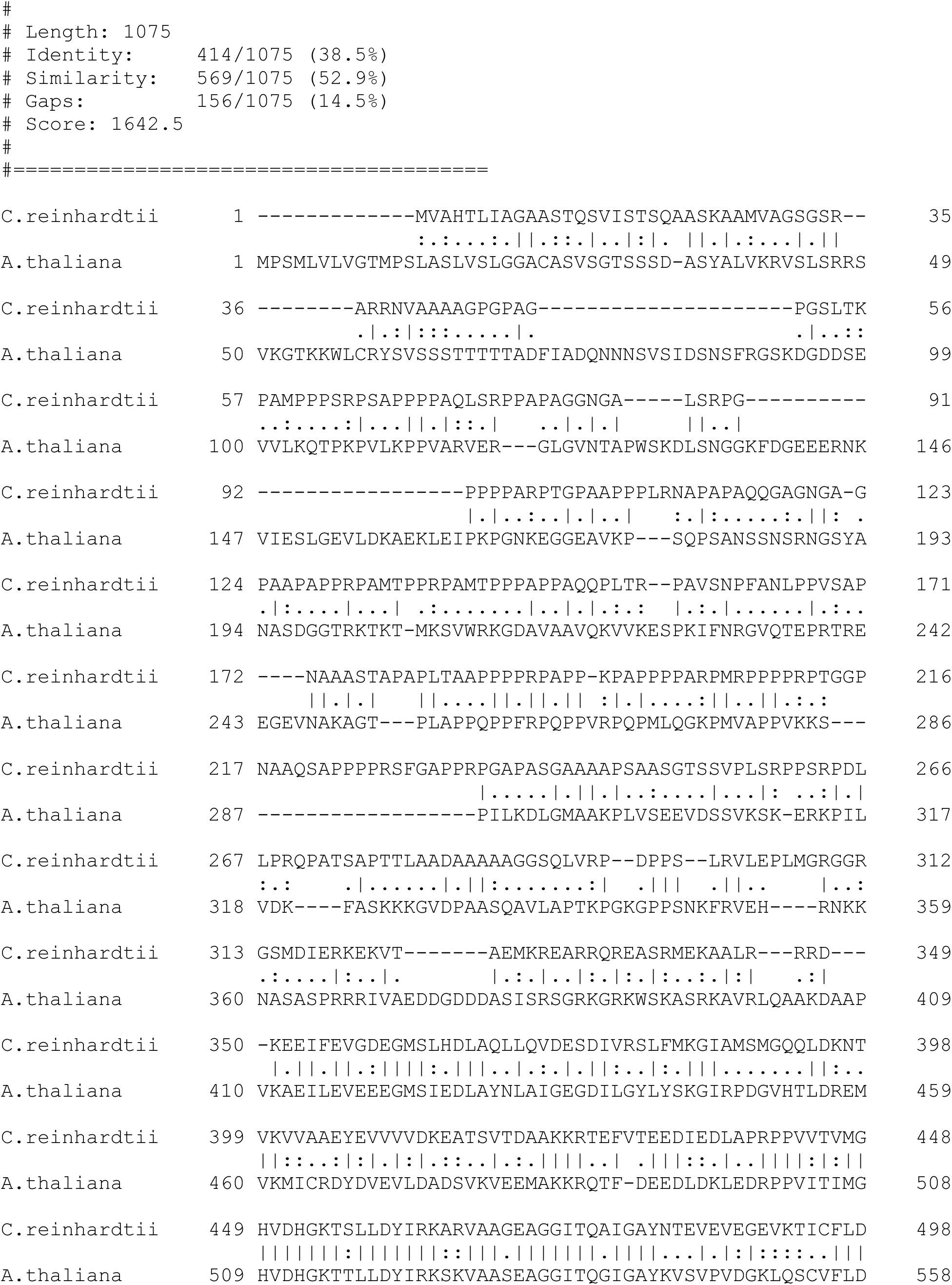
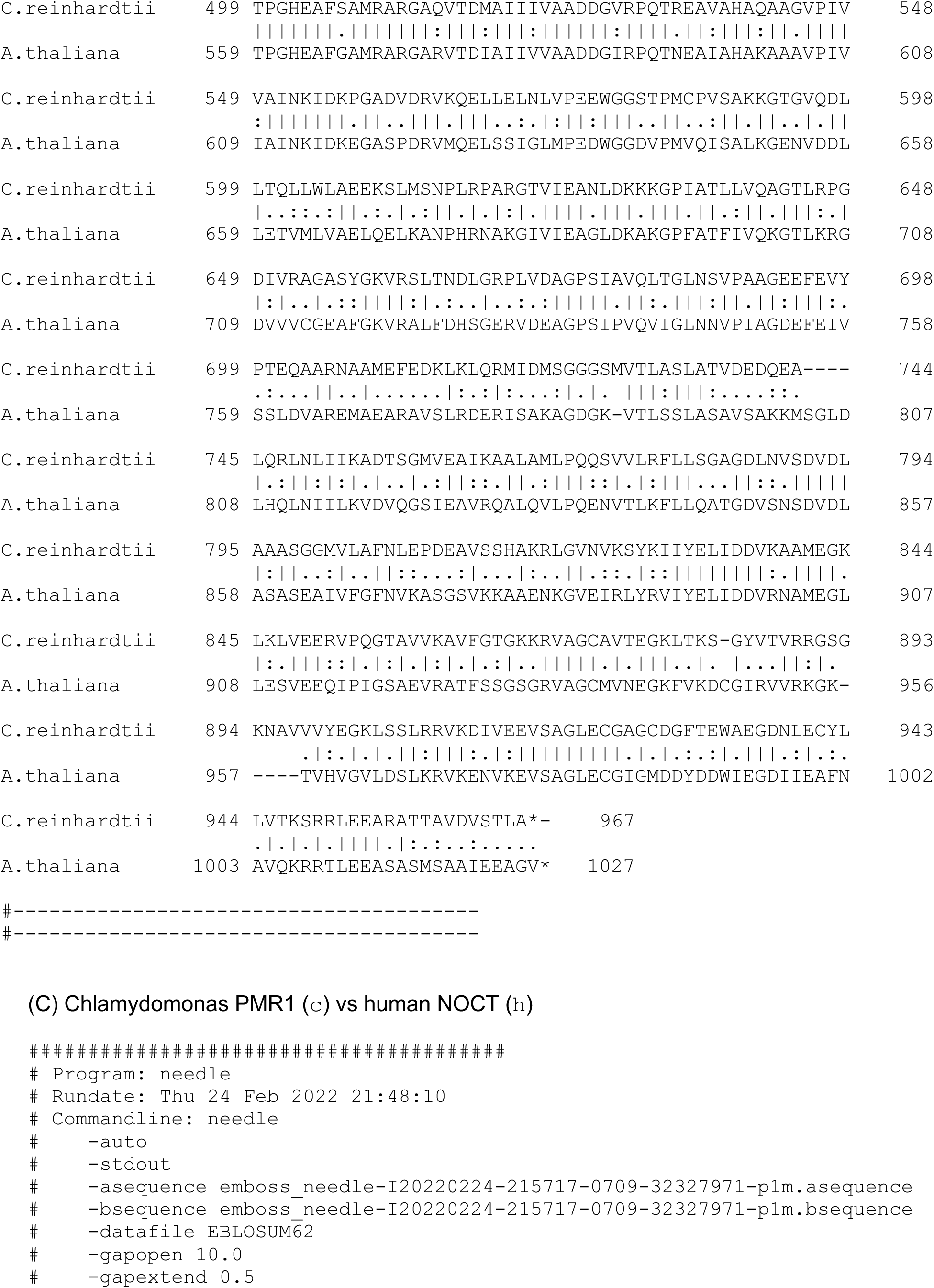
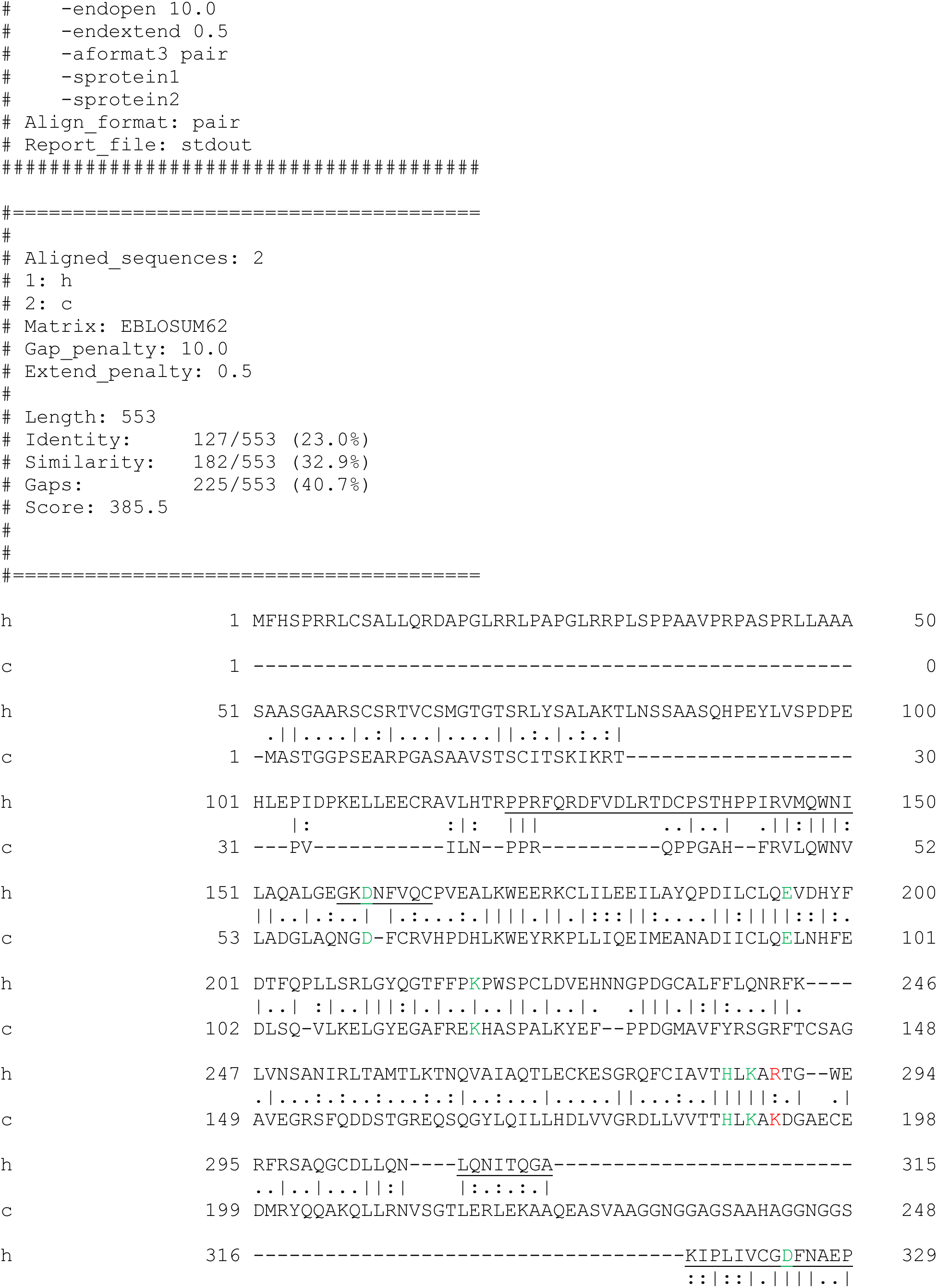
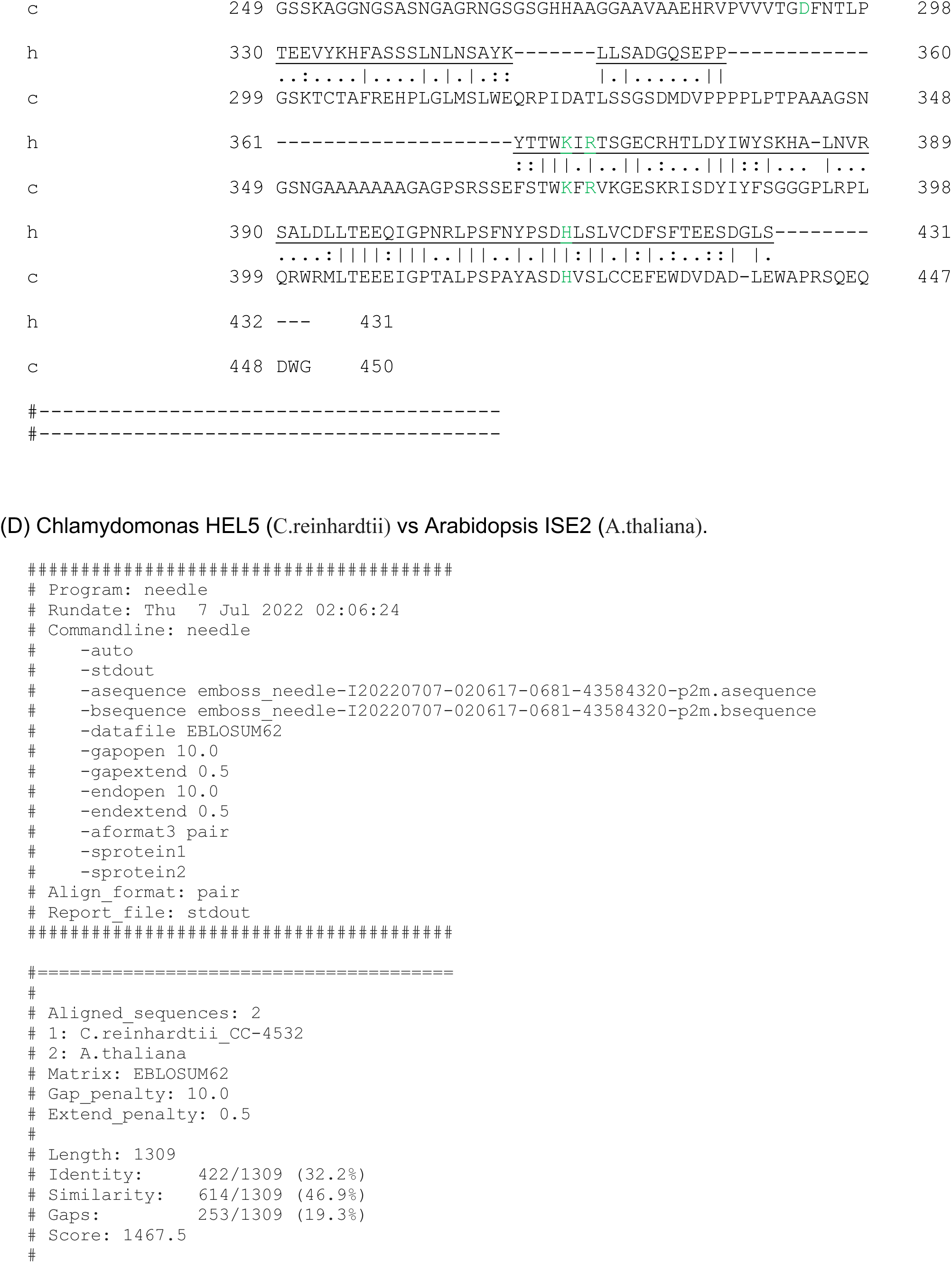
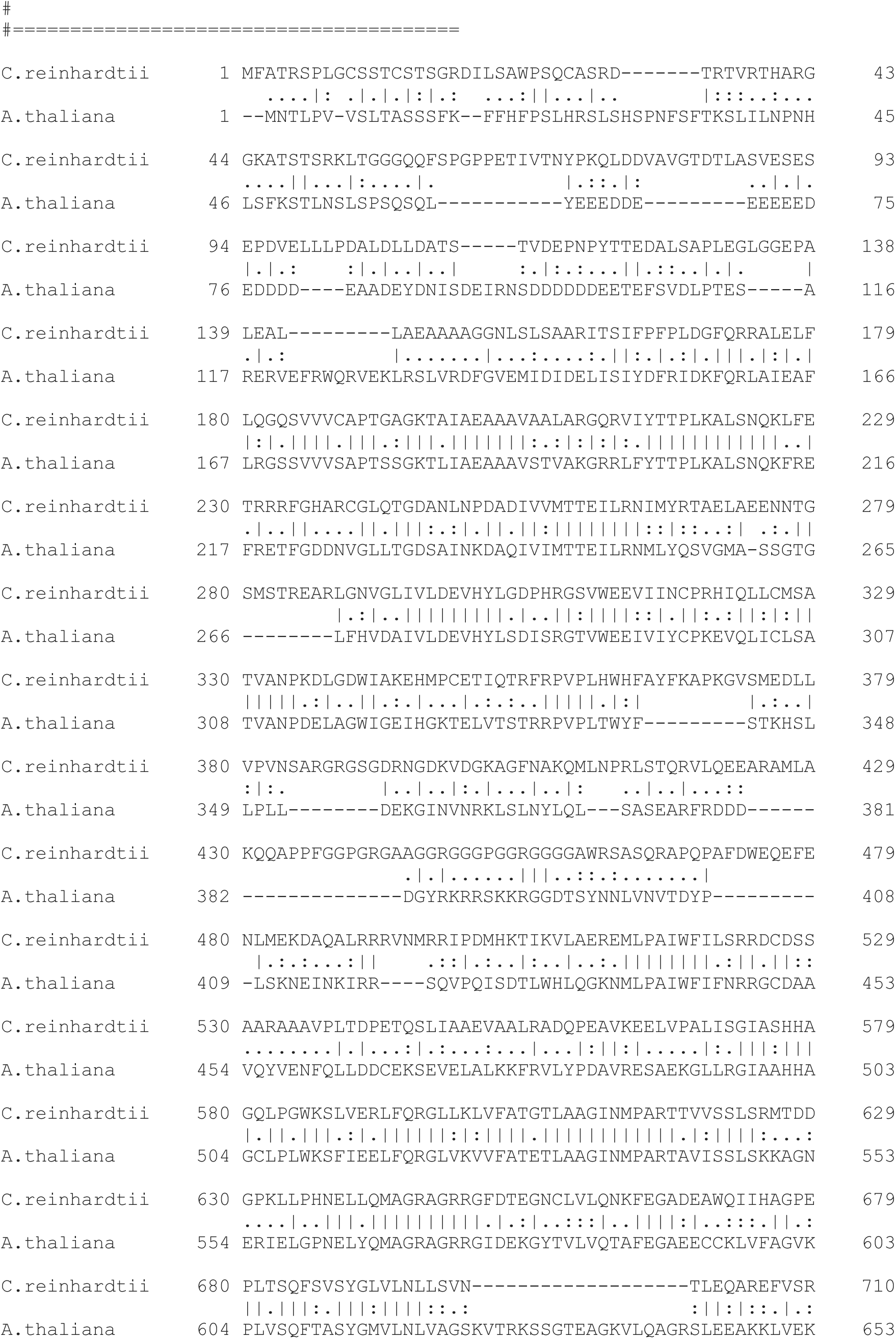
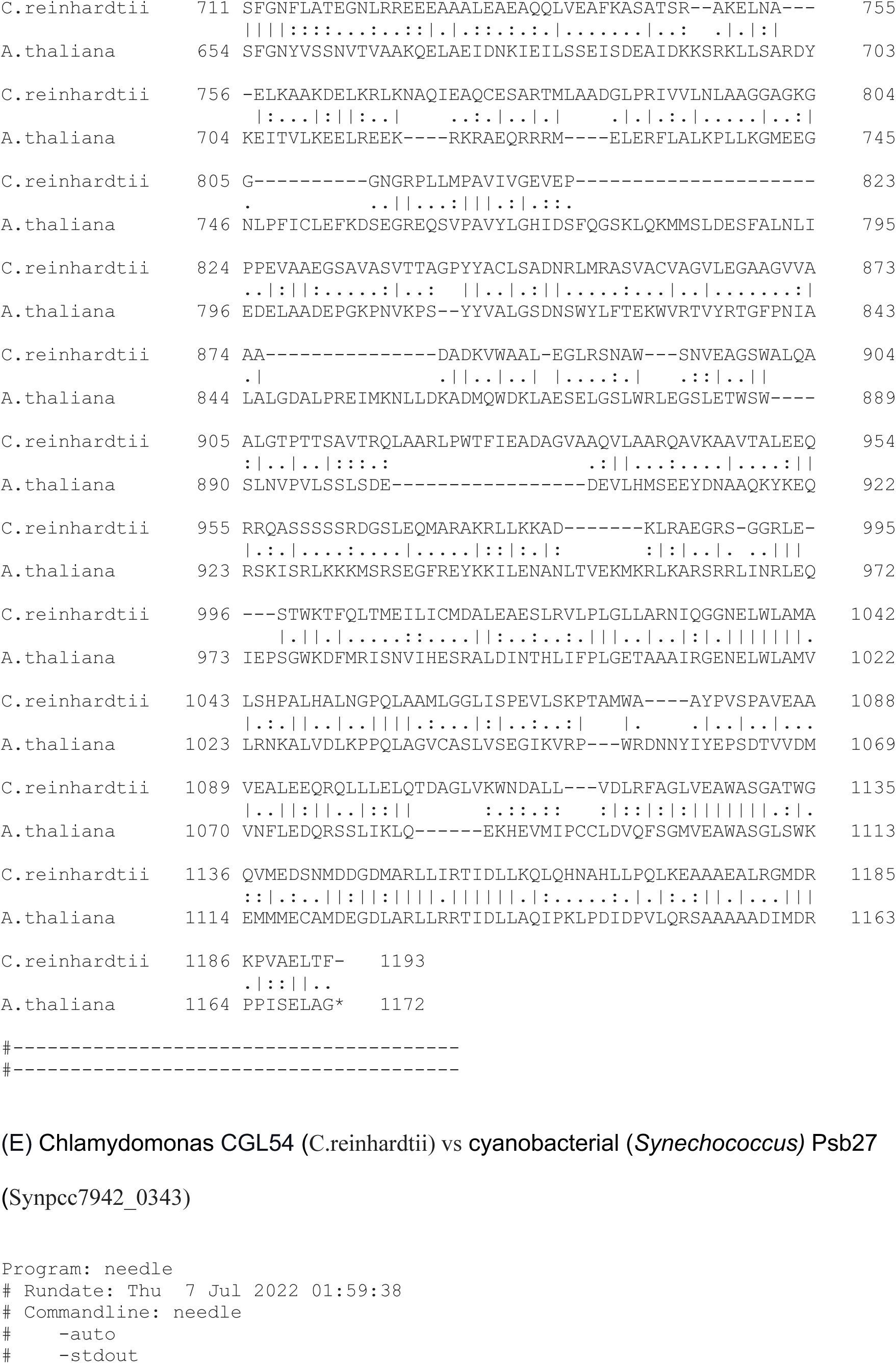
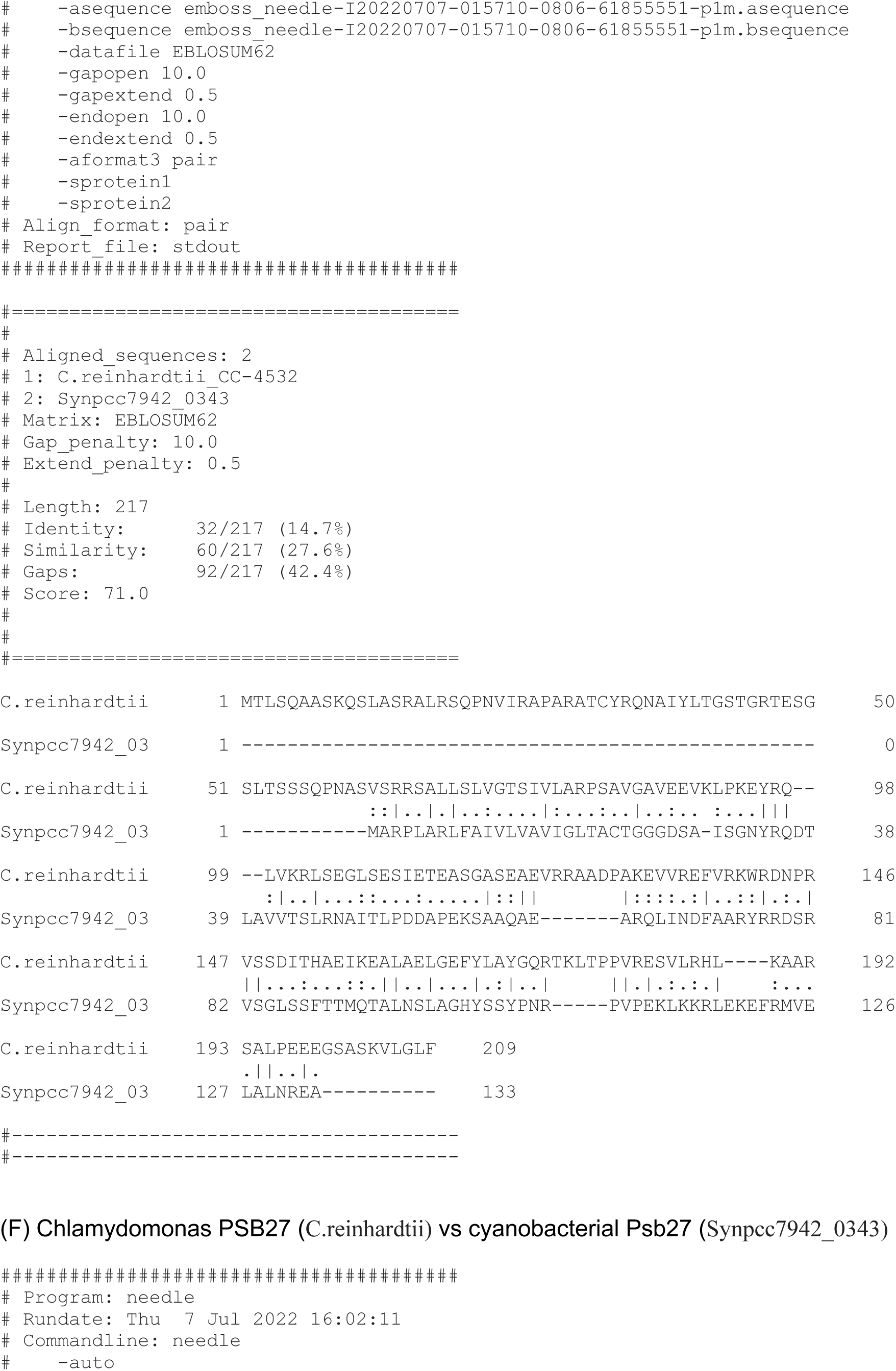
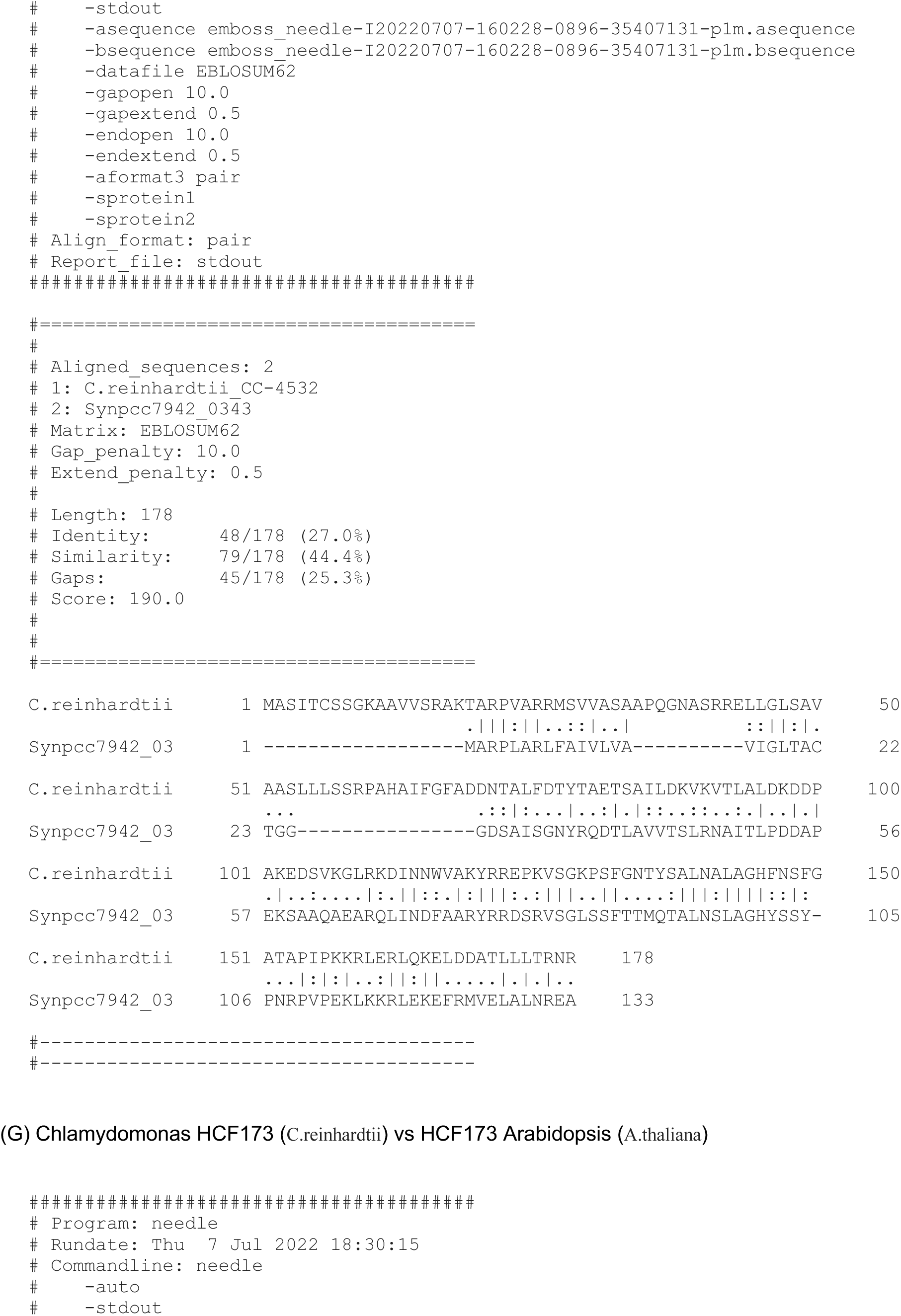
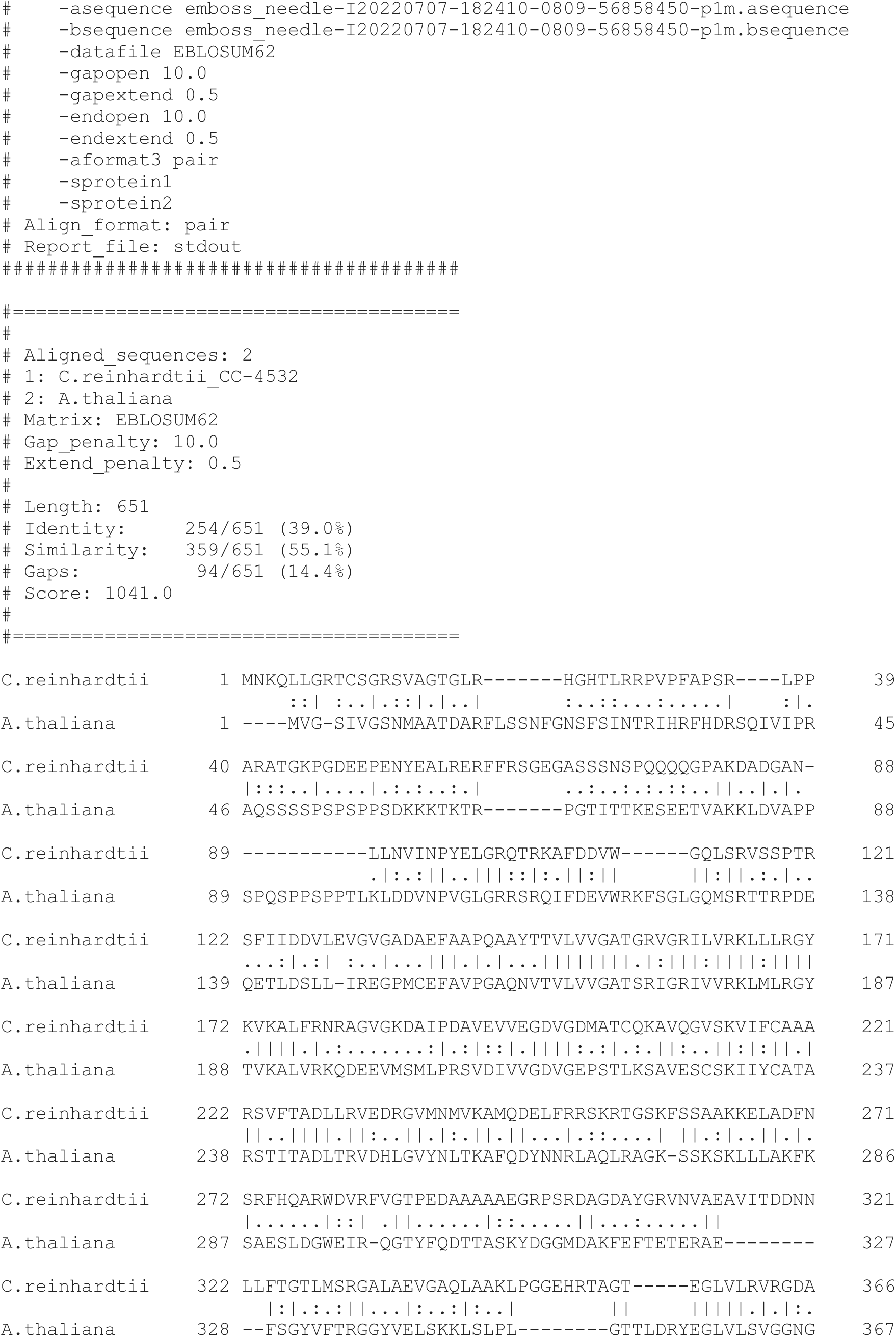
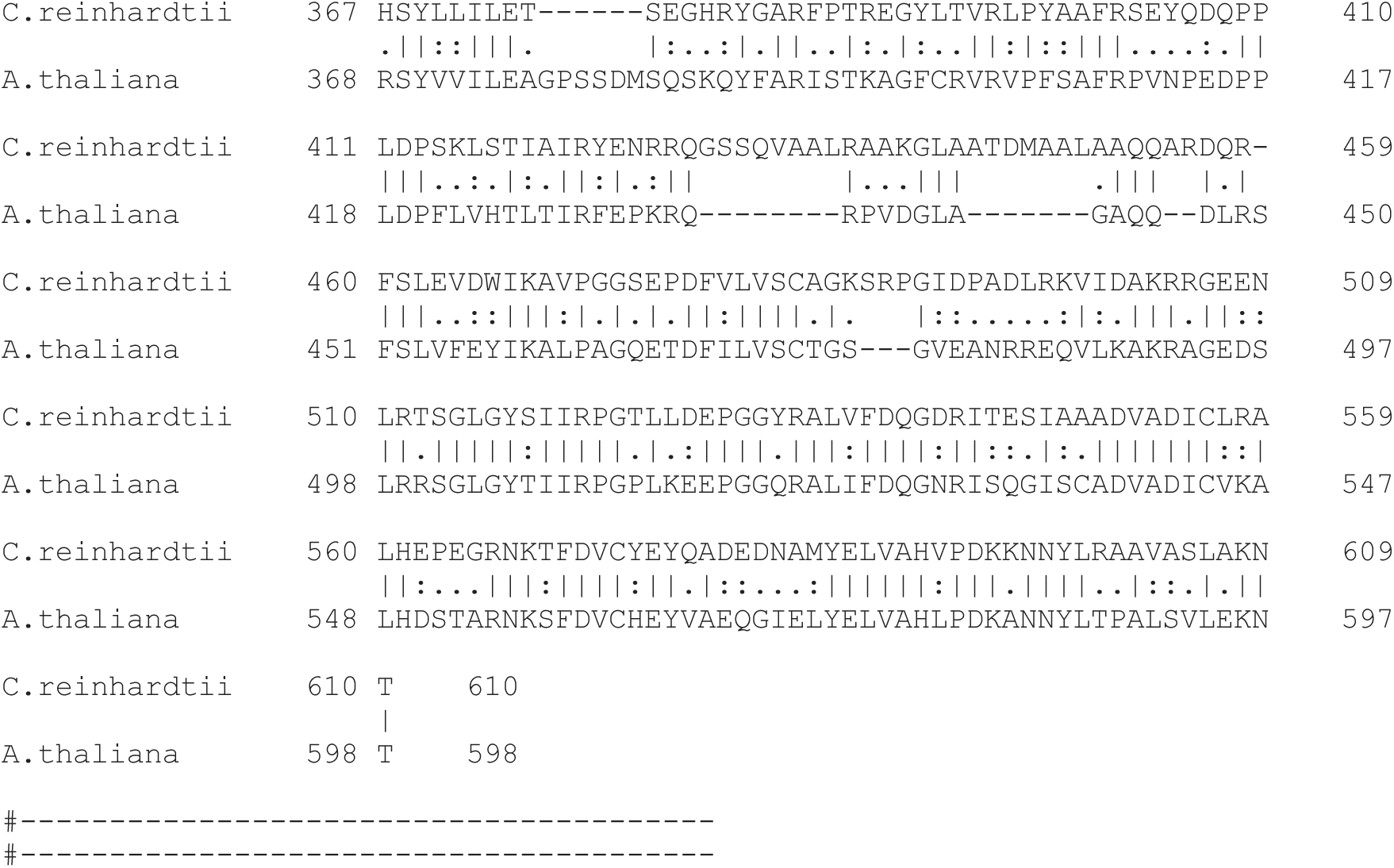
Sequence homology.

**Table S1 Phenotypic data of mutants and barcodes and mutants in the initial set of 1,781 mutants, related to Figure 1**

**Table S2 Hits from the pooled backcrossing experiments, related to Figure 1 and STAR Methods**

**Table S3 Protein localizations and suggested functions of other rescued genes, related to Figure 2**

**Table S4 Genes represented in the proteomic experiments, related to Figure 4**

**Table S5 - Proteomic data, related to Figure 4**

**Table S6 ROGEs affecting chloroplast genes, from the literature and from our data set, related to discussion**

**Table S7 Rescued mutants, the rescued gene, and the plasmids used for the rescue process, related to STAR Methods**

